# Deciphering complete archaic introgression sequences in modern human genomes

**DOI:** 10.64898/2026.07.23.740208

**Authors:** Mingyu Suo, Aoyue Bi, Quanyu Chen, Dan Yu, Linghan Jiang, Anguo Liu, Yiqing Yang, Hanyao Wang, Yanqing Sun, Lei Nie, Ruixue Chen, Qiangzhen Yang, Xiaotao Wang, Haishuai Wang, Yongyong Shi, Dan Zhang, Dongya Wu, Guojie Zhang

## Abstract

Genetic introgression from archaic hominins has profoundly reshaped the genetic diversity and adaptive potential of modern humans, yet the full catalog of introgressed sequences, particularly those residing in structurally complex regions has remained elusive. Here, we present ASMaid (ASseMbly-based archaic introgression detector), a Hidden Markov Model-based framework that leverages haplotype-resolved pangenome assemblies to identify archaic-derived sequences with unprecedented completeness. By integrating both single-nucleotide genotype and structural variation (SV) signals, ASMaid captures significantly more intact archaic segments than conventional reference-based approaches. Applying ASMaid to a global panel of 610 phased human genome assemblies, we show that non-African individuals carry approximately 79.8 Mbp of Neanderthal and 8.3 Mbp of Denisovan sequences, representing substantial increases over previous estimates, respectively. Notably, we detected several centromere-spanning archaic segments, including EAS-specific calls on chromosomes 5 and 7. Our assembly-based approach uncovered 1,701 archaic-derived SVs, revealing a previously overlooked layer of archaic functional legacy. High-frequency introgressed loci are enriched in pathways associated with metabolism, immunity, and nervous system (e.g. *CTNNA2* linked to early-onset schizophrenia risk), underscoring the fundamental role of introgression in modulating modern human traits. Notably, we identified dozens of loci potentially facilitating local adaptation, such as *PRDM16* involved in adipocyte differentiation and cold tolerance, and *CSGALNACT2* associated with chondroitin sulfate synthesis. Furthermore, our analysis delineates three distinct Denisovan introgression pulses in Eastern Eurasian genomes, in which the first two pulses are shared across East Eurasian and Oceanian populations, while the third remain primarily exclusive in East Asians. Reflecting these complex introgression events, 31 Denisovan-derived segments, including the *TBX15-WARS2* locus, are inferred to have been introduced via at least two events. This comprehensive map of archaic introgression provides a fundamental resource for understanding how ancient gene flow continuously shapes human phenotypic diversity and adaptation.

## Introduction

Archaic introgression from Neanderthals and Denisovans has profoundly shaped the genetic architecture of modern humans outside Africa, introducing variants that modulate local adaptation, disease susceptibility and phenotypic diversity across populations^1–6^. Neanderthal-derived alleles have been linked to immune signaling, skin pigmentation and metabolic homeostasis^7,8^, while Denisovan ancestry has facilitated high-altitude adaptation in Tibetans and influenced immune profiles across Asian populations^9,10^. Despite nearly two decades of research following the first archaic genome sequenced, a full catalog of introgressed genomic features remains elusive, obscuring the full extent to which ancient admixture affects human health and adaptation.

Current detection methods (e.g. IBDmix, Sprime, HMMmix, ArchaicSeeker), rely on single-nucleotide variations (SNVs) genotyping from short-read alignments in the “easy” regions of standard reference genomes, leading to systematic underestimation of archaic ancestry^11–14^. Despite successfully identifying thousands of introgressed segments, substantial inconsistencies persist across methods^15^, and critical gaps remain: archaic-derived structural variants (SVs) and genomic haplotypes within complex or repetitive regions, alongside their functional contribution to modern human traits have been poorly characterized. Recent advances in long-read sequencing have catalyzed human pangenome studies, with hundreds of nearly telomere-to-telomere assemblies of modern humans now available, which provide a fundamental basis for excavating the whole-genome full spectrum of archaic introgression, including these genomic “dark” regions^16–20^.

Here, we present ASMaid (Assembly-based archaic introgression detector), a Hidden Markov Model (HMM) that leverages phased human pangenome assemblies to identify complete introgression sequences with unprecedented resolution. By integrating SNVs and SV signals, ASMaid identifies intact haploid-resolved introgressed segments invisible to conventional approaches. Applying ASMaid to 610 phased genome assemblies from global populations, we discovered ∼79.8 Mbp of Neanderthal and ∼8.3 Mbp of Denisovan sequences per non-African individual, representing substantial increases over previous estimates. Notably, several centromeres in modern humans are inferred to be sourced from archaic introgression, including those on chromosomes 5 and 7 exclusive to East Asians. We identified over 1,700 archaic-derived SVs, and characterized their functional impacts on gene expression and disorder risk. Furthermore, dozens of population-stratified introgression genes possibly underlie human traits and adaptation, such as *PRDM16* related to adipocyte differentiation and cold tolerance, and *CSGALNACT2* associated with chondroitin sulfate synthesis. Specifically, by decomposing Denisovan components in Eurasian genomes, we proposed a three-pulse model of Denisovan introgression: two shared with Oceanians, along with a third in East Eurasia. Collectively, this comprehensive archaic introgression map fundamentally advances our understanding of how ancient gene flow shapes the genomic architecture of modern humans.

## Results

### ASMaid detects complete archaic introgression sequences from haplotype-resolved assemblies

To characterize the complete archaic introgression landscape, we developed ASMaid (Assembly-based archaic introgression detector), which directly analyzes haplotype-resolved genome assemblies to identify introgressed segments with full completeness (**Fig. 1a**; **Methods**). Unlike conventional approaches that infer introgression from diploid genotypes, ASMaid operates on phased assemblies, enabling detection of all variant types including structural changes that have remained largely invisible to previous methods. The core innovation of ASMaid is a HMM that simultaneously integrates SNV genotypes and copy number (CN) signals of SVs, providing dual and complementary evidence for introgression events. We utilized ten East African individuals from Human Genome Diversity Project (HGDP) as a background panel to distinguish true introgression from incomplete lineage sorting (ILS), and applied stringent filters to temporarily mask complex regions where CN inference may be confounded, including short tandem repeats (STRs/VNTRs), centromeres and ribosomal DNA (rDNA; **Methods**).

**Fig. 1.**
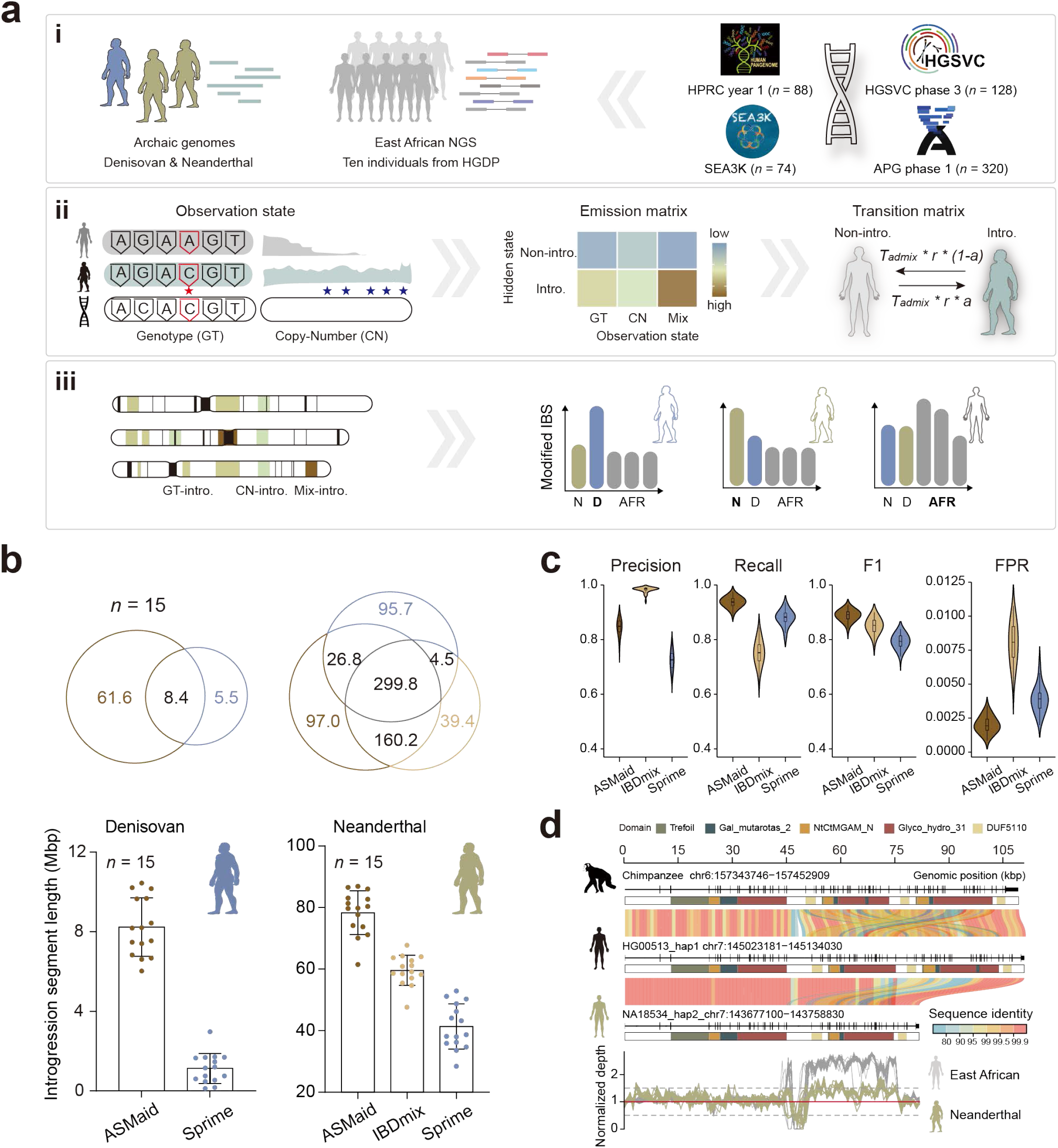
Identifying complete archaic introgression sequences using ASMaid. **a**, Schematic of the three-phase ASMaid workflow. (i) Data preparation: sequencing reads from four archaic genomes (three Neanderthals and one Denisovan) and ten East African individuals are mapped to each assembly from multiple pangenome projects (including APGp1, HPRCy1, HGSVC3, and SEA3K). (ii) HMM decoding: a multi-modal emission model integrates discrete SNV genotypes (GT) and copy number of structural variation signals (CN) into a single framework. (iii) Refinement: a modified IBS (Identity By State) is utilized to recalibrate ancestry and filter candidate introgressed segments. **b**, Performance comparison of ASMaid against established methods (IBDmix and Sprime) using 15 non-African individuals from the 1000 Genomes Project (1KGP). Top: Venn diagram illustrating the total genomic length of introgressed segments (Mbp) identified by each approach. Bottom: total identified introgression length per individual. **c**, Benchmarking on msprime-simulated data. Performance metrics, including precision, recall, F1-score, and false positive rate (FPR), are evaluated based on segment-level concordance (see Methods). **d**, Representative case of copy-number (CN)-based introgression at the *MGAM* locus. Top: Local synteny of *MGAM* homologs across chimpanzee (*Pan troglodytes*; GCA_028858775.2), a non-introgressed modern human assembly (HG00513_hap1) and an introgressed assembly (NA18534_hap2). Gene structures and predicted protein domains are annotated. Bottom: Normalized sequencing depth profiles for assemblies carrying the introgression signals, when short reads from East African and Neanderthal genomes are mapped. Dashed lines indicate the expected diploid normalized depth range (0.5-1.5).

We evaluated the performance of ASMaid by comparing with two widely used tools, IBDmix^13^ and Sprime^11^, using 15 non-African individuals from the 1000 Genomes Project (1KGP), whose phased genome assemblies have been generated by the Human Pangenome Reference Consortium year 1 (HPRCy1; **Supplementary Table 1**). ASMaid identified an average of 20.1-Mbp additional Neanderthal and 7.6-Mbp additional Denisovan sequences per individual, respectively, compared to the other two methods (**Fig. 1b**). These newly identified segments exhibited size distributions and match rates to archaic reference genomes comparable to those shared with existing tools (**Supplementary Fig. 1**). Notably, while ASMaid recovered the majority of archaic segments detected by IBDmix or Sprime, the uniquely identified segments were substantially enriched in repetitive and structurally complex genomic regions (**Supplementary Fig. 2**).

To validate the authenticity of these newly segments, we calculated the genetic distance between each segment across modern human populations and archaic individuals (**Supplementary Fig. 3**). We further compared the phylogenetic signals obtained from haplotype-resolved assemblies *versus* diploid genotypes in matched genomic regions. This analysis revealed that conventional diploid-based methods frequently encounter ambiguity when resolving heterozygous sites, producing conflicting signals that obscure introgression signatures, particularly in complex regions (**Supplementary Fig. 4a**). In contrast, ASMaid’s haplotype-resolved approach provides unambiguous signals for each chromosome copy, yielding clear phylogenetic placement even in genomic contexts where diploid inferences struggle (**Supplementary Fig. 4a**). The phylogenetic reconstruction provides support for the expected archaic introgression topology for the vast majority of the ASMaid-specific introgressed segments (**Supplementary Fig. 4b** & **4c**). Furthermore, simulation-based benchmarking demonstrated ASMaid’s robustness, with precision, recall, F1-score, and false positive rate (FPR), generally outperforming previous methods across multiple parameter settings (**Fig. 1c**; **Supplementary Fig. 5**), albeit with a trade-off between high recall and moderate precision for ASMaid.

ASMaid’s dual-signal approach integrating both genotyping information from SNVs and CN information from SVs enabled detection of introgression events fundamentally invisible to conventional methods. Across the tested 15 individuals, a total of 287 Neanderthal-derived and 80 Denisovan-derived segments exhibited substantial support from CN signals at >200 sites (**Supplementary Fig. 6**). To illustrate this advantage, we examined the *MGAM* gene, which encodes maltase-glucoamylase, a brush-border membrane enzyme critical for the final step of starch digestion^21^. Assemblies carrying Neanderthal ancestry at this locus revealed a structurally truncated MGAM architecture with a reduced set of functional domains (**Fig. 1d**; **Supplementary Table 2**). While the MGAM architecture in non-introgressed modern humans closely mirrors the ancestral state observed in great apes, comprising 16 to 17 domains, the introgressed Neanderthal-derived haplotype retains only 11 domains. Sequencing depth profile confirmed this copy number alteration, showing a specifically depleted pattern in Neanderthal genomes that contrasts with the doubled depth observed in East African individuals, providing independent evidence for the structural divergence of this archaic haplotype (**Fig. 1d**).

### Global landscape of complete archaic introgression sequences

We applied ASMaid to a panel of 610 phased human genome assemblies from multiple pangenome studies, including Asian Pan-Genome project phase 1 (APGp1), the Human Genome Structural Variation Consortium phase 3 (HGSVC3), the Human Pangenome Reference Consortium year 1 (HPRCy1) and SEA3K studies^16,18,20,22^ (**Supplementary Fig. 7**), generating the most complete and diverse archaic introgression maps at single-base resolution across global populations to date. In non-African (non-AFR) populations, ASMaid identified an average of 43.1-Mbp (10.8 to 54.1) Neanderthal and 4.4-Mbp (2.3 to 7.2) Denisovan sequences identified per haploid assembly (**Fig. 2a**; **Supplementary Table 3**). Compared to the reference-coordinates, the introgression sequences based on self assembly have slightly larger sizes, reflecting reference bias in genomic proportion statistics (**Supplementary Fig. 8**). When combining maternal and paternal haplotypes, non-AFR individuals carried ∼79.8 Mbp (20.3 to 95.0) of Neanderthal and ∼8.3 Mbp (4.8 to 12.3) of Denisovan ancestry sequences (**Supplementary Fig. 9**; **Supplementary Table 4**). Consistent with previous findings^7,23^, the East Eurasians (East Asian, EAS: 82.3 and 8.7 Mbp for Neanderthal and Denisovan, respectively; Southeast Asian, SEA: 79.7 and 7.6 Mbp) have more archaic sequences than Western Eurasians (European, EUR: 73.0 and 6.9 Mbp). An increase of ∼21.1-Mbp Neanderthal and ∼6.3-Mbp Denisovan introgression segments over those detected by IBDmix and Sprime were yielded across APGp1 individuals, respectively, resulting in additional calls of at least 100 Mbp of Denisovan-derived and 156 Mbp of Neanderthal-derived sequences across the EAS superpopulation, despite a significant correlation in segment lengths between ASMaid and other tools (**Supplementary Fig. 10**; **Supplementary Table 4**). Totally, across all 610 assemblies, we identified 7,415 non-redundant Neanderthal-derived genomic chunks totalling 1,227.5 Mbp and 4,616 Denisovan-derived chunks totalling 257.9 Mbp in the coordinates of T2T-CHM13. Of these, 236 Neanderthal chunks (249.6 Mbp) and 20 Denisovan chunks (4.2 Mbp) reached high frequencies (HF) over 40% across all non-AFR genomes (**Fig. 2b**; **Supplementary Fig. 11**). The high-resolution haplotype-based detection enabled identification of rare introgression events, with 21.5% of Neanderthal chunks (1,595, totalling 66.4 Mbp) and 32.6% of Denisovan chunks (1,505, 51.4 Mbp) appeared as singleton among non-AFR assemblies, highlighting ASMaid’s sensitivity for detecting low-frequency archaic haplotypes that contribute to individual genetic diversity (**Fig. 2b**; **Supplementary Tables 5** and **6**).

**Figure 2.**
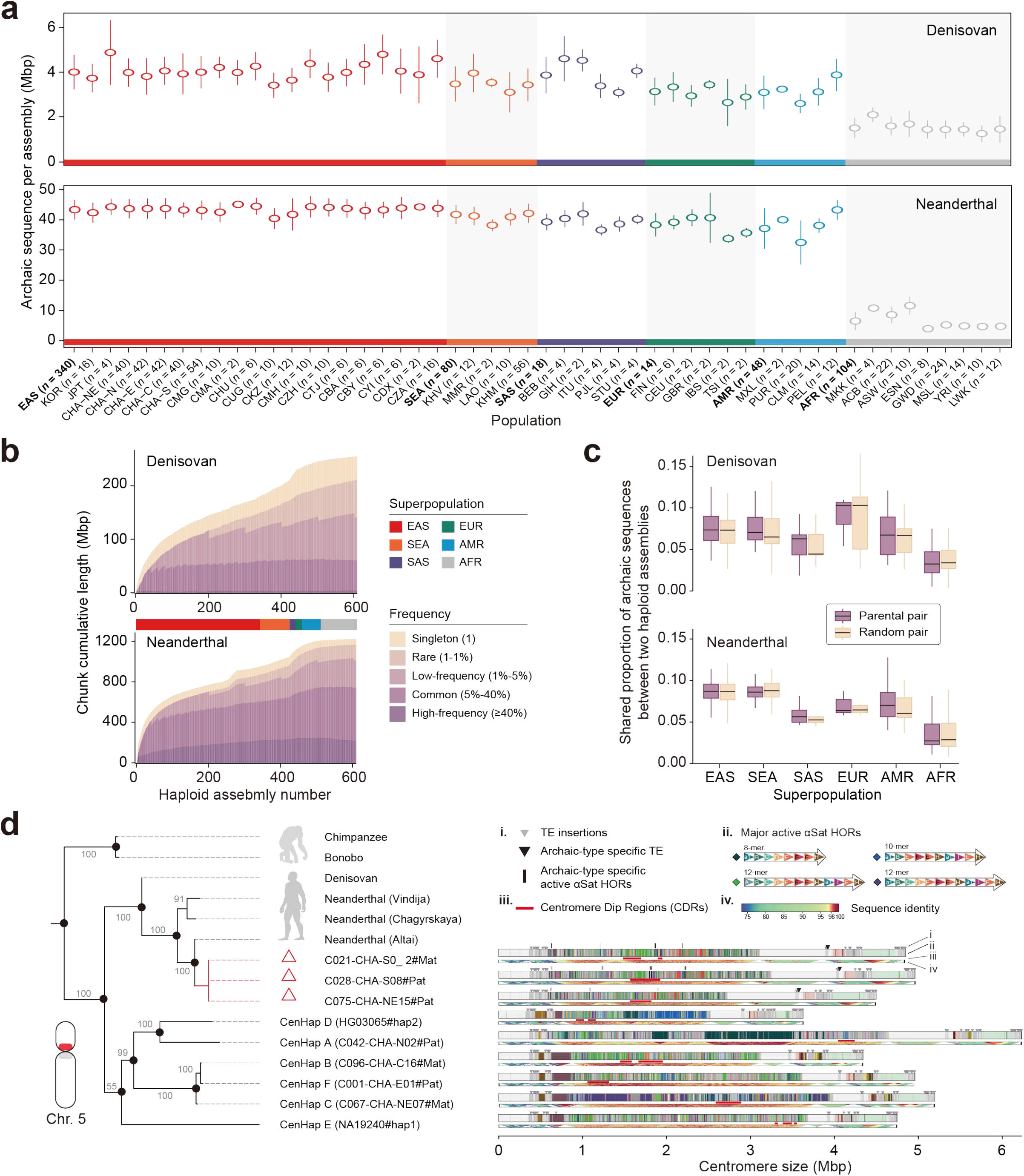
Global profiling of archaic introgression segments across human haplotype-resolved genome assemblies. **a**, Length of haploid-resolved archaic introgression segments from Denisovan (top) and Neanderthal hominin (bottom) across human populations as identified by ASMaid. Only autosomes are included. **b**, Cumulative curves of archaic segment lengths in modern human genomes. The non-redundant archaic sequences for Denisovan (top) and Neanderthal (bottom) are added sequentially by superpopulation in the following order: EAS (East Asian), SEA (Southeast Asian), SAS (South Asian), EUR (European), AMR (American) and AFR (African). The frequencies are calculated based on total allele counts with the current dataset. **c**, Heterogeneity in archaic segments between modern human genomes. **d**, Characterization of a Neanderthal-derived centromere-spanning introgression on chromosome 5. A maximum-likelihood phylogeny among main centromere haplotypes of chromosome 5 is built using pericentromeric flanking sequences, with 1000 replicates. Structural compositions of major centromere haplotypes are illustrated, highlighting key features of the archaic-type centromere. (i), lineage-specific transposable element (TE) insertions and higher-order repeat (HOR) structures; (ii), organization of major active HOR variants; (iii), centromere dip regions identified by methylation profiling; and (iv), local sequence similarity indicative of homogenization.

Introgressed segments exhibited substantial variation in length and frequency across individuals, reflecting differential post-introgression recombination and selection pressures. We identified 196 ultra-long archaic segments longer than one mega-base pair (Mbp) from 38 chunks, with the longest segment approaching 2 Mbp (**Supplementary Table 7**). For example, 43 haploid assemblies have a ∼1.4-Mbp Neanderthal-derived segment at 4q28.1-4q28.3 (N2007) and seven have a ∼1.1-Mbp Denisovan segment at 1p34.1-1p33 (D100; **Supplementary Fig. 12a**). The genomic locations of these ultra-long segments exhibited significantly lower recombination rates than those short archaic fragments (**Supplementary Fig. 12b**). Additionally, we noticed striking heterogeneity of archaic ancestry between haplotypes (**Fig. 2c**). Less than 10% were shared between the maternal and paternal haploid within a diploid individual, comparable to random haplotype pairs. The pronounced haplotype heterogeneity was in line with the relatively openness of the archaic-segment growth curve, suggesting the significance of haploid-resolved identification for comprehensively cataloging archaic introgression.

While recent studies have documented low levels of archaic ancestry in African populations, attributed to back-migration from Eurasia^13^, ghost introgression from unknown archaic humans^24,25^ or human-to-archic introgression^26,27^, the complete extent and characteristics of this ancestry have remained poorly characterized. Our analysis detected an average of 6.9-Mbp Neanderthal and 1.8-Mbp Denisovan ancestry in AFR haploid assemblies, respectively, similar as previous estimates in Africans when considered as the diploid level^13,23^ (**Fig. 2a**; **Supplementary Tables 3** and **4**). This result further suggested the evaluated archaic sequences in non-AFR human genomes are not systematic artifacts. For Neanderthal introgression, 98.4% of the 298.8-Mbp archaic segments in AFR genomes overlapped with those in non-AFR superpopulations, with the highest sharing with EUR, which supported the back-migration hypothesis (**Supplementary Fig. 13**; **Supplementary Table 8**). The proportion of Neanderthal ancestry showed clear stratification within African ancestry, where Africans from America (ACB and ASW) display the most abundant, followed by West African coastal populations (GWD and MSL; **Fig. 2a**). Similarly, for Denisovan introgression, we observed ∼66.8 Mbp of archaic sequences in AFR genomes, in which 95.7% are shared with non-AFR genomes (**Supplementary Fig. 13**).

### Centromere-spanning archaic introgression segments

Notably, we identified 40 candidate archaic-derived introgression segments that span entire centromeric regions across six chromosomes (**Extended Data Fig. 1**; **Methods**). These events occurred at low frequencies on chromosomes 12 (*n* = 25), 7 (*n* = 6), 5 (*n* = 3), 11 (*n* = 3), 6 (*n* = 2), and singleton on chromosome 10, consistent with expected signature of neutral evolution on centromeric sequences. Phylogenetic analysis of flanking pericentromeric regions confirmed elevated sequence similarity between modern human carriers and archaic hominin genomes (**Supplementary Fig. 14; Supplementary Table 9**). For the five acrocentric chromosomes, a complete assessment of centromere introgression was hindered by poor assembly of the short (p) arms, which consist primarily of ribosomal DNA arrays and remain challenging to assemble even with long-read technologies^19^. However, we detected introgression signals adjacent to the centromeres on the long (q) arms of these chromosomes in multiple individuals, indicating the possibility of centromere introgression on these five chromosomes as well (**Supplementary Fig. 15**). While centromere introgression has been documented in yeast^28^, rice^29^ and human^30^, it should be noted that on certain chromosomes (6, 11 and 12), we observed similar signals in African individuals (**Extended Data Fig. 1**). This pattern implies that alternative evolutionary processes merit consideration, including genetic drift in recombination-suppressed centromeric regions maintaining ancient haplotypes, or incomplete-lineage sorting (ILS) where ancestral polymorphisms predating the modern human-archaic split were randomly retained in different lineages, or ancient introgression events in African genomes from deeply divergent archaic populations^24^. Therefore, these cases remain ambiguous to be interpreted as definitive introgression.

Notably, chromosomes 5 and 7 exhibited centromere introgression exclusively within the EAS superpopulation, spanning 6.4 and 10.6 Mbp on average, respectively, with robust support from continuous introgression signals across both flanking pericentromeres (**Extended Data Fig. 1**). Taking chromosome 5 as an example (**Fig. 2d**), the introgressed sequences in both pericentromeres showed complete linkage in all carrier assemblies, suggesting a single, relatively recent introgression event (**Supplementary Fig. 16**). This perfect linkage across the centromere, a region where recombination is overall suppressed, provides strong evidence against ILS. Under ILS, the p-arm and q-arm haplotypes would have evolved independently over hundreds of thousands of years prior to the modern human-Neanderthal split, making their coordinated presence in the same individuals extremely unlikely by chance. Moreover, ILS would predict stochastic retention of ancestral haplotypes across non-AFR populations with no geographic bias, whereas these centromere haplotypes are restricted to EAS individuals, consistent with a geographically localized admixture event and the known pattern of elevated archaic introgression in EAS. By contrast, introgression would transfer both pericentromeric regions simultaneously as a single unit, while naturally producing a geographically localized signal (**Supplementary Fig. 16**).

The archaic origin of these centromere haplotypes is further corroborated by their distinctive sequential architecture. The three unrelated assemblies from APGp1 exhibited a special centromere haplotype (CenHap) distinguished from the common CenHaps in other modern human genomes. Functional centromeres for the three chromosomes were characterized within highly homogenized intervals as centromere dip regions (CDRs) with decreased methylation (**Fig. 2d**). The Neanderthal-type CenHap harbored unique transposable element (TE) insertions and High-Order-Repeat (HOR) variants, such as C25H641 (8-mer), C25H927 (22-mer), C25H1100 (14-mer) and C25H272 (8-mer; **Fig. 2d**)^31^. The convergent presence of this precise molecular composition in three unrelated EAS individuals is therefore highly unlikely to reflect either ILS or parallel evolution, and instead provides strong evidence for direct transfer from the Neanderthal lineage. Collectively, the discovery of centromere-spanning introgression represents a previously unknown circumstance of large-scale genomic exchange between archaic and modern humans, provoking new thinking about the genomic scale of archaic introgression and evolutionary dynamics of centromeric sequences in humans.

### Archaic structural variants shape modern human genomes

While previous studies of archaic introgression have primarily focused on SNVs, the functional impact of larger SVs, including insertions, deletions, and inversions spanning hundreds to thousands of base pairs, remains largely unexplored, despite growing evidence that SVs are often subject to stronger selective pressures and play crucial roles in human local adaptation^1,32^. Here, we identified a repertoire of 1,701 putatively archaic-derived SVs (pAID-SVs) across 610 genome assemblies, comprising 1,386 Neanderthal-associated and 315 Denisovan-associated variants (allelic frequency ≥10 haploids; **Fig. 3a**; **Supplementary Table 10**). These pAID-SVs were identified by comparing allele frequencies (AFs) between introgressed and non-introgressed genomes (*P* value < 5×10^-8^, Fisher’s exact test), and further validated through mapping profiles of coverage and clipping information from archaic hominin genomes. Notably, 79.4% of Neanderthal and 77.1% of Denisovan-associated SVs were validated with high confidence (HC) present in at least one archaic genome (**Fig. 3a**). Their size distribution and repeat composition mirror those SVs originated among modern humans, with a predominant composition of SINE/Alu elements (**Supplementary Fig. 17**).

**Fig. 3.**
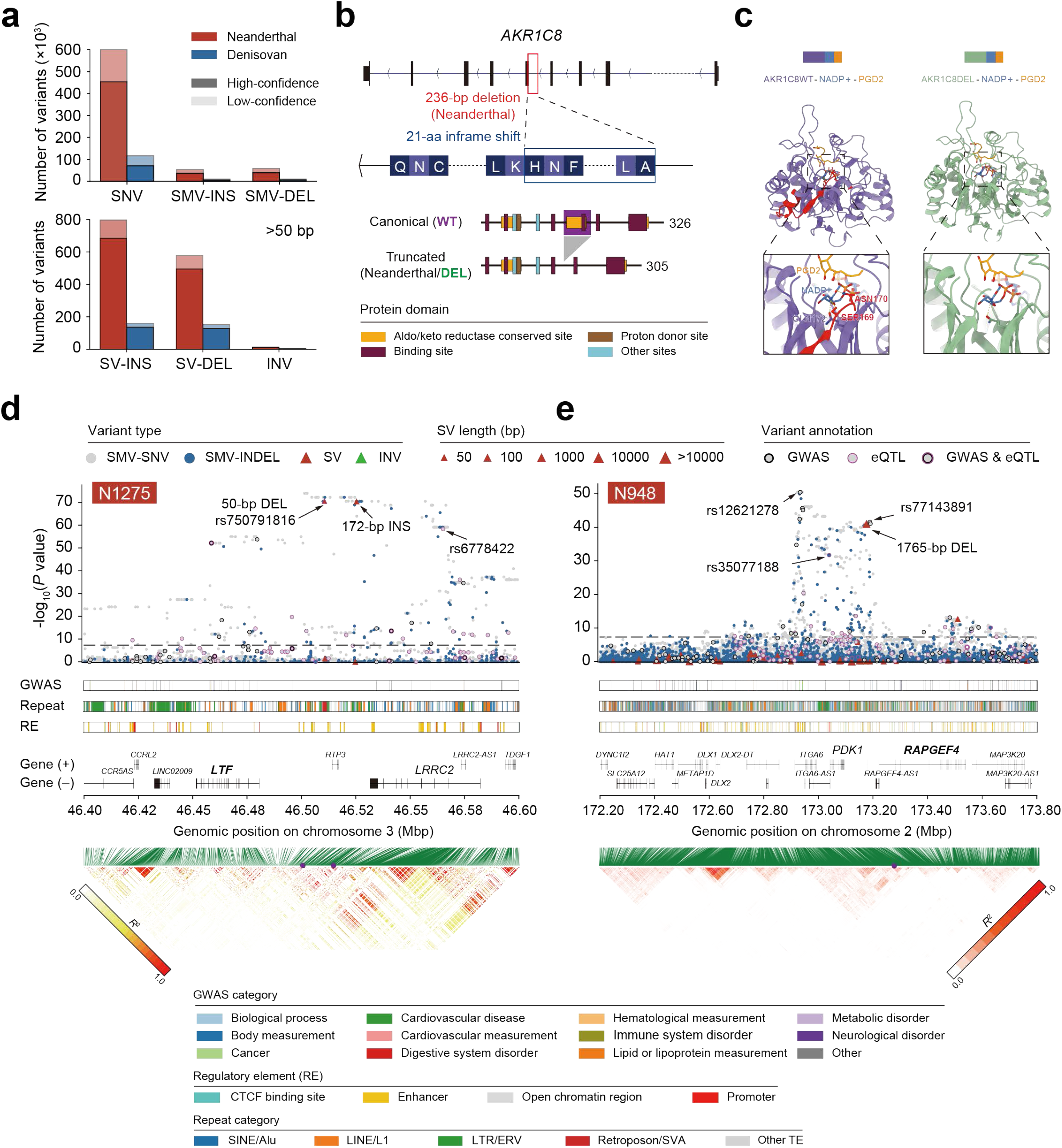
Structural variants derived from archaic introgression in modern human genomes. **a**, Quantification of putative archaic introgression-derived variants, stratified by variant type (top: small variants; bottom: structural variants) and confidence level. **b**, Characterization of a specific 236-bp Neanderthal-derived deletion within the *AKR1C8* gene. This deletion introduces a 21-amino acid in-frame shift, resulting in the loss of protein domains as annotated by Uniprot and PROSITE. **c**, Comparative structural modeling of wild-type (WT; canonical) and truncated (DEL; deletion) AKR1C8 protein structures. Left: WT structure, with the deleted region highlighted in red; right: modeled DEL structure. Both structures are in the complex with NADP^+^ and PGD2. Zoomed-in views with hydrogen bonds indicated by yellow dashed lines illustrate that the DEL protein specifically loses the hydrogen bonds normally mediated by Ser169 and Asn170. **d** and **e**, Genomic views of the loci 3p21.31 (N1275) and 2q31.1 (N948). Top: introgression significance signals (Fisher’s exact test) for each variant, color-coded by variant type and functional annotation, including GWAS and eQTL signals. Middle: integrated tracks for GWAS signals (colored by trait category), repetitive elements (annotated by Repeatmasker and colored by major repeat class), regulatory elements (Ensembl 113), and gene annotations (Refseq). Bottom: genetic linkage heatmaps, with focal SVs indicated by purple dots. SMV, small variant; INV, inversion.

Among these pAID-SVs, 747 Neanderthal-derived and 147 Denisovan-derived SVs overlap with 682 and 122 genes, respectively (**Supplementary Fig. 17; Supplementary Table 11**), with 19 directly disrupting protein-coding sequences. Notable examples include the complete loss of *CDK11A*, a gene involved in cell cycle regulation^1^, via a ∼53-kbp Neanderthal deletion, and a 31-kbp deletion of the pharmacogene *CYP2A6*, critical for drug metabolism^33^ (**Supplementary Fig. 18**). Furthermore, a particularly striking example might demonstrate how SVs can fundamentally alter protein function that a 236-bp Neanderthal-derived deletion (14.5% frequency in non-AFR genomes) partially removes the fifth exon of *AKR1C8*, which encodes an aldo-keto reductase involved in prostaglandin biosynthesis^34,35^ (**Supplementary Fig. 19**). Crucially, this variant causes an in-frame loss of 21 amino acids, eliminating the NADP+ binding site (Q5T2L2) and the conserved catalytic domain of aldo-keto reductase family (IPR018170; **Fig. 3b** & **3c**). Such a structural alteration results in the complete removal of an active domain, likely exerting biological impacts on prostaglandin metabolism.

Beyond coding disruptions, we identified 77 Neanderthal-derived and 19 Denisovan-derived SVs overlapping regulatory elements (REs), suggesting the potential for altering gene expression (**Supplementary Fig. 17**; **Supplementary Table 10**). We associated these archaic SVs with expression quantitative trait loci (eQTL) and genome-wide association study (GWAS) signals. One of the most medically relevant examples involves the 3p21.31 locus (*SLC6A20*-*CCR3*), previously identified as the major genetic risk factor for severe COVID-19^8,36^. We identified two previously unreported archaic SVs within this region in strong linkage (*R*^2^ = 0.96): a 50-bp deletion (rs750791816) and a 172-bp insertion, both exhibiting elevated frequencies in South Asian (SAS) and EUR superpopulations and signatures of recent positive selection (**Fig. 3d**; **Supplementary Fig. 20a** & **20b**). Functionally, the 50-bp deletion is associated with the up-regulation of *LTF*, an antiviral gene active against SARS-CoV-2 and HIV^37^, in skeletal muscle (GTEx) and blood (**Supplementary Fig. 20c**). Conversely, both SVs are highly linked to a pAID-SNV (rs6778422; *R*^2^ = 0.79 and 0.83, respectively) that downregulates gene expression of tumor suppressor *LRRC2* in pituitary (GTEx) and blood^38^ (**Supplementary Fig. 20d**).

Another case involves cardiovascular disease risk at the locus 15q25.1 (N6137), where a Neanderthal-derived 321-bp SINE/Alu insertion, frequent in EAS and SEA, sits upstream of *ADAMTS7* within a cluster of enhancers. This gene is genetically associated with the coronary artery disease risk^39^. This archaic insertion appears to alter *ADAMTS7* expression, potentially modulating cardiovascular disease susceptibility, and showed the recent positive selection signal (**Supplementary Fig. 21**). Similarly, at 2q31.1, a 1,765-bp Neanderthal deletion, prevalent in EAS and SEA superpopulations (**Fig. 3e**; **Supplementary Fig. 22a**), lies upstream of *RAPGEF4*, a gene critical for neural signaling and implicated in autism spectrum disorders^40^. This deletion shows extremely strong linkage with an adjacent pAID-SNV (rs77143891; *R*^2^ = 0.97) associated with attention deficit hyperactivity disorder (ADHD), implying a role for archaic SVs in contributing to neurological phenotype variation. Another linked pAID-InDel (rs35077188; *R*² = 0.55) acts as a *cis*-eQTL, down-regulating *PDK1*, a key metabolic enzyme in hypoxia response across multiple tissues (**Supplementary Fig. 22b**). Additionally, we identified a linked pAID-SNV (rs12621278; *R*^2^ = 0.46) located within an enhancer of *ITGA6,* a gene encoding an integrin essential for sperm-egg fusion and implicated in tumorigenesis (**Fig. 3e**), shows protective effects against prostate carcinoma^41^.

Regarding Denisovan introgression, within the *EPAS1*-*TMEM247* region (2p21, D456) responsible for high-altitude adaptation and hypoxia response in the Tibetan population, a 3.4-kbp deletion (dbsv66240) was identified upstream of *TMEM247*, despite its absence in the Altai Denisovan reference. This variant has been proposed to regulate the expression of local genes by functioning as a super-enhancer^42,43^. Similarly, locus *TBX15-WARS2* (1p12, D201) contains a 316-bp Alu deletion, highly prevalent in EAS and AMR superpopulations, which is strongly linked to a *cis*-eQTL variant (rs10923738; *R*^2^ = 0.95) and three GWAS signals (*R*² > 0.8) associated with body measurement and eye morphology^44,45^ (**Supplementary Fig. 23**). Additionally, at 12p12.2 (D3376), we discovered a 319-bp Denisovan-derived insertion, frequent in SAS and EUR, that lies downstream of *SLCO1C1*, a gene encoding a thyroid hormone transporter critical for regulating thyroid hormone availability in the central nervous system^46^. Although Denisovan reference displayed non-introgression allele via read mapping, we inferred this allele was probably from another Denisovan lineage, similar as the 3.4-kbp deletion at the *EPAS1* locus (**Supplementary Fig. 24a**). This insertion is linked with a *cis*-eQTL variant (rs1515764; *R*^2^ = 0.88) that decreases *SLCO1C1* expression (**Supplementary Fig. 24b**), potentially affecting cognitive function and metabolic regulation through altered thyroid hormone signaling in the brain. Collectively, these findings demonstrate that haplotype-resolved assemblies reveal a previously hidden layer of archaic introgression through thousands of SVs with complex regulatory effects across immunity, metabolism, and neural development, fundamentally expanding both the catalog of introgressed sequences and our understanding of their functional impacts on modern human adaptation.

### Functional and clinical implication of archaic introgression

Beyond individual loci, we investigated whether archaic introgression systematically influenced specific biological pathways. We identified 1,905 genes harboring high-frequency introgression (>40%) from Neanderthal (1,803) or Denisovan (102) in modern humans (**Methods**; **Supplementary Table 12**). Both Neanderthal and Denisovan introgression showed enrichment for defense response, including viral defense from Neanderthals (GO:0051607; BH-adjusted *P* = 6.01×10^−5^) and defense against other organisms from Denisovans (GO:0098542; BH-adjusted *P* = 4.47×10^−5^; **Supplementary Fig. 25**). Moreover, both showed particularly strong enrichment for neurological functions (**Supplementary Table 13**). Genes with Neanderthal introgression were significantly enriched for schizophrenia associations (BH-adjusted *P* = 4.18×10^−8^), while Denisovan genes were enriched for response to antipsychotic therapy (BH-adjusted *P* = 5.12×10^−3^). Furthermore, we observed population differences in functional enrichment of HF genes (**Supplementary Tables 14** and **15**). Denisovan introgressed genes showed population-specific enrichment for drug metabolism (KEGG:hsa00982, BH-adjusted *P* = 0.0310) in EUR, and lipid metabolism (KEGG:hsa00062, BH-adjusted *P* = 0.0142) in SEA, while they linked to developmental processes enriched in EAS (e.g. GO:0043588, BH-adjusted *P* = 7.73×10^-4^), SAS (e.g. GO:0070372, BH-adjusted *P* = 2.36×10^-5^), and SEA (e.g. GO:0070372, BH-adjusted *P* = 0.0110). As compared, Neanderthal introgression genes in developmental pathways were significantly enriched only in SEA (e.g. GO:0008544, BH-adjusted *P* = 9.41×10^-10^). These patterns suggest that archaic introgression might provide adaptive advantages tailored to specific genetic backgrounds or environmental contexts, fueling the phenotypic diversity of modern human populations.

To quantify clinical impact of these legacies, we cross-referenced archaic variants with the GWAS catalog. We identified 7,323 Neanderthal and 1,613 Denisovan pAID-SMVs, alongside 25 Neanderthal and five Denisovan pAID-SVs, that overlap with GWAS signals (**Supplementary Tables 16** and **17**). Similarly, 13,527 Neanderthal and 3,176 Denisovan variants function as significant expression quantitative trait loci (eQTLs), collectively associated with the expression of thousands of genes (**Supplementary Table 18**). Additionally, 62 Neanderthal and 11 Denisovan pAID-SVs were identified as being associated with eQTLs (**Supplementary Table 19**). These regulatory variants span the functional categories identified in enrichment analysis linked to disease susceptibility and gene regulation. For example, a 2.8-kbp Neanderthal deletion that fully deleted the *HLA-U* pseudogene located upstream of *HLA-A*, overlapping with GWAS locus rs2524005, previously associated with bipolar disorder, schizophrenia, and cancer^47,48^. Additionally, a Neanderthal-derived 81-bp deletion, rs1566617917, within a SINE/MIR element acts as a *cis*-eQTL for multiple immunity-related genes.

The enrichment of archaic introgression in neurologically related pathways prompted us to examine whether specific introgressed variants contribute to psychiatric disease risk. We associated archaic-derived variants with susceptibility loci for early-onset schizophrenia (EOSCZ), identified using whole-genome data from 1,251 cases and 1,123 controls collected by the APG associated program^49^. This analysis identified 114 Neanderthal-derived variants in *CTNNA2* at 2p12 that were significantly associated with EOSCZ risk (FDR < 0.05; **Extended Data Fig. 2**; **Supplementary Table 20**). *CTNNA2* encodes a neural cell adhesion protein critical for neuron migration and synaptic formation, and has been independently identified as a susceptibility gene for schizophrenia and bipolar disorder^50–52^. These introgressed segments lie upstream regulatory region of *CTNNA2*, harboring one pAID-SNV (rs79285146) significantly associated with EOSCZ (introgression Fisher’s exact test *P* = 1.96×10^-45^; EOSCZ FDR *P* = 0.0223) located in an enhancer (1,449 bp from the transcription start site). This locus also harbors additional variants associated with cancer (rs1864550) and nervous system functions (rs1159598 and rs7570469), suggesting pleiotropic effects (**Extended Data Fig. 2**). While these associations require functional validation, they illustrate how archaic-derived regulatory variants may have influenced the neuropsychiatric disease risk in modern humans.

### Population-stratified archaic introgression and selection

To understand how archaic introgression contributed to population-specific adaptation, we characterized the landscape of introgression across superpopulations (**Supplementary Tables 5** and **6**). Among 223 high-frequency (HF, >40%) Neanderthal-derived segments in EAS, 179 (166.7 Mbp), 77 (85.5 Mbp), and 71 (77.7 Mbp) are shared with SEA, SAS and EUR, respectively (**Supplementary Fig. 26a**). Within this reservoir, 738 introgression sites exhibited hallmarks of positive selection in EAS (**Supplementary Fig. 27**). A core set of 43 chunks is commonly maintained across all five superpopulations, including N7288 (20q13.33) associated with neurodevelopment (*KCNQ2*) and pancreas development (*PPDPF*)^7^, N7011 (19q13.11) related to metabolic homeostasis^53,54^ (*RHPN2*, *LRP3* and *SLC7A10*), N4299 (9q33.3) involved in embryonic development (*LMX1B*) and genome stability (*ZBTB43*; **Supplementary Fig. 26a**). We observed significant up-regulation of *ZBTB43* expression in blood for the Neanderthal genotypes (*P* value < 0.05; *t* test; **Supplementary Fig. 28**). The demographic kinship between EAS and SEA is reflected in their HF shared chunks (**Supplementary Table 5**), encompassing the N1281 (3p21.31) involving *HYAL* gene cluster (skin UV defense) and *SLC38A3*^7,55^ (glutamine transportation), N6440 (16q24.3) related to tooth development (*CPNE7*) and skin pigmentation and aging^56^ (*DPEP1*, *MC1R*, *DEF8*), N4544 (10q23.31) about kidney metabolic abnormalities (*SLC16A12*), and N5340 (12q24.31) about cholesterol metabolism (*SCARB1*; **Supplementary Fig. 26a**). Additionally, 32 HF Neanderthal segments appear restricted to EAS, such as N870 (2q21), which contains *MGAT5*, encoding Golgi enzyme modulating complex N-glycan branching with pivotal roles in oncogenesis and immune regulation^57^ (**Supplementary Table 5**).

A notable example is N12 (1p36.32), a chunk present in all non-AFR superpopulations but reaching substantially higher frequencies in East Eurasia (**Fig. 4a**). This region harbors *PRDM16*, a master metabolic regulator that governs the shift toward thermogenic brown and beige fat, a critical survival mechanism against environmental cold^58,59^ (**Supplementary Fig. 29**). We identified 180 informative sites (171 SNVs, 7 InDels and 2 SVs) supporting introgression at this locus, including 37 localized to regulatory elements (e.g. enhancers and CTCF motifs), implying functional impacts on gene expression alteration (**Supplementary Table 21**). Haplotype network and composition confirmed the origination from Neanderthal (**Fig. 4b**; **Supplementary Fig. 30**). Using rs12041848 as a proxy for the archaic haplotype, which located within an enhancer (EH38E1312562) and a CTCF binding site, a strong signature of recent positive selection in EAS on the archaic allele (A) was observed from the Extended Haplotype Homozygosity (EHH) analysis (**Fig. 4c**). Intriguingly, the archaic allele was virtually absent in Western and Central Eurasia, and its frequency exhibits a striking latitudinal gradient (*P* value = 0.005, *R* = −0.584, Pearson correlation test) in East Eurasia, suggesting the gradual selection driven by cold adaptation (**Fig. 4c**). Similarly, N5782 (14q23.1), containing *PRKCH*, is extensively enriched in EAS and SEA (**Extended Data Fig. 3a**; **Supplementary Fig. 31**). This signaling kinase is pivotal for cell differentiation and proliferation, particularly in epithelial and skin tissues^60^. Haplotype network indicated Neanderthal ancestry (**Extended Data Fig. 3b**; **Supplementary Fig. 32**). We identified 25 regulatory and one missense introgression variants, including locus rs17098433 located within both a promoter and a CTCF binding site (**Supplementary Table 21**). The EHH plot strongly indicated the selection to the archaic allele, highlighting the local adaptation (**Extended Data Fig. 3c**). Importantly, the missense variant (rs2230500) is a known risk factor for ischemic stroke in Chinese and Japanese populations^61,62^. The high-risk allele is restricted to the Western Pacific (EAS, SEA and OCE), likely maintained through genetic hitch-hiking with adjacent linked adaptive loci under selection (**Supplementary Fig. 33**).

**Fig. 4.**
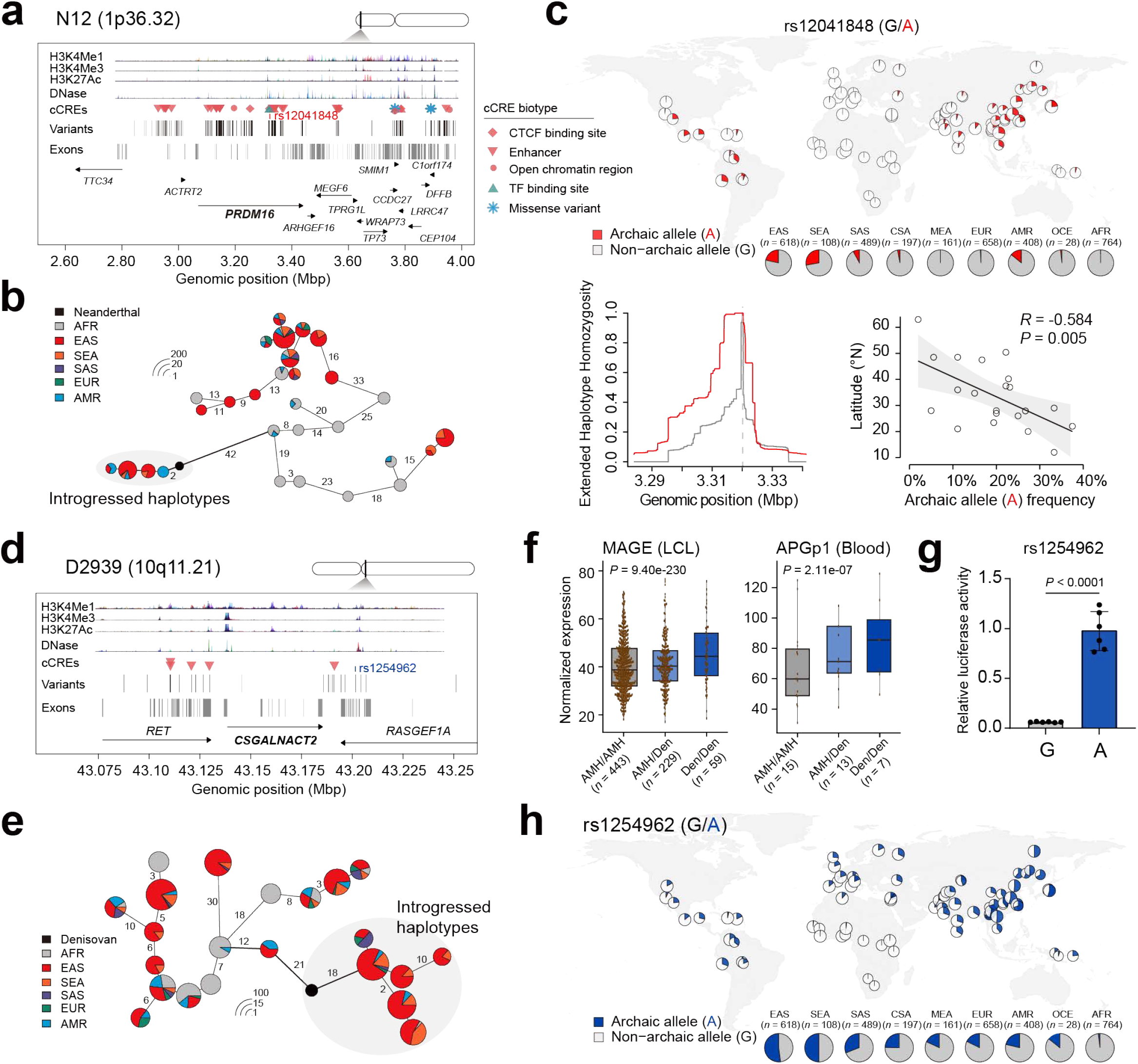
Haplotype characterization of adaptive archaic introgression at the *PRDM16* and *CSGALNACT2* loci. **a** and **d**, Genomic architecture and regulatory landscapes of the focal introgressed archaic chunks. a, Neanderthal chunk N12 (1p36.32). d, Denisovan chunk D2939 (10q11.21). In both panels, all introgressed variants are marked by vertical black lines, with variants overlapping candidate *cis*-regulatory elements (cCREs) further indicated by distinct symbols according to their predicted regulatory class. Tracks for chromatin accessibility (DNase I hypersensitivity), and histone modification signals (H3K27ac, H3K4me3, H3K4me1) combined from all cell types of ENCODE are shown. **b** and **e**, Haplotype networks of core adaptive introgressed regions. Networks are built from the most frequent haplotypes in the Neanderthal chunk N12 and Denisovan D2939 chunks. Each node represents a unique haplotype; its radius is proportional to log₂(number of carriers) plus a minimum offset for visibility. Pie sections show the superpopulation composition. Edge lengths correspond to the number of pairwise differences between connected haplotypes. **c**, Geographic distribution, selection and geographical association of a selected Neanderthal-derived variant rs12041848, using the 1KGP and HGDP data. **F**, Association between Denisovan introgression and *CSGALNACT2* expression. Boxplots show normalized expression grouped by genotype at the proxy locus rs1254962. Center lines represent the median, boxes show the interquartile range, and whiskers extend to the full expression range. **g**, Dual-luciferase reporter assay for rs1254982 (G/A) in HEK293-T cells. Data are presented as mean ± s.e.m. Significance levels are calculated using two-sided *t*-test (*n* = 6 replicates for each of the two groups). **h**, Geographic distribution of the Denisovan-derived rs1254982 allele, across global populations from the 1KGP and HGDP data.

In stark contrast to the widespread Neanderthal contribution, Denisovan contribution to modern humans is considerably more circumscribed. Only 17 Denisovan-derived chunks reached high frequency in EAS, of which 12 are shared with SEA (**Supplementary Fig. 26b**; **Supplementary Table 6**). Totally 21 out of 299 introgression sites showed positive selection signals in EAS (**Supplementary Fig. 27**). Besides the well-known loci, such as *TBX15*-*WARS2* locus (1p12, D201) associated with cold adaptation^63^, and *S100A10*-*FLG2* (1q21.3, D221) within the Epidermal Differentiation Complex (EDC) region linked to skin barrier establishment^22^, we noticed several novel Denisovan segments (**Supplementary Fig. 26b**). A prominent discovery is the D2939 (10q11.21) chunk encompassing *CSGALNACT2*, which encodes a chondroitin N-acetylgalactosaminyltransferase, essential for chondroitin sulfate synthesis (**Fig. 4d**; **Supplementary Fig. 34**). Haplotype network analysis supported its introgression from a Denisovan source (**Fig. 4e**; **Supplementary Fig. 35**). We observed 22 introgressed small variants including five in the regulatory enhancer regions, and a 1.4-kbp intronic deletion spanning an *Alu* element and ERVK (**Supplementary Table 21**). Blood and lymphoblastoid cell line (LCL) expression data suggested these introgressed variants function as *cis*-eQTLs up-regulating the expression of *CSGALNACT2* (**Fig. 4f**). To pinpoint the causal variants underlying the expression alteration, we performed luciferase reporter assays on 22 introgression variants. While some variants exhibited inhibitory effects, the archaic allele (A) of rs1254962 demonstrated significant transcriptional activation (**Fig. 4g**; **Supplementary Fig. 36**). Micro-C chromatin maps further confirmed a physical interaction between the downstream introgression locus and *CSGALNACT2* promoter (**Supplementary Fig. 34**). Notably, this locus is located within a ZNF707 binding motif site, situated near a cluster of enhancers (marked by H3K27ac and H3K4me3; **Supplementary Fig. 34**). This G-to-A transition likely reduces the inhibitory effect of ZNF707 on downstream enhancers, therefore activating the expression of *CSGALNACT2*, like an insulator loss^64^. Using rs1254962 as a proxy for the archaic haplotype, we observed a frequency gradient across Asia, peaking in EAS (0.52) and SEA (0.50), with diminishing frequencies in South and Central Asia (0.31 and 0.25; **Fig. 4h**). The EHH analysis further evidenced longer haploblocks for the archaic allele, confirming a strong selection (**Supplementary Fig. 37**). Another striking example is the metabolism-associated gene *CH25H* from chunk D3015 (10q23.31), a hub integrating lipid metabolism and immune response^65^ (**Extended Data Fig. 3d**; **Supplementary Fig. 38**). Distinct archaic-type haplotypes were observed in non-AFR populations (**Extended Data Fig. 3e**; **Supplementary Fig. 39**), with the representative promoter variant rs4417181 (T/C) reaching substantially high frequency of archaic allele in East Eurasia and positive selection (**Extended Data Fig. 3f**).

### Three Denisovan introgression pulses in East Eurasian genomes

The geographic and temporal distribution of Denisovans across East Eurasia and Oceania, documented through both fossil and DNA evidence, suggests a complex history of interactions with modern humans^66–72^. Multiple introgression events have been proposed based on genetic evidence, but the number, timing and geographic origins of these admixture pulses have remained contentious^11,22,73^. Here we systematically assessed the sequence identity of Denisovan-introgressed segments across haploid assemblies by calculating the match rate (MR) of each segment to the Altai Denisovan reference genome (**Methods**). Gaussian mixture modeling of MR distributions revealed that 29.0% (78/269) of haploid genomes were optimally modeled by three Denisovan components, providing assembly-based evidence for at least three distinct introgression events (**Fig. 5a**; **Supplementary Figs. 40-42; Supplementary Tables 22** and **23**). This three-pulse signature was independently corroborated using Sprime on short-read data^11^, particularly in Tibetan (CZA) and East Han Chinese (CHA-E) populations (**Supplementary Fig. 43**; **Supplementary Table 24**). Moreover, this three-pulse pattern was also prevalent in SEA assemblies (25.0%; 8/32), where their MR distributions fit well with the models observed in EAS, suggesting shared ancestral events despite differential retention (**Supplementary Fig. 41**).

**Fig. 5.**
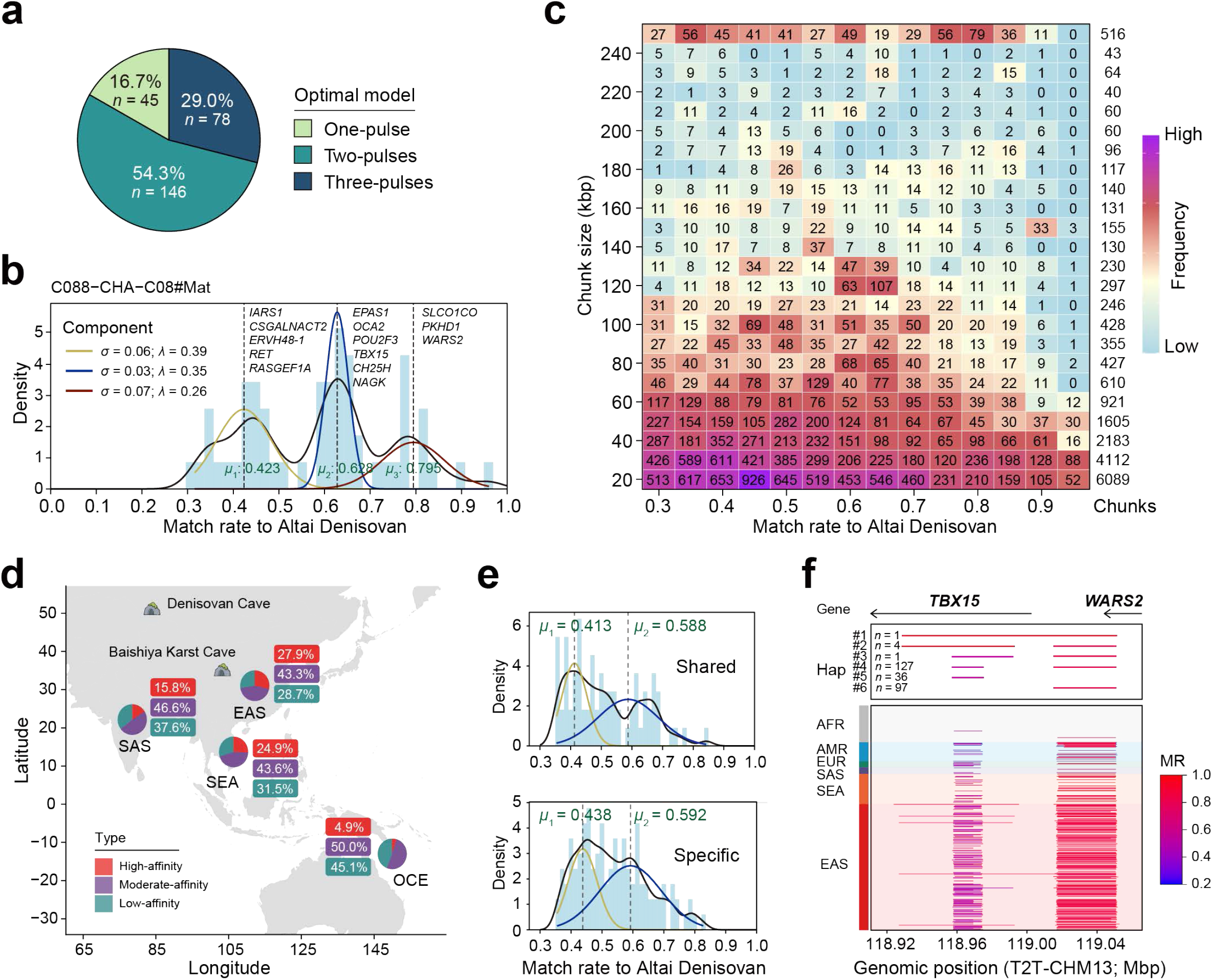
Multiple episodes of Denisovan introgression in East Eurasia. **a**, Gaussian mixture model fitting for the number of Denisovan introgression events across East Asian haploid genomes. **b**, Distribution of match rates to the Altai Denisovan genome for a representative haploid genome (C088-CHA-C08#Mat). The densities are fitted with a three-component Gaussian mixture model (colored curves). Peaks represent distinct introgression events, with vertical dashed lines indicating their means. Selected genes within haplotype segments from each peak are annotated. **c**, Heatmap of introgressed fragment counts, binned by length (*y*-axis) and match rate (*x*-axis). The rightmost column shows the total block count per length bin. The shift towards higher match rates for longer fragments indicates multiple waves of introgression, with recent episodes preserving larger genomic segments. **d**, Geographic distribution of Denisovan ancestry affinity across Asia. Pie charts show the proportion of low, moderate and high-affinity introgressed fragments. The prevalence of high-affinity segments in East Asia suggests a localized origin or southward dispersal. **e**, Comparison of Denisovan match rates in fragments shared between Melanesian and Eurasian populations (top) versus those unique to the Melanesian genome (bottom). Both distributions are best fitted by two peaks. The closely matched mean values (*μ*_1,_ *μ*_2)_ of the shared and unique components suggest that the ‘Melanesian-specific’ ancestry was derived from the same common ancestral source as the shared ancestry, not from a separate introgression event. **f**, A representative Denisovan introgressed haplotype structures (D201 at 1p12). The segment spans chr1:118,923,814-119,052,898 (bp; T2T-CHM13). From top to bottom: gene annotations; six distinct haplotype tracks, labeled by carrier count; and introgressed segments colored by match rate to the Altai Denisovan genome. Among 266 assemblies carrying introgressed segments in this region, haplotypes 4 and 6 are the most prevalent.

We assigned the introgression segments into three pulses in each haploid, by labeling low (0.30 < MR< 0.45), moderate (0.45 < MR < 0.65) and high affinity (0.70 < MR < 1) to the Denisovan reference (**Supplementary Fig. 44**; **Supplementary Table 23**), revealing the origination of several well-characterized adaptive loci from different Denisovan lineages. The moderate-affinity pulse contributed to *EPAS1* for high-altitude adaptation (MR = 0.537), *OCA2* for pigmentation (0.553), *POU2F3* for taste perception (0.544), *TBX15* for body fat distribution (0.523), *CH25H* for lipid metabolism (0.630). The high-affinity pulse introduced *SLCO1C1* for thyroid hormone transport (0.802), *PKHD1* for kidney function (0.897), *WARS2* for mitochondrial function (0.84). The low-affinity pulse contributed *IARS1* for aminoacyl-tRNA synthesis (0.32), *CSGALNACT2* for chondroitin sulfate biosynthesis (0.34), *ERVH48-1* (0.42), *RET* (0.34), and *RASGEF1A* for GTPase signaling (0.33; **Fig. 5b**). These pulses exhibited distinct segment-size distributions that reflected their respective introgression timings (**Fig. 5c**; **Supplementary Figs. 45** and **46**). Notably, ultra-long segments (>250 kbp) showed a significant enrichment around an MR of 0.8, pinpointing the most recent introgression event in EAS. Geospatial mapping of these segments revealed a pronounced concentration of high-affinity segments in EAS, particularly in North China, geographically proximal to the Altai Denisovan Cave, implying an EAS origination for this specific component (**Fig. 5d**; **Supplementary Figs. 47** and **48**).

To understand how these East Eurasian introgression pulses related to previously reported introgression in Papuans^74^ or Oceania populations^73^, we performed a comparative analysis by leveraging a Melanesian genome assembly^1^, that carried 16.55-Mbp Denisovan sequences in 264 chunks (4.04-fold and 4.13-fold in segment count and length, than those in EAS; **Supplementary Table 25**). Gaussian mixture modelling identified two distinct pulses in the Melanesian individual (*μ*_1_ = 0.435, *μ*_2 =_ 0.598, *P* value = 9.54 × 10^-6^), corresponding closely to the low-and moderate-affinity waves observed in Eurasians. This concordance implies these two pulses represent ancient introgression events shared across Oceania and Eurasia (**Supplementary Fig. 49**). Among the 194 Denisovan segments in the Melanesian genome (MR > 0.35, chunk length > 15 kbp), 36 are shared with >10 non-Oceanian assemblies (**Supplementary Table 26**). These shared segments showed no significant difference in MR or length between the Melanesian and Eurasian genomes (*P* value = 0.18 and 0.83, paired *t*-test; **Supplementary Figs. 50** and **51**). Moreover, the 117 Melanesian-specific segments exhibited a bimodal MR distribution (*μ*_1 =_ 0.438, *μ*_2 =_ 0.592, *P* value = 0.00244) and comparable sizes to shared segments, indicating their population specificity likely results from genetic drift due to bottlenecks and reduced effective population size in Oceania population rather than unique recent introgression events (**Fig. 5e**; **Supplementary Fig. 52**). In addition, to align with previous documented waves in Oceania populations, we intersected the Denisovan segments by ASMaid with previously defined pulses D1 and D2 (only >180 kbp available) in Papuans^74^. High-confidence hits suggested their correspondence to the low-affinity pulse for D2 and moderate-affinity for D1 (**Supplementary Table 27**), which implied the sharing introgression between East Eurasia and Oceania populations. We further confirmed this relationship by phylogenetic reconstruction. For example, at the *NPIPB3* locus (16p12.2), the Oceania individuals were nested in the Eurasians among the genomes with Denisovan introgression (**Supplementary Fig. 53**). Collectively, these lines of evidence suggested a model wherein low-and moderate-affinity Denisovan introgression events were shared across Oceanian and East Eurasian populations.

The complexity of Denisovan introgression is further illustrated by examining individual loci where multiple pulses contributed to the same genomic region. Among high-frequency Denisovan segments, 31 exhibited bimodal MR distribution across haplotypes (**Supplementary Table 28**), indicating admixed origination from multiple Denisovan lineages at the same locus. One of the most compelling examples involves the *TBX15*-*WARS2* region (D201) on chromosome 1 (**Fig. 5f**), associated with body fat distribution and brown adipose tissue function^63^. Among the 266 haploid genome assemblies with introgression signals, the most common introgressed haplotype (Hap4, *n* = 127) contains two distinct archaic fragments spanning *TBX15* (∼15 kbp) and *WARS2* (∼34 kbp) with markedly divergent MRs of 0.53 and 0.85, respectively, indicating introgression from both moderate-and high-affinity Denisovan lineages. Notably, a singleton haplotype containing an exceptionally long Denisovan segment (∼121 kbp) with a high MR of 0.75 was observed in an EAS individual, suggesting the relict of the latest specific introgression event in EAS. The observed variation in haplotype structure reflects a mosaic of diverse recombination histories and underscores the complexity of archaic segments following introgression.

## Discussion

By leveraging high-quality phased assemblies, we identified a substantially larger reservoir of archaic sequences than reported by previous methods such as IBDmix^13^ and Sprime^11^. This increased sensitivity is attributable to the ASMaid framework to scan haploid assemblies independently, capturing low-frequency or singleton segments typically obscured in population-scale analyses. By operating within self-assembly coordinates, rather than alignment against the easy region of a single reference (commonly GRCh38 or GRCh37, <80% of whole-genome length), ASMaid bypasses the reference bias inherent in standard linear frameworks while integrating the signals of single-base substitutions and copy-number variations to provide a multi-layered evaluation of introgression diversity. A critical refinement is the recalibration of the ancestral background. While West African populations like the Yoruba from Ibadan, Nigeria, have traditionally served as non-introgressed controls^7,14,68^, emerging evidence of widespread “ghost” introgression across West Africa necessitates a paradigm shift in background selection^13,75^. By employing an East African panel from HGDP as a baseline, we mitigated the impact of African archaic signals while balancing computational efficiency. Although a ten-individual panel is a pragmatic compromise that may not fully eliminate the noise of ILS, it represents a significant step toward a geographically nuanced baseline, particularly for Denisovan components which persist at low frequencies and exhibit high heterogeneity across detection methods^15^.

These methodological advancements specifically enhanced our detection capacity in complex or repetitive genomic regions. One of the most representative findings is the discovery of large centromere-spanning archaic segments with low frequency in modern humans, which also has been reported in recent HPRC and HGSVC studies^32,76^. Unlike typical segments that undergo rapid fragmentation via recombination, the suppressed recombination of centromeres enables the preservation of ancient haplotypes across hundreds of thousands of years. While the functional consequences remain to be elucidated, the distinct HOR compositions of archaic centromere haplotypes could potentially influence chromosome segregation, stability, or meiotic drive^77–79^. Furthermore, by comparing introgressed and non-introgressed homologous sequences, we identified a broad spectrum of putatively archaic-derived structural variants. These findings suggest that archaic introgression provided structural “modules” that likely exerted more profound phenotypic effects than single nucleotide changes, though their specific functions require further experimental validation.

The spatiotemporal complexity of Denisovan introgression remains a major point of contention, partially due to the historical lack of a unified analytical framework across diverse global populations^10,80^. While Sprime analysis suggested two waves in EAS, with one high-affinity pulse with a match rate of around 0.8 and the other moderate-affinity of 0.5, South Asian and Papuan populations have archaic components with moderate affinity^11^. Subsequently, Gaussian mixture modeling analysis decomposed the affinity distribution into two components with close affinity to Denisovan reference in Oceania or Papuan populations, but distinct from the recent high-affinity component predominant in EAS^73,74^. Here, our implementation of the ASMaid framework across 610 assemblies provides evidence for a more intricate three-pulse model. By decomposing match rate distributions, we identified two low-to-moderate affinity pulses shared between Eurasians and Oceanians, alongside a third, high-affinity pulse predominant in EAS. This aligns with recent observations from mainland Southeast and South Asians^22,23,81^, suggesting that previously “missing” components were obscured by the technical limitations.

The sharing of archaic components between Oceanian and Eurasian populations invites two competing hypotheses: introgression in a common ancestor residing in Southeast Eurasia versus a later back-migration from Oceania to Eurasia10. However, resolving this depends fundamentally on the assumed phylogenomic kinship—specifically whether Oceanians represent a sister group to EAS75,82 or an outgroup to the EAS-EUR split73,74—as simulation-based inferences like fastsimcoal2 or Approximate Bayesian Computation (ABC) are exceptionally sensitive to such topological priors. A misplaced phylogeny can lead to significant misinterpretations of gene flow and effective population sizes. Furthermore, the mosaic state of identified archaic segments, exemplified by the co-introgression of divergent lineages at the TBX15-WARS2 locus, reveals that Denisovan introgression was not a sequence of isolated events but a “braided stream” of genetic exchanges. These segments are subject to heterogeneous selective pressures and regional recombination, further complicating demographic inference. Resolving these intricate interplays requires a transition to unified frameworks that incorporate the full spectrum of global diversity, particularly underrepresented populations, to systematically map the archaic contributions shaping human dispersal and phenotypic variation.

## Acknowledgements

We thank all members of the Asian Pan-Genome project for their insightful suggestions. This study was funded, in part, by the National Key R&D Program of China (2024YFA1802500), Fundamental and Interdisciplinary Disciplines Breakthrough Plan of the Ministry of Education of China (No. JYB2025XDXM508), the funding from International Institutes of Medicine of Zhejiang University (KY2022-098), Basic Research Center Program (32388102), the New Cornerstone Science Foundation through the XPLORER PRIZE and the K.C.Wong Education Foundation to G.Z., the National Key R&D Program of China (2025YFC2708200) and the National Natural Science Foundation of China (No. 82394424) to D.Z., the National Key R&D Program of China (2025YFC3410300) to D.W., the China Postdoctoral Science Foundation (2024M752766) and the Postdoctoral Fellowship Program (Grade C) of the China Postdoctoral Science Foundation (GZC20241502) to A.B.

## Author contributions

G.Z. and D.W. conceived and designed this study. G.Z. and D.Z. supervised this study. M.S. and D.W. designed and implemented the ASMaid framework. M.S. and A.B. evaluated the performance of ASMaid and profiled the archaic introgression landscape for global human genomes. M.S., Y.S., A.L. and L.N. analyzed the archaic-like centromere haplotypes. Q.C., H.W. and D.W. performed the analysis of archaic-derived structural variation and functional enrichment. A.B., D.Y., R.C. and Y.Y. analyzed population stratification and haplotypes. A.B., M.S. and D.W. performed the analysis of multiple Denisovan introgression pulses. L.J., Y.H., Q.Y., X.W., H.W., Y.S. and D.Z. contributed to data interpretation and provided critical revisions. D.W., M.S., A.B., Q.C. and G.Z. wrote the manuscript, with input and final approval from all authors.

## Competing interests

All authors declare no competing interests.

## Methods

### Detecting archaic introgressed segments with ASMaid

#### Overview

To identify archaic introgressed segments in high-quality human genome assemblies, we developed ASMaid, an analytical framework built on Hidden Markov Model (HMM) that integrates both sequence and structural variation signals. The pipeline accepts multi-sample VCF files generated by aligning short-read sequencing data from archaic hominins and contemporary human populations to targeted assembly. In this study, our archaic genomes include two Neanderthal individuals, Altai^75^ and Chagyrskaya^83^, and one Denisovan^68^. To establish a robust introgression-free ancestral background and minimize the confounding effects of ILS, we implemented a stochastic sampling strategy using HGDP dataset^23^. For each run, we randomly selected a composite baseline of ten East African individuals (three from Mbuti, three from Bantu Kenya, two from San, and two from Bantu South African population), to represent the ancestral human lineage with sufficient genetic diversity.

We applied ASMaid to 610 haplotype-resolved genome assemblies sourced from four major projects: APGp1 by APG Consortium^20^, HGSVC3 by The Human Genome Structural Variation Consortium^18^, HPRCy1 by The Human Pangenome Reference Consortium^16^, and SEA3K by the Consortium of Anthropological Research in Southeast Asia and Southwest China^22^, providing a global representation of haplotype-level variation. Genome assemblies from APGp1, HPRCy1 and HGSVC3 are phased using trio information or high-coverage Hi-C or Strand-seq data, while SEA3K samples are partially phased, which may introduce minor biases in local ancestry resolution. ASMaid integrates genotype (GT) likelihoods with read-depth-based copy number (CN) estimation in its HMM framework to refine the boundaries of introgression tracts. To ensure high-confidence calls, candidate segments underwent rigorous post-processing, including filtration of complex repetitive regions and identity-by-state (IBS)-based recalibration.

### Read mapping and variant calling

#### Processing modern human sequencing data

Raw paired-end FASTQ reads were quality-filtered using fastp v0.23.4^84^ (parameters: -u 30 -q 20) to remove low-quality bases and adapters. Filtered reads were aligned to each target assembly using BWA-MEM v0.7.17^85^. Resulting BAM files were filtered with samtools v1.19.2^86^ to retain only primary alignments with mapping quality (MAPQ) ≥ 30 (parameters: -q 30 -F 2048 -F 256). PCR duplicates were marked and removed using sambamba markdup v1.0.0^87^, followed by coordinate sorting and indexing via samtools. Variants were called using DeepVariant v1.6.0^88^ with the WGS model for high-accuracy variant detection across diverse genomic contexts.

#### Processing ancient hominin data

Raw genomic sequences for the Neanderthal and Denisovan were retrieved from the Max Planck Institute for Evolutionary Anthropology repository (https://www.eva.mpg.de/genetics/genome-projects/). To mitigate the specific challenges of ancient DNA degradation, we applied specialized filters: merged and paired-end reads with ≥5 bases of quality <15 were discarded. Base quality recalibration was performed to reduce the quality score of any T bases in the first or last two positions of each read to 2. Processed ancient DNA reads were aligned to the target assemblies using BWA-aln algorithm with parameters optimized for short, damaged fragments (parameters: -n 0.01 -l 16500 -o 2), followed by BWA-samse. Read alignments with MAPQ ≥ 30 were retained and duplicates were removed using sambamba markdup. Variant calling was subsequently performed using GATK HaplotypeCaller in GVCF mode^89^ (v4.1.8.1).

#### Joint calling and VCF processing

Single-sample GVCFs from modern and ancient individuals were integrated into a unified multi-sample VCF using GLnexus v1.4.1^90^. To ensure data quality, we applied the following post-merging filters: (1) Allelic filtering: only biallelic variants were retained; (2) Genomic masking: To minimize false positives in structurally complex regions, we masked centromeres, variable number tandem repeats (VNTRs), short tandem repeats (STRs), and rDNA regions; (3) Mapping coverage normalization: To ensure cross-sample comparability across heterogeneous sequencing platforms, read depth (DP) values were normalized by the global median and stored as a NORM_DP INFO tag in the VCF files. This standardized metric facilitates the identification of copy number (CN)-based supportive sites within the ASMaid HMM framework by providing a consistent empirical distribution.

### ASMaid workflow

#### Hidden Markov Model framework

ASMaid assigns the whole genome into two discrete hidden states: Ancestral (*S*_0_, non-introgressed) and Introgressed (*S*_1_). The model operates on a sequence of observations derived from both genotype (GT) and copy number (CN) signals. The transitions between states are governed by a transition matrix *A*, initialized with biological parameters: recombination rate (*r* ≈ 2.3×10^-8^ per bp per generation), estimated time since admixture (*T* ≈ 2,000 generations), and expected admixture proportion (*m* ≈ 0.02). Transition probabilities are defined as:

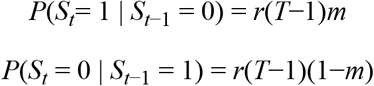

#### Multi-modal emission model

To integrate discrete SNV genotypes and structural variation signals into a single framework, we define a joint observation space *O* ∈ {0,1,2,3} at each genomic site *i* based on the concordance between the target archaic sample and the East African background panel. Specifically, genotype (GT) support (*S_gt_*) is identified at positions where the archaic individual carries a reference allele (*GT_anc_* ∈ {0/0, 0/1}) while the reference frequency in the East African background is below a threshold *τ_gt_* (e.g. *τ_gt_*=0). Parallelly, copy-number (CN) support (*S_cn_*) is defined by sites where the archaic individual maintains a standard diploid state (0.5 ≤ *DP*_anc ≤_ 1.5, where *DP* refers to normalized depth) while a proportion ≥*τ_cn_* (e.g. *τ_cn_*=1) of the background panel exhibits abnormal coverage (*DP* < 0.5 or *DP* > 1.5). Observations are categorized as: *O*=3 (dual support *S_gt_*∧*S_cn_*), *O*=1 (*S_gt_* only), *O*=2 (*S_cn_* only), and *O*=0 (no archaic-specific signals). This multi-modal categorization allows ASMaid to prioritize high-confidence introgressed segments through a hierarchical emission model, where *P*(*O*=3|*S*_1_) >*P*(*O*=1|*S*_1_) > *P*(*O*=2|*S*_1_). By setting *P*(*O*=1|*S*_1_) > *P*(*O*=2|*S*_1_), the model inherently treats genotype calls as more robust indicators than local depth variation, ensuring that depth-based introgression calls require reinforcement through spatial clustering.

#### Model training and posterior decoding

Model parameters, including the emission and transition matrices, are refined using the Baum-Welch algorithm (expectation-maximization) until the log-likelihood converges (tolerance 10^-4^). The most likely state path is decoded using Maximum A Posteriori (MAP) algorithm, with site-specific posterior probabilities calculated. Introgressed segments are defined as continuous runs of sites assigned to the introgressed state (*S*_1_).

#### Weighted match rate for evolutionary inference

To quantify genetic affinity between detected segments and archaic sources, we defined a weighted match rate (*W*). The calculation of the affinity metric *W* is based on informative sites, which are categorized into: 1. Archaic-supportive sites (*n_intro_*): target assembly shares a specific allele or copy-number state with the archaic individual that is rare or absent in the East African background (Observations *O* ∈ {1,3} for GT-based and *O* ∈ {2,3} for CN-based supportive sites). 2. Archaic-divergent sites (*n_div_*): archaic individual carries a derived state relative to the reference assembly (e.g. homozygous alternative genotype 1/1, or a lineage-specific CN).

For each modality (GT and CN), a modality-specific match rate (*R*) was calculated as the proportion of supportive sites among all informative markers:

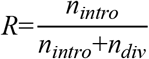

The weighted match rate *W* integrate GT and CN signals, weighted by the number of supportive signals in each modality:

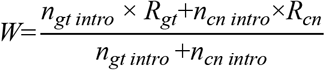

### Refining introgression segment candidates

#### Secondary filtering for complex genomic regions

Candidate segments were cross-referenced with a mask of repetitive/complex regions, including centromeres, VNTRs, STRs and rDNA arrays. Despite pre-filtering variants in these regions, the HMM framework may erroneously link two genuine introgressed signals flanking a repetitive region into a single segment. We excluded segments where >50% of length overlapped masked regions (labeled “MASK”) to mitigate potential mapping artifacts and assembly errors.

#### Integrated IBS metric

A combined identity-by-state (IBS) metric evaluates the homology between each sample and the reference assembly, integrating GT and CN information. 1. GT-based IBS: Proportion of reference alleles (0 alleles) relative to total alleles in the segment. 2. CN-based IBS (*S*_dp)_: To quantify similarity in CN space, we define a depth-similarity function (*S*_dp)_ that maps normalized depth (*v*) to (0,1], where a value of 1 indicates perfect parity with the reference:

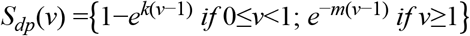

Attenuation coefficients were set to *k*=*m*=2, ensuring exponential decay of similarity toward 0 as normalized depth deviates from 1. The final IBS score is a weighted average of the GT-based and CN-based components, where the weights are proportional to the supportive sites for each modality.

#### Ancestry recalibration

Considering the independent identification of archaic segments in an assembly from Neanderthal and Denisovan, for the commonly identified segments, we utilized a step of ancestry recalibration to phase the archaic origins. Integrated IBS scores are used to assign origins of candidate archaic introgression segment calls, comparing archaic (Altai Neanderthal, Chagyrskaya Neanderthal, and Denisovan) and modern African (East African) genomes. Segments with highest IBS score in any East African individual are labeled “ambiguous” and excluded (removing ILS-related false positives); Segments passing the East African filter are assigned to “Denisovan” if their Denisovan IBS score is highest and exceeds the Altai Neanderthal IBS score by >0.05.

Otherwise, they were labeled “Neanderthal”. Finally, we performed a consistency check against the initial ASMaid predictions. Only segments whose IBS-based ancestry labels matched their ASMaid-predicted archaic origin were retained for subsequent analysis.

### Performance evaluation

#### Simulation

##### Demographic modeling and genomic sequence generation

To rigorously evaluate ASMaid’s performance, we conducted coalescent simulations using msprime v1.3.3^91^ to generate a ground-truth archaic introgression dataset. We constructed a multi-population demographic model encompassing African (AFR), non-African (non-AFR) and Neanderthal lineages, with the following parameters: ancestral effective population sizes (*N_e_*_)_ = 15,000 for AFR, 2,000 for the non-AFR founders, and 1,500 for Neanderthals; Neanderthal-human split at 16,000 generations before present (BP), and out-of-Africa (OOA) dispersal at 2,400 generations BP, and a specific introgression pulse from Neanderthals into the non-AFR population between 2,000 and 1,980 generations BP with a migration proportion of 0.0015, and an exponential growth model (*r* = 0.02) starting 200 generations BP. Using a mutation rate of 2.4×10^-8^ (Jukes-Cantor model) and a recombination rate of 1.0×10^-8^ per base per generation, we simulated 100 Mbp of genomic sequence for 100 non-AFR, 40 AFR and two archaic haplotypes. Ground-truth introgression segments are defined by tracing genealogical tree sequences: a segment is definitively classified as introgressed if its ancestral node resides within the Neanderthal population prior to the split time but during the gene flow interval.

##### Data generation for ASMaid

We initially converted these msprime-simulated tree sequences into a master VCF file comprising haploid genotypes for 100 non-African (non-AFR), 40 African (AFR), and two archaic individuals relative to a unified reference. To generate ASMaid-compatible inputs, we transformed the master VCF into sample-specific diploid VCFs for each non-African individual, effectively simulating the variant representation of short-read mapping against personalized reference genomes. For each target non-AFR haplotype assembly, we polarized all variants in the archaic and African panels relative to the target (treating the target individual as the reference coordinate system). Specifically, if the target carries an alternative allele (1) at a given locus, the alleles in all other samples (Neanderthal and 20 African individuals, formed by merging simulated haplotypes into diploid genotypes) are inverted to maintain the relative allelic state. VCF records where all samples are 0/0 after polarization are filtered out. This approach ensures that the HMM decoding is based on the density of archaic-shared alleles deviating from the target’s specific genetic background.

##### Evaluation metrics

Performance is quantified using seven distinct metrics across two dimensions: length-based and count-based statistics. Length-based metrics included Precision (*L*_overlap/_*L*_call)_, Recall (*L*_overlap/_*L*_true)_, and the F1-score (Precision ×Recall), alongside the False Positive Rate (FPR = *L*_incorrect/_(*L*_total-_*L*_true)_), where *L*_total,_ *L*_true,_ *L*_call,_ *L*_overlap,_ and *L*_incorrect r_epresent the total simulated genomic length (e.g. 100 Mbp), the cumulative length of the ground-truth introgressed tracts derived from the tree sequences, the total length of all segments predicted by the model, the total length of the intersection between the predicted segments and the ground-truth tracts and the total length of predicted segments that do not overlap with any ground-truth introgressed regions, respectively. Complementarily, count-based metrics (Precision, Recall and F1-score) were calculated based on the proportion of predicted segments that overlapped with true segments, ensuring an assessment of both genomic coverage and tract-level detection.

To optimize ASMaid for archaic introgression detection, we systematically evaluated its key parameters and filtration criteria using simulated datasets. First, we assessed the impact of the East African size (*n*_AFR =_ 5, 10, 15, 20) as a background control. While larger panels marginally improved accuracy, we selected *n*_AFR =_ 10 for all downstream analyses to achieve an optimal balance between high F1-scores and computational efficiency. Second, we evaluated the non-supportive genotype frequency threshold-the maximum allowable frequency of the reference allele within the East African panel for a site to be considered archaic-supportive. Simulations indicated that maximum stringency achieved optimal performance; thus, we set this threshold to 0, requiring a complete absence of the reference allele in the East African background. Finally, we refined the post-decoding filters by prioritizing a posterior probability of >0.9 to ensure high-confidence segment calls. Segment length filters were adaptively determined based on the specific genomic context and assembly resolution (detailed in subsequent sections), ensuring robust performance across heterogeneous genomic environments.

##### Comparative analysis with IBDmix and Sprime

To benchmark ASMaid, we compared its performance with two established population-based introgression detection methods, IBDmix^13^ and Sprime^11^. Publicly available introgression maps are retrieved from https://zenodo.org/records/14552025 (based on the T2T-CHM13 reference) for IBDmix^92^, and from https://data.mendeley.com/datasets/y7hyt83vxr/1 for Sprime (originally based on hg19) with subsequent harmonization to the T2T-CHM13 coordinate using liftOver (v362; https://hgdownload.soe.ucsc.edu/admin/exe/linux.x86_64/liftOver). Given that these methods primarily utilized the 1000 Genomes Project (1KGP) short-read data, we restricted initial comparison to 15 non-AFR individuals overlapping between 1KGP and HPRCy1/HGSVC3. We further extended IBDmix and Sprime to the APGp1 dataset, utilizing short-read sequencing data.

To align with ASMaid’s haplotype-resolved predictions, ASMaid segments with a posterior probability >0.9 are retained, and maternal and paternal calls are merged for each individual. For Sprime, Neanderthal segments are defined by a match rate >0.6 to the Altai Neanderthal and <0.4 to the Altai Denisovan, while Denisovan segments require a match rate >0.4 to the Altai Denisovan and <0.3 to the Altai Neanderthal, as previously described^11^. For IBDmix, segments with a length >50 kbp and a LOD score >4 are kept. For performance comparison using simulation data, we doubled the simulated non-AFR haploids into homozygous diploid genotypes, enabling direct comparison of ASMaid’s haplotype-resolved sensitivity with IBDmix and Sprime.

### Global profiling of archaic introgression segments

#### Filtration strategy

We ran ASMaid for each of 610 haploid assemblies from APGp1, HPRCy1, HGSVC3 and SEA3K. To ensure high-confidence detection of archaic introgression, we filtered raw segment calls for each assembly with a posterior probability of >0.9. To facilitate comparison, the coordinates of archaic segments across assemblies were aligned to the T2T-CHM13 via Liftoff v1.6.3^93^. Introgression segments across all 610 assemblies were merged and aggregated into non-redundant genomic chunks, labeled with a chunk ID with prefix “N” and “D” representing Neanderthal and Denisovan chunks, respectively. A chunk was flagged as a potential ILS product and excluded if it met either of the following criteria: the proportion of African introgressed individuals among all carriers of the chunk ≥60%, or ≥3 individuals in the African LWK population (Luhya in Webuye, Kenya, which represent the closest to an introgression-free East African background) carried the chunk, with a cumulative introgressed length proportion >50%. For the comparative statistics presented in Fig. 2a, an additional length filter (>20 kbp) was applied. Conversely, downstream assembly-level analyses retained all segments (post-initial filtration) to capture fine-scale introgression landscapes.

#### Frequency and heterogeneity

We classified the identified chunks into five categories based on their prevalence across the 610 assemblies: singleton (*n* = 1), rare (frequency <1%), low-frequency (1-5%), common (5-40%), and high-frequency (≥40%). To assess the saturation of the archaic sequences, we performed growth curve analyses. For cumulative length, samples were iteratively added, recalculating the union of introgressed sequences and dynamically updating frequency labels. For cumulative chunk counts, we utilized the final 610-sample aggregate as a reference set, determining the proportion of the total “introgressed pangenome” captured with each incremental addition. Inter-assembly similarity was quantified using the Jaccard Index, defined as the intersection of introgressed lengths between two haploid assemblies divided by their union. To evaluate the heterogeneity within parental pairs, observed values were compared against random pairs from the same population.

#### Inferring the source of archaic segments in African genomes

To delineate the spatial source of archaic ancestry in AFR, all putative archaic segments in AFR genomes were merged and compared against a merged set of non-African segments. To account for the substantial variation in sample sizes across superpopulations, a downsampling approach was employed. 16 haploid assemblies were randomly sampled from each superpopulation to standardize the discovery power. The overlap in total introgressed sequence length across standardized subsets was then calculated to evaluate the proportion of archaic segments shared between African genomes and other superpopulations.

#### CN-based introgression call at the *MGAM* locus

The *MGAM* loci across assemblies were annotated using Liftoff v1.6.3 from the GRCh38.p14 (GENCODE v47) reference^93^. Comparative analyses were restricted to assemblies containing the MANE Select transcript. Coding sequences (CDS) were extracted and translated via gffread v0.12.8 (https://github.com/gpertea/gffread), followed by protein domain identification using HMMER v3.4 (https://www.ebi.ac.uk/Tools/hmmer/home) against the Pfam database (E-value < 1×10^-5^). Structural variations and synteny of *MGAM* genes between human and chimpanzee assemblies (GCA_028858775.2) were identified using minimap2 v2.24^94^ and visualized with SVbyEye^95^. To facilitate cross-assembly copy number (CN) comparison, the *MGAM* region was partitioned into 100 bins. For each assembly, we calculated the mean normalized depth per bin for both the East African and Altai Neanderthal populations.

### Centromere-spanning archaic introgression segments

#### Identifying centromere-spanning candidates

Despite excluding variants in highly repetitive regions (centromeres, VNTRs/STRs, and rDNAs), long-range signals from flanking regions can facilitate HMM transitions across these gaps, resulting in anomalously long introgression segments. To systematically characterize these events, we implemented a specialized curation strategy. For non-acrocentric chromosomes, segments were classified as centromere-spanning if the HMM-inferred introgression track physically spanned the centromeric gap, with at least three high-confidence supportive variants on both p-arm and q-arm flanks. For acrocentric chromosomes (13, 14, 15, 21 and 22), given the frequent p-arm truncation in genome assemblies, potential introgression was defined by segments overlapping the first 200 kbp of the q-arm adjacent to the centromere boundary.

#### Phylogenetic analysis

To confirm the archaic origin of centromeric candidates, we performed localized phylogenetic reconstruction using homologous introgressed flanking sequences with stringent filters. The 100-kbp regions flanking T2T-CHM13 centromeres were lifted over to all 610 assemblies using minimap2 (v2.24). Centromere flanks were retained only if across-assembly length variation was minimal, defined by a coefficient of variation (CV) < 0.05 and a range ratio (max-min) / mean < 0.5 (**Supplementary Table 9**). Minigraph-Cactus v2.9.9^96^ was used to construct a local pangenome graph and the decomposed VCF was merged via bcftools v1.16^86^ with four high-coverage archaic genomes (Altai, Chagyrskaya, and Vindija Neanderthals, and the Denisovan) and two outgroups (*Pan paniscus* and *Pan troglodytes*). Variants were restricted to T2T-CHM13 “Easy” regions and filtered using PLINK v1.9^97^ with parameters: --geno 0.2 --mind 0.2 --maf 0.0005 --vcf-hap-call h. Filtered VCFs were converted to FASTA format and maximum-likelihood trees were inferred using IQ-TREE v2.3.5^98^ (GTR+G+I model, 1000 replicates) and visualized via iTOL (https://itol.embl.de/).

#### Centromere structure

A comparative analysis of higher-order repeat (HOR) composition and transposable element (TE) organization in centromeres of chromosome 5 was conducted, leveraging fine annotations of assembled centromeres from a companion study^31^. HOR variants were categorized into three groups: exclusive to introgressed centromeres, exclusive to non-introgressed centromeres, or shared between both groups. For shared variants, a two-sided Wilcoxon rank-sum test was applied to assess copy-number differences between haplotype groups, identifying HOR variants significantly expanded or contracted in introgressed centromeres. Additionally, genomic coordinates of all annotated TEs intersecting α-satellite arrays were extracted and compared. Structural features unique to introgressed haplotypes, such as lineage-specific insertions, were highlighted. One representative centromere from each of non-introgressed centromere haplotypes (CenHap A-F) was randomly sampled and illustrated.

### Archaic-derived structural variants

#### Detecting putatively archaic-introgression derived variants

To identify putatively archaic-introgression derived variants (pAID-Vars), we constructed pangenome graphs for each introgression chunk using Minigraph-Cactus v2.9.9^96^ (MC), with T2T-CHM13v2 as the reference. Variants were decomposed from the bubbles in the MC graph, and the bubble VCF file was subsequently processed using the VCF preparation pipeline (https://github.com/eblerjana/genotyping-pipelines/tree/main/prepare-vcf-MC), and SV collapsing pipeline (https://github.com/Han-Cao/collapse-bubble). Briefly, bubbles with >20% of the haplotypes carrying missing alleles were removed, and bubbles were decomposed to resolve all nested variant alleles, with annotations added to the ID tag of the INFO field, which contained IDs encoding all nested variant alleles of each of the bubble vcf records. Furthermore, variants were refined by vcfwave v1.0.13^99^, followed by SV merging to remove redundancy, concatenation of overlapping variants, and merging of genotypes for duplicated records, resulting in a deduplicated and non-overlapping VCF file. Additionally, the inversions were identified using a hybrid calling approach. Non-redundant inversion calls were integrated from two callers PAV^100^ (v2.4.6) and LSGvar (v1.0.0, https://github.com/Hanjunmin/LSGvar) using bedtools intersect (v2.31.1) with the parameter: -f 0.5 -r. Subsequently, for each variant, allele frequencies were compared between introgressed and non-introgressed haplotypes using Fisher’s exact test. Variants with *P* values <5e-8, were designated as pAID-Vars.

#### Genotyping pAID-Vars in Neanderthal and Denisovan genomes

For small variants (pAID-SMVs, including SNVs and InDels), we first called variants using GATK (v4.1.8.1) for three Neanderthal (Altai, Chagyrskaya, and Vindija) and one Denisovan genomes, and merged the calls with GLnexus (v1.4.1). Since the T2T-CHM13 reference may represent either a modern or an archaic allele at specific loci, we employed different validation strategies. When T2T-CHM13 represents a non-archaic allele, genotypes were initially fixed to “1/1” for all variant records, and then compared to the pAID-SMVs utilizing truvari v5.3.0^101^ with the option “-r 250 -p 0.5 -P 0.5 -t --pick multi -d -s 1 -S 1”, considering potential inconsistent representations of the same InDel variants. Those pAID-SMVs tagged as “TP” (true positive) in the output were designated as high-confidence (HC). When T2T-CHM13 represents an archaic allele, such pAID-SMVs theoretically should not appear in variant calling results if fixed in archaic hominins. However, definitive identification remains challenging due to ancestral polymorphisms and potential calling errors arising from limited availability of high-depth archaic WGS data. To maintain stringency, we only extracted variants with homozygous alternative genotypes (“1/1”) called from at least one archaic genome, and compared them to pAID-SMVs.

For structural variants (pAID-SVs), validation was performed by analyzing mapping coverage profiles and sequence clipping signals from archaic genomes. Archaic WGS data were aligned to T2T-CHM13 using BWA-ALN and BWA-MEM (v0.7.17) for collecting mapping depth and clipping information, respectively. When T2T-CHM13 represents a non-archaic allele, a pAID-SV was defined as HC if the mapping depth was less than 50% of the whole-genome average or if clipping signals were observed. Owing to the low coverage of the Vindija and Chagyrskaya genomes, only the Altai Neanderthal was utilized for depth-based assessments. When T2T-CHM13 represents an archaic allele, a pAID-SV was classified as HC if the mapping depth exceeded 50% of the whole-genome average or if the count of clipping reads were less than half of the whole-genome depth.

#### Structural modelling of the AKR1C8 protein

To elucidate the structural consequence of the 236-bp Neanderthal-derived deletion, we predicted the tertiary structures of both the wild-type (WT) and the truncated AKR1C8 protein. Modelling was performed using AlphaFold (3.0.1)^102^, simulating the protein in complex with the cofactor NADP^+^ and the ligand PGD2. The predicted models were subsequently subjected to energy minimization and structural refinement using PyRosetta v2025.06^103^ to optimize side-chain geometries and thermodynamic stability. Structural alignment and molecular visualization were performed using UCSF ChimeraX v1.10 RC^104^.

#### Gene expression

Expression analysis on archaic introgression genes was approved by the Biomedical Research Involving Humans Ethics Committee in the Fourth Affiliated Hospital of Zhejiang University School of Medicine (Approval No. K2022126) and the Ethics Committee in Women’s Hospital, School of Medicine, Zhejiang University (Approval No. IRB-20260088-R). We generated RNA-seq expression data from blood samples of 35 APGp1 individuals. Peripheral blood was collected from participants using PAXgene Blood RNA Tubes (PreAnalytiX GmbH, Hombrechtikon, Switzerland; Cat. no. 762174). For each individual, 1 ml of whole blood was obtained and stabilized according to the manufacturer’s instructions. Total RNA was extracted using TRIzol reagent (Tiangen Biotech, Beijing, China; Cat. no. DP424) following the manufacturer’s protocol, and quantified using a Qubit® Fluorometer (Thermo Fisher Scientific, USA) with the Equalbit RNA HS Assay Kit (Invitrogen, USA; EQ211-02, Vazyme). RNA integrity was verified using an Agilent 2100 Bioanalyzer (Agilent Technologies, USA). High-quality RNA samples were used for library construction according to standard protocols for poly(A)-selected mRNA sequencing. Libraries were sequenced on the DNBSEQ-T7 platform (BGI, Shenzhen, China) to generate paired-end reads.

Clean paired-end RNA-seq reads were aligned to T2T-CHM13 following GTEx v8 pipeline (https://github.com/broadinstitute/gtex-pipeline) using STAR v2.7.10b^105^ with specified “--twopassMode Basic --waspOutputMode SAMtag --varVCFfile sample.vcf” for each sample and then quantified with RNA-SeQC v2.4.2^106^. Gene expression levels were normalized using the R package edgeR v4.6.3^107^. Specifically, normalization factors were determined via the function *calcNormFactors* using the TMM (Trimmed Mean of M-values) method. Subsequently, the normalized expression values were obtained as counts per million (CPM) via the function *cpm* with the parameter “log = F”. Expression dataset MAGE were generated from lymphoblastoid cell lines in 1KGP^108^.

#### Genetic linkage and extended haplotype homozygosity (EHH)

Linkage disequilibrium (LD) was calculated using vcftools v0.1.17^109^, and visualized using LDBlockShow v1.41^110^. The EHH for each pAID-SV was calculated using the function *calc_ehh* in the R package rehh v3.2.2^111^ and the EHH plots were generated using the *plot* function to evaluate the signals of positive selection.

### Functional and clinical implication

Functional enrichment analysis for high-frequency introgressed genes was performed for Gene Ontology (GO) terms and Kyoto Encyclopedia of Genes and Genomes (KEGG) pathways using clusterProfiler v4.8.3^112^. Diseases association enrichment was conducted against the OMIM, KEGG, and NHGRI-EBI GWAS catalogs with KOBAS v3.0^113^. Risk alleles for early-onset schizophrenia were determined based on a cohort of 1,251 cases and 1,123 controls from one companion study in APGp1^49^.

### Population stratification

#### Selection

To detect signatures of positive selection within high-frequency introgression chunks, we calculated the number of segregating sites by length (nSL) statistic for all bi-allelic SNVs with a minor allele frequency (MAF) ≥ 0.01 using selscan^114^ (v2.0.0) with default parameters (--nsl, --maf 0.01). Raw scores were calculated on a per-SNV basis and normalized within allele frequency bins using the accompanying norm program. SNVs with |nSL| > 2 were considered as candidates for positive selection. Genes containing such candidate SNVs were considered under putative selection. To further assess geographic patterns, the global distribution of representative archaic variants was visualized using 1KGP and HGDP datasets^115^.

#### Functional annotation

To prioritize potentially functional archaic variants, we annotated these variants using the Variant Effect Predictor^116^ (VEP, v.95) to determine the most severe consequence for each transcript. To identify regulatory roles, we intersected these variants with a comprehensive set of regulatory annotations from ENCODE, including candidate cis-regulatory elements (cCREs), DNase I hypersensitivity sites, and histone modification marks for active enhancers (H3K27ac), active promoters (H3K4me3), and poised/active regulatory elements (H3K4me1). Variants overlapping these regulatory features, as well as non-synonymous coding variants, were considered of high priority.

#### Haplotype network

To characterize the evolutionary relationships among introgressed haplotypes, we delineated core regions exhibiting clear stratification between archaic and modern human haplotypes. Using the geneHapR package^117^, we generated haplotype summaries for 610 haploid genomes together with the relevant archaic (Neanderthal or Denisovan) reference genome via the table2hap and hap_summary functions, and subsequently constructed a pairwise genetic distance matrix using get_hapNet. To simplify visualization, only the most common haplotypes for each introgressed chunk were retained for network plotting by plotHapNet.

#### Luciferase reporter assays

To evaluate the regulatory effects of candidate SNPs, luciferase reporter assays were performed using constructs derived from the pGLO plasmid (Fubio Biopharmaceutical Technology Co., Ltd., Suzhou, China). For each of the 22 variants in the region of *CSGALNACT2*, DNA fragments harboring the corresponding allelic variants were synthesized and cloned into the reporter vector using the Hieff Clone® Plus One Step Cloning Kit (Yeasen Biotechnology, Shanghai, China; Cat. no. 10911ES50) according to the manufacturer’s instructions. All constructs were verified by Sanger sequencing prior to functional assays. HEK293T cells (Cyagen Biosciences Inc., Suzhou, China) were cultured at 37 °C in a humidified incubator with 5% CO₂ and seeded into 96-well plates at a density of 1×10⁴ cells per well. Cells were transfected using Lipofectamine™ 3000 Transfection Reagent (Thermo Fisher Scientific, Waltham, MA, USA; Cat. no. L3000008) together with P3000™ Enhancer Reagent (Thermo Fisher Scientific). For each well, 100 ng of plasmid DNA was co-transfected with 0.2 μl P3000 reagent and 0.3 μl Lipofectamine 3000 following the manufacturer’s protocol. After 6 hours, the medium was replaced with a fresh complete medium, and cells were incubated for an additional 48 hours. Luciferase activity was measured using the Dual-Glo® Luciferase Assay System (Promega, Madison, WI, USA; Cat. no. E2920) according to the manufacturer’s instructions. Luminescence was detected using a Multiskan™ FC Microplate Photometer (Thermo Fisher Scientific; Cat. no. 51119000) with a 2-s integration time per well. Firefly luciferase activity was normalized to Renilla luciferase activity to control for transfection efficiency. Relative reporter activity was calculated relative to the internal control construct. Statistical significance between groups was assessed using a two-tailed unpaired *t*-test.

### Inferring Denisovan introgression pulses

#### Inference of introgression pulses by Gaussian mixture modeling

To determine the optimal number of Denisovan introgression events per haploid genome, we analyzed the distribution of match rates (MR) to the Altai Denisovan reference for each of the 610 haploid assemblies. To ensure sufficient statistical power, we retained only assemblies with at least 50 introgressed segments longer than 20 kbp. For each assembly, we fitted one-, two- and three-component Gaussian mixture models, truncated at MR = 0.3 to exclude segments with atypically low similarity to the archaic reference. Parameters (mean *μ*, standard deviation *σ* and mixing weight *λ*) were estimated by maximum likelihood using R package mixtools^118^.

Model selection was performed using likelihood-ratio tests (LRT). The test statistic for comparing a *k*-component model against a (*k*−1)-component model is Λ = 2(ln*L_k_* _−_ ln*L_k_*_−1)_, which asymptotically follows a χ² distribution with three degrees of freedom (the number of additional parameters: *μ*, *σ* and *λ*). A model with *k* components was considered a significant improvement over the (*k*−1)-component model if the LRT *p*-value was <0.05. We applied this criterion hierarchically: the three-component model was accepted if it significantly outperformed the two-component model; otherwise, the two-component model was tested against the single-pulse model.

Robustness checks across varying segment-length thresholds (10-25 kbp) and minimum segment-count cutoffs (30-60) confirmed that the proportion of East Asian haploid genomes supporting three pulses remained stable. Final analysis identified 90 haploid genomes from EAS, SEA and AMR superpopulations meeting the criteria above (≥50 segments, ≥20 kbp), that were optimally fitted by a three-component model. Based on the distribution of fitted means (*μ*₁, *μ*₂, *μ*₃) across these 90 assemblies, we defined three affinity classes: low (0.3 < MR < 0.45), moderate (0.45 < MR < 0.65) and high (0.7 < MR < 1.0). This classification was used to assign each introgressed segment to one of the three inferred pulses. Geographic mapping of three-pulse frequencies revealed a latitudinal cline within EAS, with highest proportions in northern China and a southward decline, suggesting that the core region of the most recent (high-affinity) Denisovan introgression was geographically proximal to the Altai and Baishiya Karst caves.

#### Comparing Denisovan segments between Melanesian and Eurasian genomes

To investigate whether the low-and moderate-affinity Denisovan pulses observed in East Eurasians are shared with Oceanian populations, we leveraged a high-coverage Melanesian genome assembly (HGDP00550) generated via PacBio single-molecule real-time (SMRT) sequencing data^1^ (mean coverage 75.2×; subread length ∼18.2 kbp). This assembly was treated as a pseudo-haploid representation. Using the ASMaid pipeline, we identified 173 Denisovan-derived segments in the Melanesian genome (MR >0.35, length >20 kbp). Gaussian mixture modelling was performed on the MR distribution of these segments. The optimal model was determined by likelihood-ratio tests comparing nested models (three-vs. two-component, and two-vs. one-component), with significance assessed at *P* < 0.05. Genomic coordinates of archaic segments were lifted to T2T-CHM13, and compared with Eurasian introgression sequences. For optimal overlap with Eurasian segments, we relaxed the length threshold to >15 kbp, resulting in 194 segments for comparative analysis. Concordance in MR and segment length for shared segments was assessed using paired *t*-tests. We note that, because the Melanesian genome is a pseudo-haploid assembly, the results should be interpreted with appropriate caution regarding potential phasing or heterozygosity artifacts. To further validate the evolutionary relationship, we performed phylogenetic reconstruction on shared segments using genotype data from the Simons Genome Diversity Project^82^ (SGDP). Maximum-likelihood trees were inferred with IQ-TREE v2.3.5 under the GTR+G+I substitution model with 1,000 ultrafast bootstrap replicates^98^.

#### Aligning with previously reported Denisovan pulses in Papuan

A previous study identified two Denisovan introgression waves in Papuans, termed D1 (recent) and D2 (older), based on mismatch index^74^ (*m*_D)_. We sought to align our MR-based pulses with these events, to examine their sharing or independence. We first lifted the hg38 coordinates of introgression segments (84 segments for D1 totalling 21.0 Mbp; and 80 for D2 totalling 21.6 Mbp; all longer than 180 kbp as provided) to T2T-CHM13. Haploid-level chunks from our study were intersected with these D1 and D2 intervals. Although 20 D1 and 30 D2 segments could be hit by our chunks, we applied stringent filters to ensure high confidence: matched length > 50 kbp, accounting for >30% of the haploid chunk, and >15% of the D1/D2 segment. Singleton hits were further removed. After filtering, seven D1 and seven D2 segments were retained for comparative analysis.

#### Identifying loci with multiple introgression pulses

To investigate genomic regions contributed by multiple Denisovan lineages, we focused on high-frequency introgression segments shared across haplotypes. We selected chunks (>15 kbp) present in at least 50 haploid assemblies, retaining only segments that span more than 10% of the total chunk length. Out of 180 high-frequency Denisovan chunks, 31 (17.2%) exhibited a bimodal match-rate (MR) distribution, confirmed by two-component Gaussian mixture models (likelihood-ratio test *P* < 0.05). Haplotype structures were further dissected to resolve the evolutionary history of multiple pulses at the same locus, as exemplified by the *TBX15*-*WARS2* region.

## Code availability

The custom scripts and computational pipelines used in this study are publicly accessible via Github at https://github.com/Asian-Pan-Genome/ArchaicIntrogression. The ASMaid is accessible at https://github.com/Asian-Pan-Genome/ASMaid under the MIT license. Additional software used in this study, including Sprime (version: 20May22.855) and IBDmix (v1.0.1), can be obtained from their respective repositories: https://github.com/browning-lab/sprime and https://github.com/PrincetonUniversity/IBDmix.

## Data availability

Genome assemblies of HPRCy1 are downloaded from the Human Pangenome Reference Consortium at https://data.humanpangenome.org/assemblies. HGSVC3 assemblies are downloaded from https://ftp.1000genomes.ebi.ac.uk/vol1/ftp/data_collections/HGSVC3/working/. Assemblies from SEA3K are provided by the Consortium of Anthropological Research in Southeast Asia and Southwest China (CASEAC). The Melanesian genome assembly is available at NCBI under BioProject ID PRJNA522307. WGS short reads of HGDP individuals are sourced from https://ftp.1000genomes.ebi.ac.uk/vol1/ftp/data_collections/HGDP/data/. Genotypes of SVs and small variants for 1KGP individuals are downloaded from https://ftp.1000genomes.ebi.ac.uk/vol1/ftp/data_collections/1KG_ONT_VIENNA/release/v1.1/shapei t5-phased-callset/ and https://s3-us-west-2.amazonaws.com/human-pangenomics/index.html?prefix=T2T/CHM13/assemblies/variants/1000_Genomes_Project/chm13v2.0/Phased_SHAPEIT5_v1.1/. Annotation of regulatory elements is downloaded from ENCODE at https://hgdownload.soe.ucsc.edu/goldenPath/hg38/database/wgEncodeReg*. GWAS catalog (version: 2025-09-04) is downloaded from https://www.ebi.ac.uk/gwas/docs/file-downloads. *cis*-eQTL sites are downloaded from GTEx V10 (https://www.gtexportal.org/home/downloads/adult-gtex/qtl). Reference genomes of T2T-CHM13, chimpanzee and bonobo are sourced from NCBI under accession numbers GCF_009914755.1, GCF_028858775.2 and GCF_029289425.2, respectively. Expression data of 1KGP lymphoblastoid cell lines are obtained from Multi-ancestry Analysis of Gene Expression (https://github.com/mccoy-lab/MAGE). Archaic segments detected in 610 human genome assemblies via ASMaid have been deposited on Github (https://github.com/Asian-Pan-Genome/ArchaicIntrogression). Detailed annotations of centromeres are available at https://github.com/Asian-Pan-Genome/Centromere. Putatively archaic-introgression derived structural variants and their annotations are available in the Supplementary Tables.

---

**Extended Data Fig. 1.**
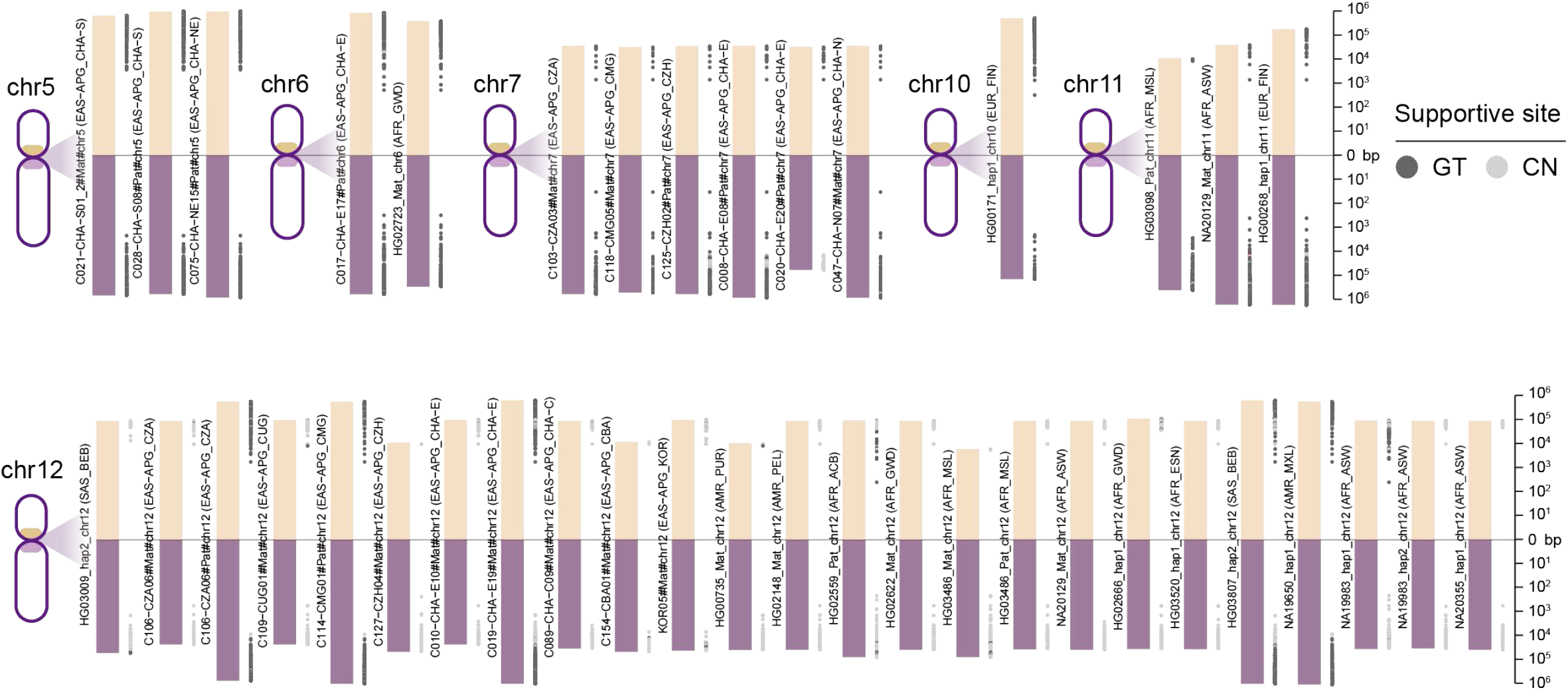
Candidate archaic introgression segments spanning centromeric regions. Genomic profiles are shown for six representative chromosomes harboring centromere-spanning introgression across 40 haploid assemblies. For visualization, centromeres are collapsed into a point (zero). Evidence for archaic introgression is indicated by signals from SNV genotyping (GT, dark gray) and copy number information of structural variations (CN, light gray), plotted along the pericentromeric regions of the short (p) and long (q) arms.

**Extended Data Fig. 2.**
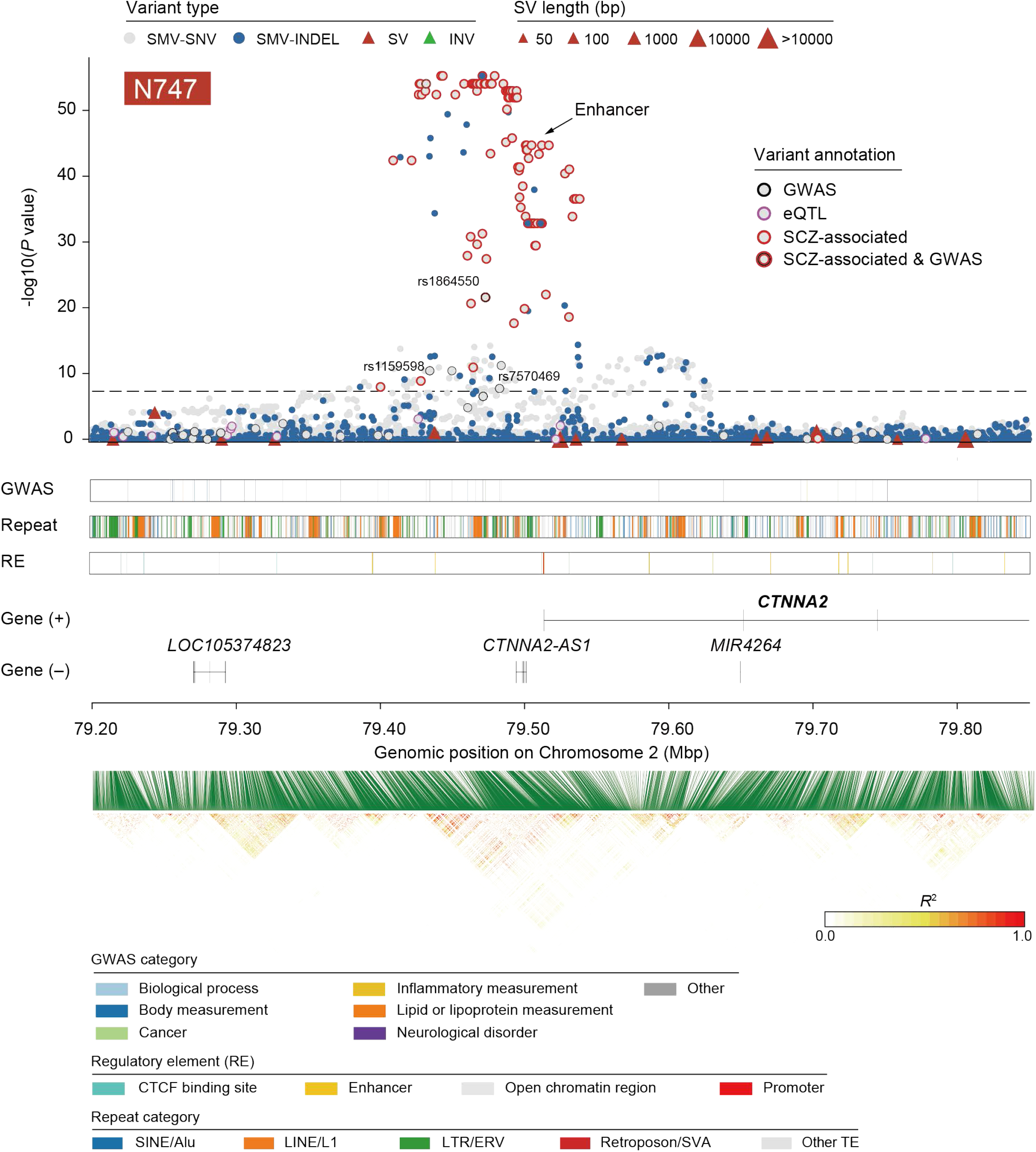
Association of archaic introgression at the *CTNNA2* locus with early-onset schizophrenia (EOSCZ) risk. The top panel displays introgression significance signals (Fisher’s exact test) for each variant, color-coded by variant type and functional annotation, including GWAS, eQTL signals, and SCZ risk alleles highlighted. The middle panel integrates tracks for GWAS signals (colored by trait category), repetitive elements (annotated by Repeatmasker and colored by major repeat class), regulatory elements (from Ensembl 113) and genes. The bottom panel shows the local genetic linkage.

**Extended Data Fig. 3.**
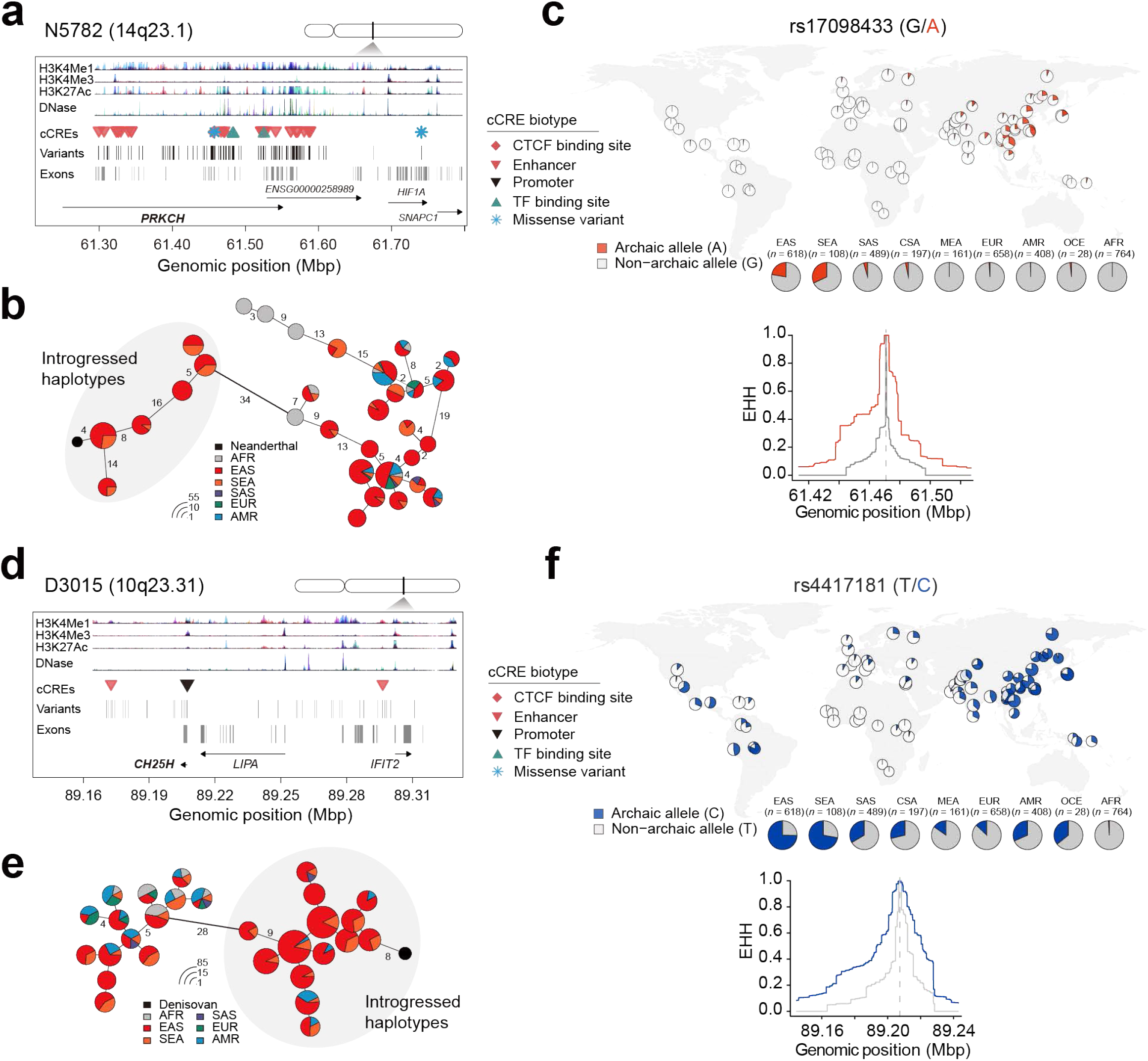
Haplotype analysis of archaic introgression genes *PRKCH* and *CH25H*. **a** and **d**, Genomic architecture and regulatory landscapes of focal archaic chunks. a, Neanderthal chunk N5782 (14q23.1). d, Denisovan chunk D3105 (10q23.31). In both panels, all introgressed variants are marked by vertical black lines, with variants overlapping candidate *cis*-regulatory elements (cCREs) further indicated by distinct symbols according to their predicted regulatory class. Tracks for chromatin accessibility (DNase I hypersensitivity), and histone modification signals (H3K27ac, H3K4me3, H3K4me1) combined from all cell types of ENCODE are shown. **b** and **e**, Haplotype networks of core adaptive introgressed regions. Networks are built from the most frequent haplotypes in the Neanderthal N5782 (b) and Denisovan D3105 (e) chunks. Each node represents a unique haplotype; its radius is proportional to log₂(number of carriers) plus a minimum offset for visibility. Pie sections show the superpopulation composition. Edge widths correspond to the number of pairwise differences between connected haplotypes. **c** and **f**, Geographic distribution and haplotype-based selection signals of archaic-derived variants. c, Neanderthal-derived variant rs17098433; f, Denisovan-derived variant rs4417181. Global allele frequencies are based on the 1,000 Genomes Project (1KGP) and the Human Genome Diversity Project (HGDP) datasets. Extended haplotype homozygosity (EHH) decay curves are shown separately for haplotypes carrying the archaic-derived allele (red/blue) and the non-archaic allele (gray).

## Notes

https://github.com/Asian-Pan-Genome/ArchaicIntrogression

https://github.com/Asian-Pan-Genome/ASMaid

### Competing Interest Statement

The authors have declared no competing interest.

