## Supplementary Figures for "Deciphering complete archaic introgression sequences in modern human genomes"

for

Suo *et al*.

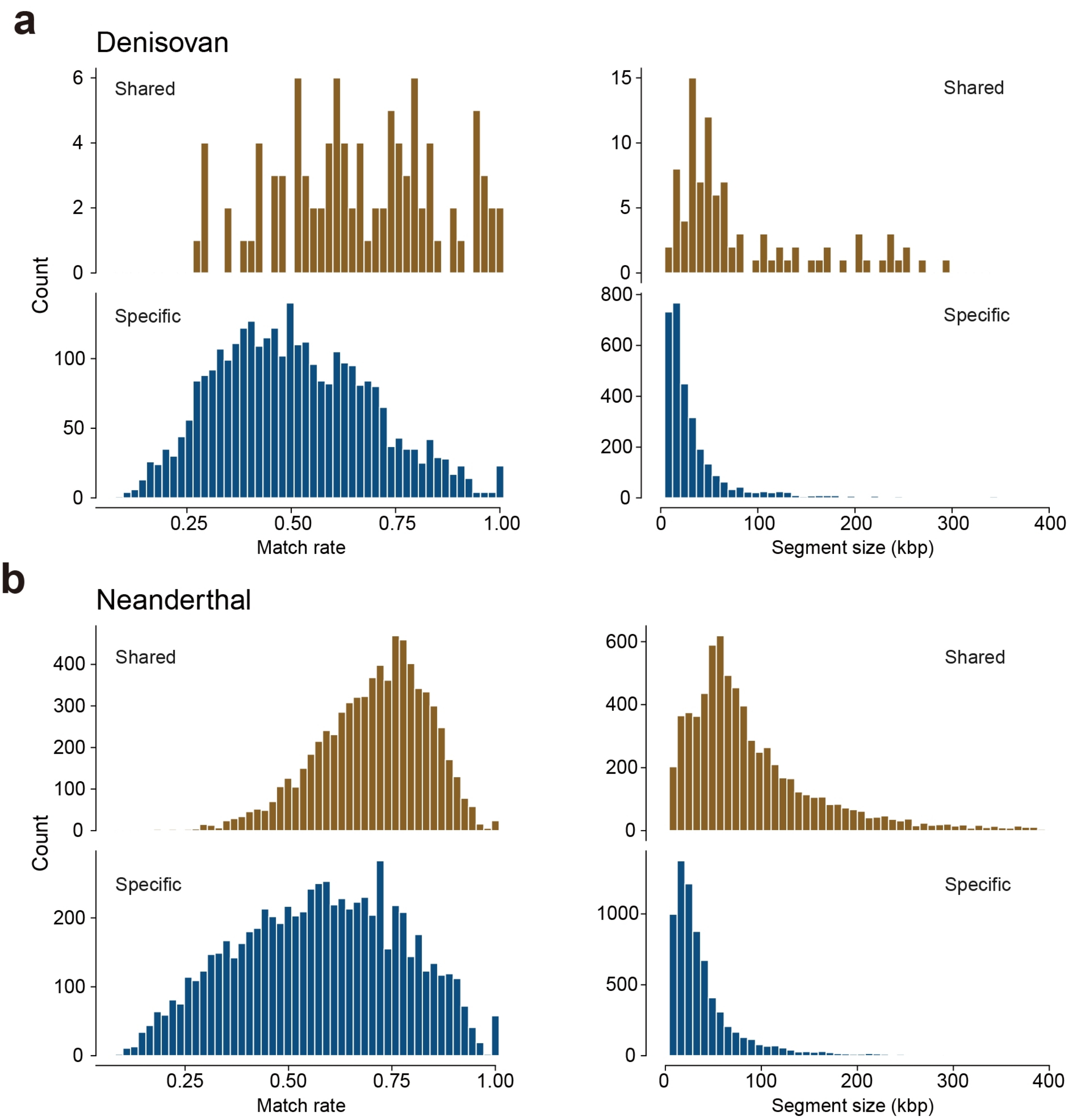


**Supplementary Fig. 1 | Match rate and segment size of archaic introgression sequences identified by ASMaid.** Haploid-level archaic segments (**a**, Denisovan; **b**, Neanderthal) from 15 non-African individuals in the 1KGP cohort are shown. Segments are categorized as either "shared" (identified by ASMaid and at least one established method IBDmix or Sprime) or "specific" to ASMaid. Denisovan archaic segments called by Sprime, and Neanderthal sequences called by IBDmix and Sprime, were obtained from previous studies (Browning *et al*., 2018; Liang *et al*., 2025). To maintain compatibility with individual-level outputs of IBDmix and Sprime, ASMaid-derived introgression segments for two haploid assemblies of each individual are merged for this comparison.

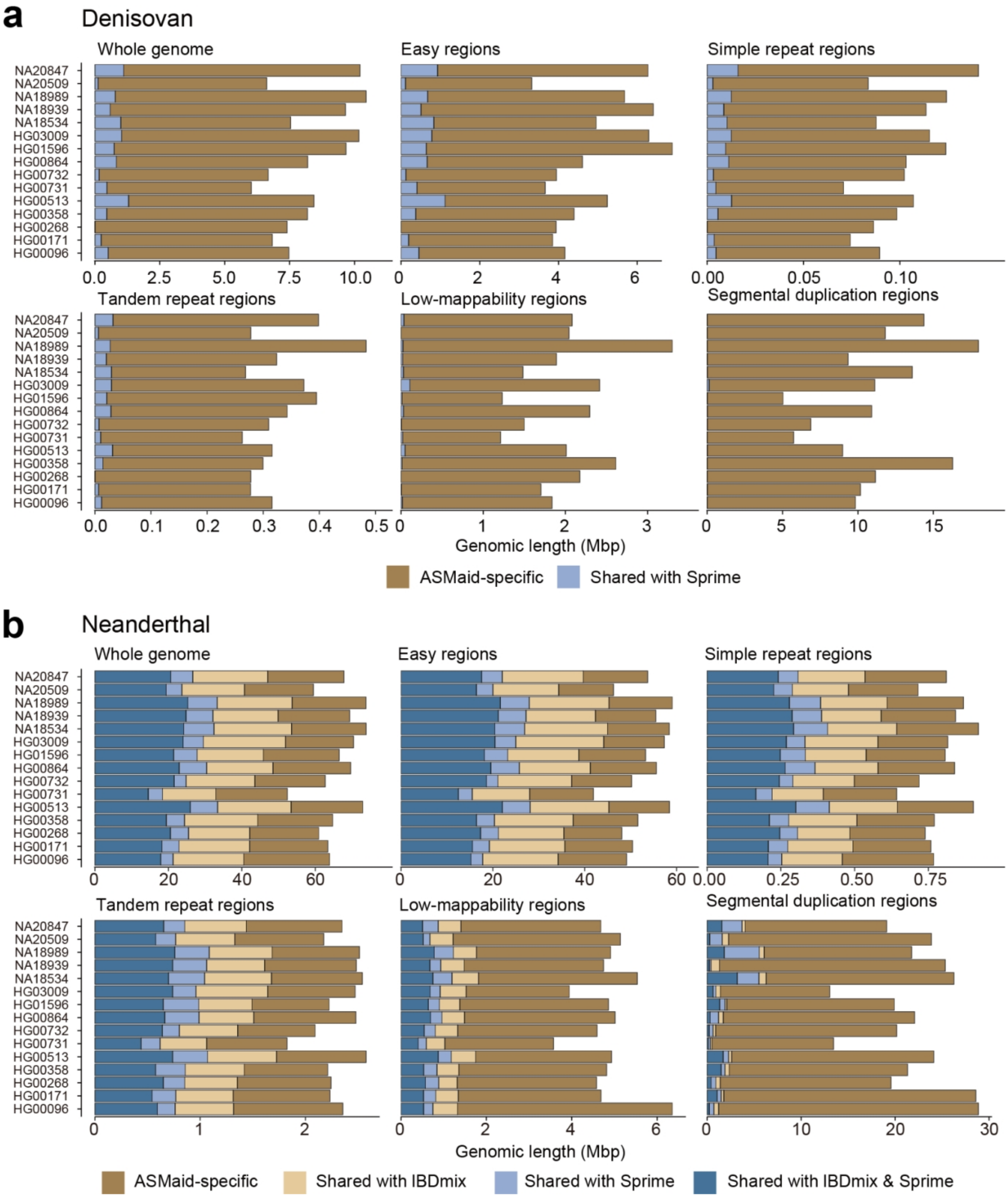


**Supplementary Fig. 2 | Genomic context of archaic introgression segments identified by ASMaid.** The genomic context of identified archaic segments is shown for Denisovan (**a**) and Neanderthal (**b**) across 15 non-African individuals from 1KGP. Five genomic contexts are categorized with different levels of repetitiveness or complexity, including easy-to-map regions (Easy), simple repeat regions, tandem repeat regions, low-mappability regions, and segmental duplications (SD) regions. Segments are further stratified as either shared with previous methods (IBDmix and Sprime) or specific to ASMaid.

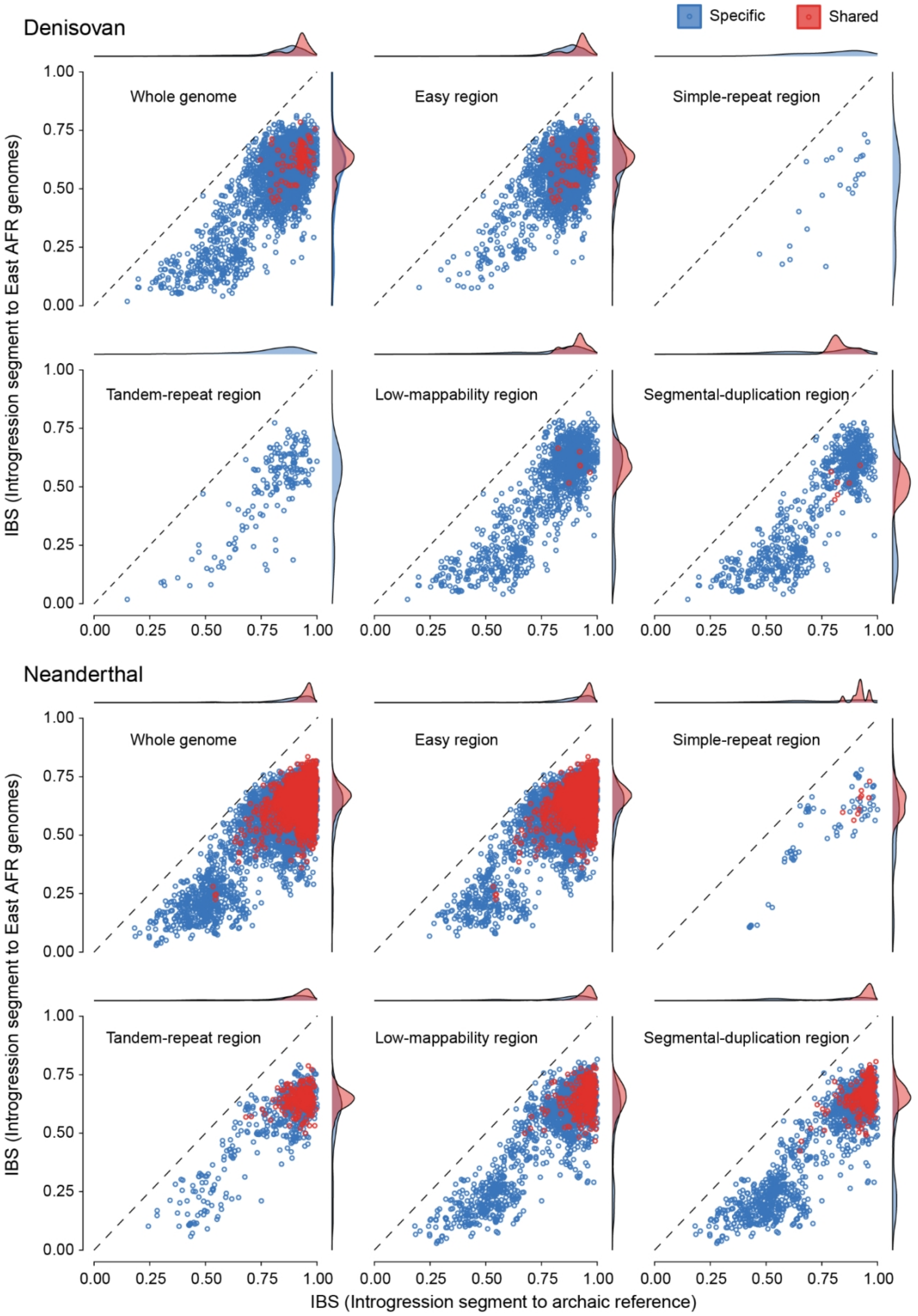


**Supplementary Fig. 3 | Genetic distance of introgression sequences in modern human genomes relative to archaic hominin and East African genomes.** Identity-by-state (IBS) statistics are used to quantify genetic distances. Distances are shown for introgressed segments across the whole genome and five genomic subsets are displayed, with comparisons between ASMaid-specific calls and those shared with Sprime and IBDmix.

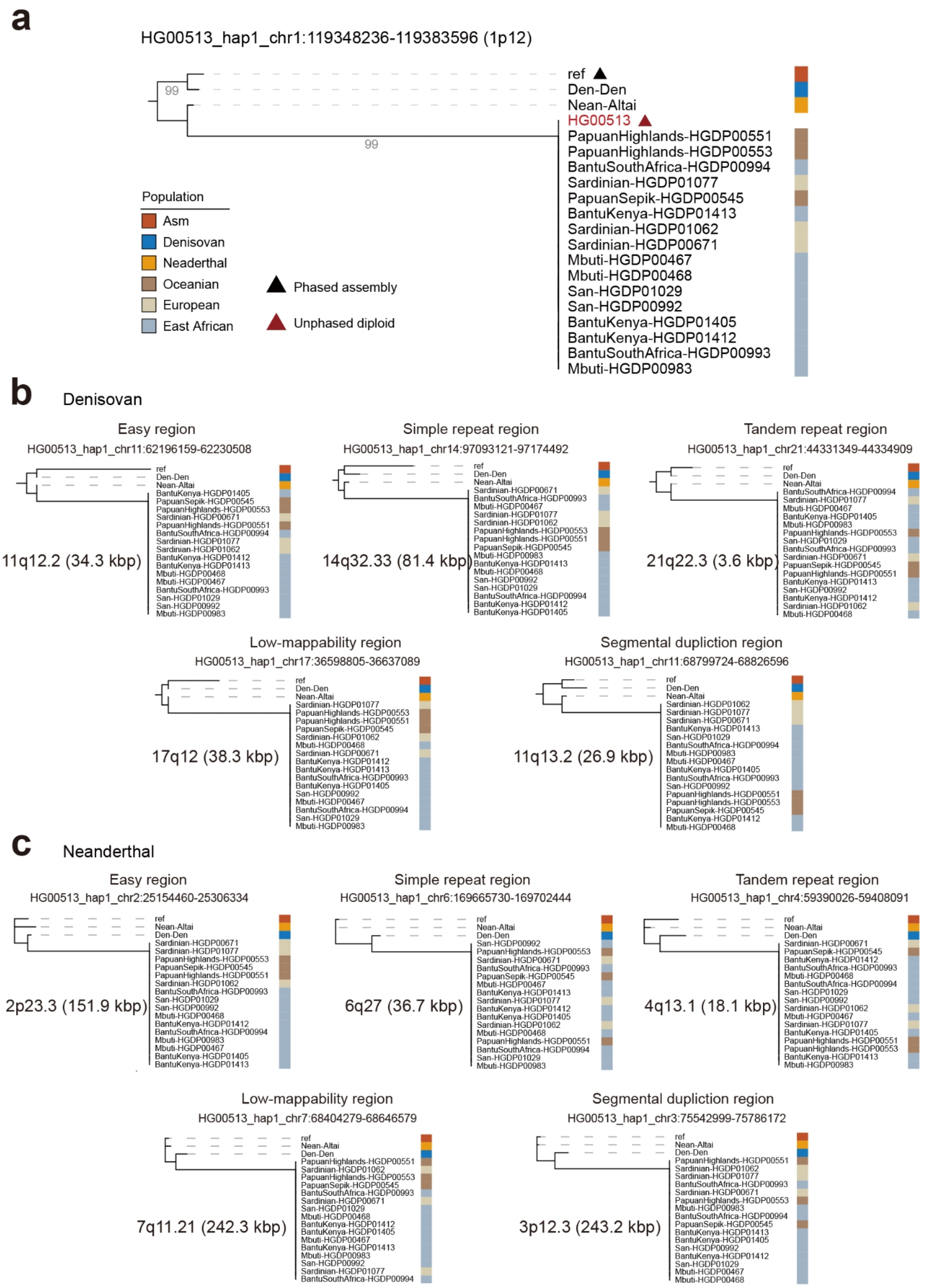


**Supplementary Fig. 4 | Phylogenetic analysis for ASMaid-specifc archaic segments in HG00513. a**, A case at 1p12 demonstrating the enhanced sensitivity of haploid-resolution analysis in ASMaid for capturing additional archaic introgression signals. In the diploid Han

Chinese individual HG00513 from 1KGP, no Denisovan introgression signals at 1p12 was detected by IBDmix or Sprime, aligning with the topological positioning. While in haploid resolution, the hap1 assembly exhibited a clear Denisovan introgression signal at this locus, with sufficient support of 98 introgression sites. **b** and **c**, Phylogenetic trees of ASMaid-specifc calls across varying genomic contexts for Denisovan (**b**) and Neanderthal (**c**), respectively. The “ref” denotes to the haploid assembly (HG00513#hap1), used as the target reference for NGS read mapping.

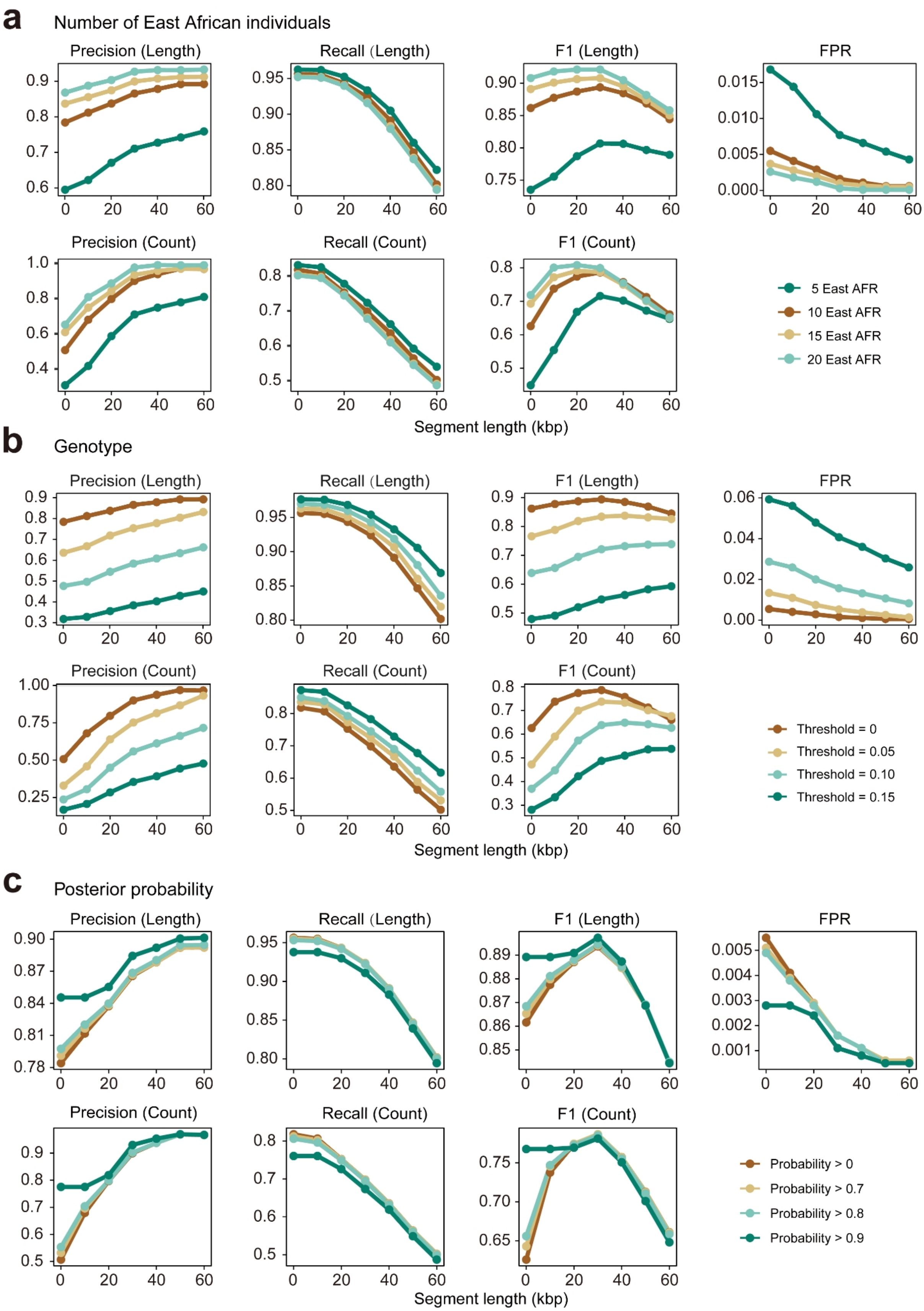


**Supplementary Fig. 5 | Benchmarking of ASMaid performance using simulation data.** Totally 100 non-African genomes (100 Mbp per genome) were simulated using msprime. The evaluation matrix includes precision, recall, F1 score and false positive rate (FPR), calculated based on both segment length and count. The *x*-axis refers to size filtering to ASMaid results by gradient thresholds (0, 10 kbp, 20 kbp, 30 kbp, 40 kbp, 50 kbp and 60 kbp). Three key parameters are assessed, including the number of East African genomes as background control (**a**), the ratio of non-supportive genotype frequency (**b**) and posterior probability for segment filtration (**c**).

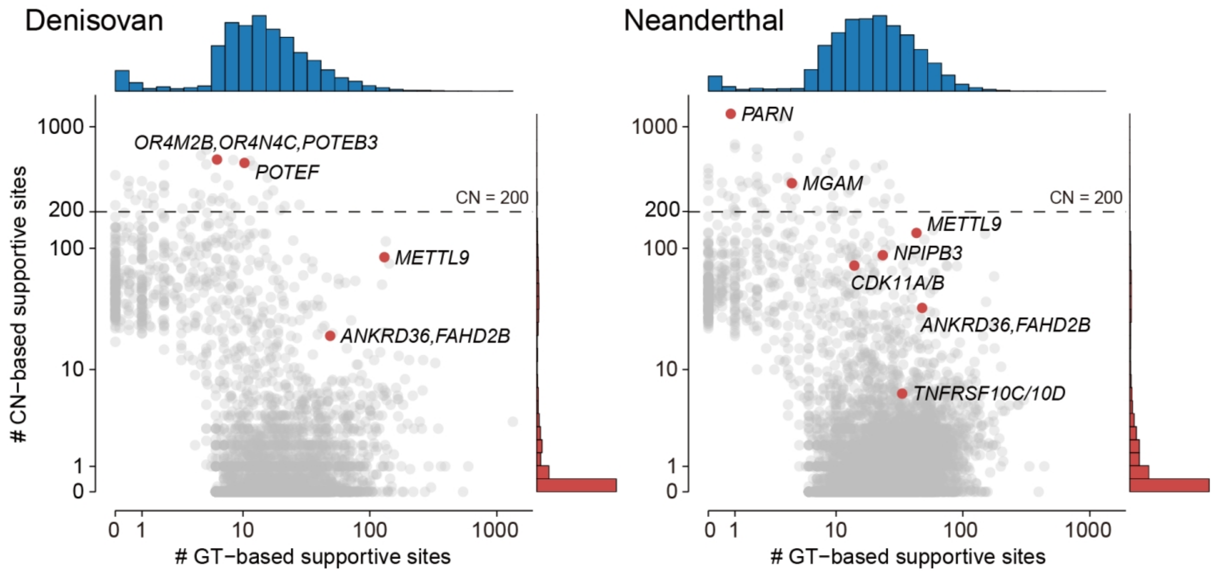


**Supplementary Fig. 6 | Supportive sites within identified archaic introgression chunks.** The average number of supportive sites per introgression chunk is shown, derived from genotype (GT) and copy number (CN) information. The horizontal dashed line represents a threshold of 200 CN-based supportive sites per introgressed chunks. Some genes are highlighted.

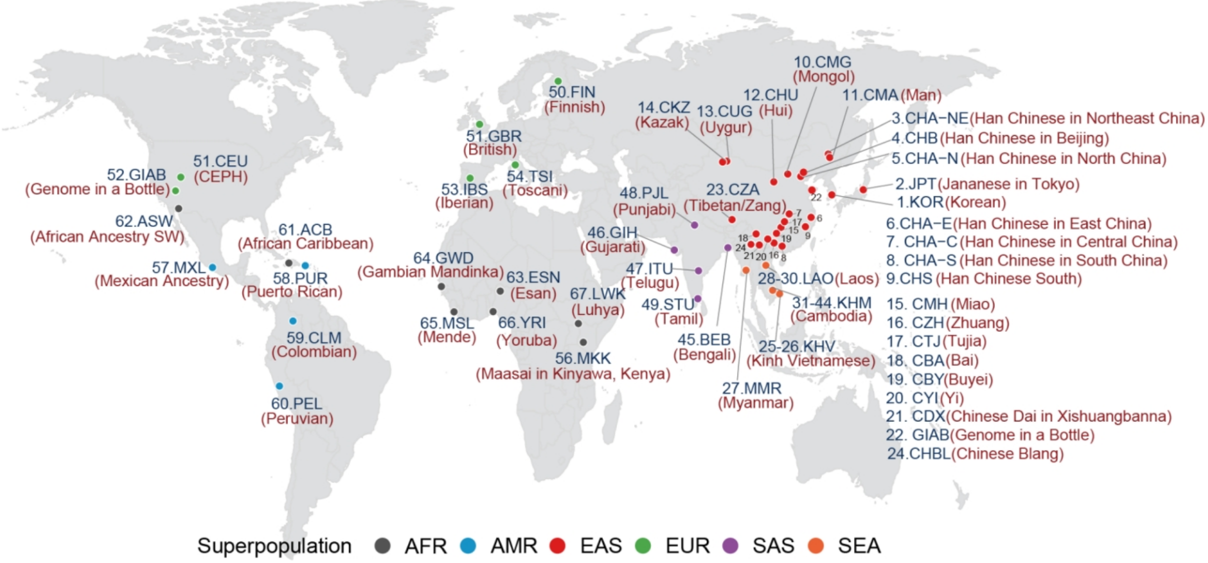


**Supplementary Fig. 7 | Geographic distribution of the global haploid human genome assemblies analyzed for archaic introgression.** The haploid assemblies included in this analysis are sourced from the pangenome projects: the Human Pangenome Reference Consortium year 1 (HPRCy1), the Human Genome Structural Variation Consortium phase 3 (HGSVC3), Asian Pan-Genome project phase 1 (APGp1) and SEA3K from the Consortium of Anthropological Research in Southeast Asia and Southwest China (CASEAC).

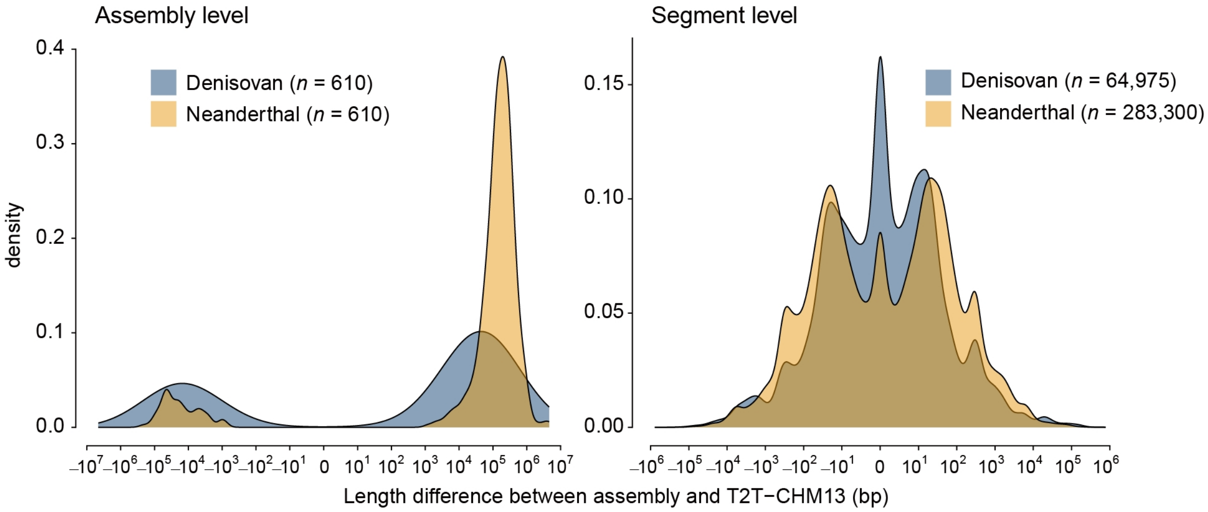


**Supplementary Fig. 8 | Comparison of archaic introgression segment lengths between self-assembly and T2T-CHM13 reference coordinates.** Distribution of length differences for archaic segments when measured in their native self-assembly coordinates *versus* their projected coordinates on the T2T-CHM13 reference. Positive values indicate segments that are larger in self-assembly coordinates.

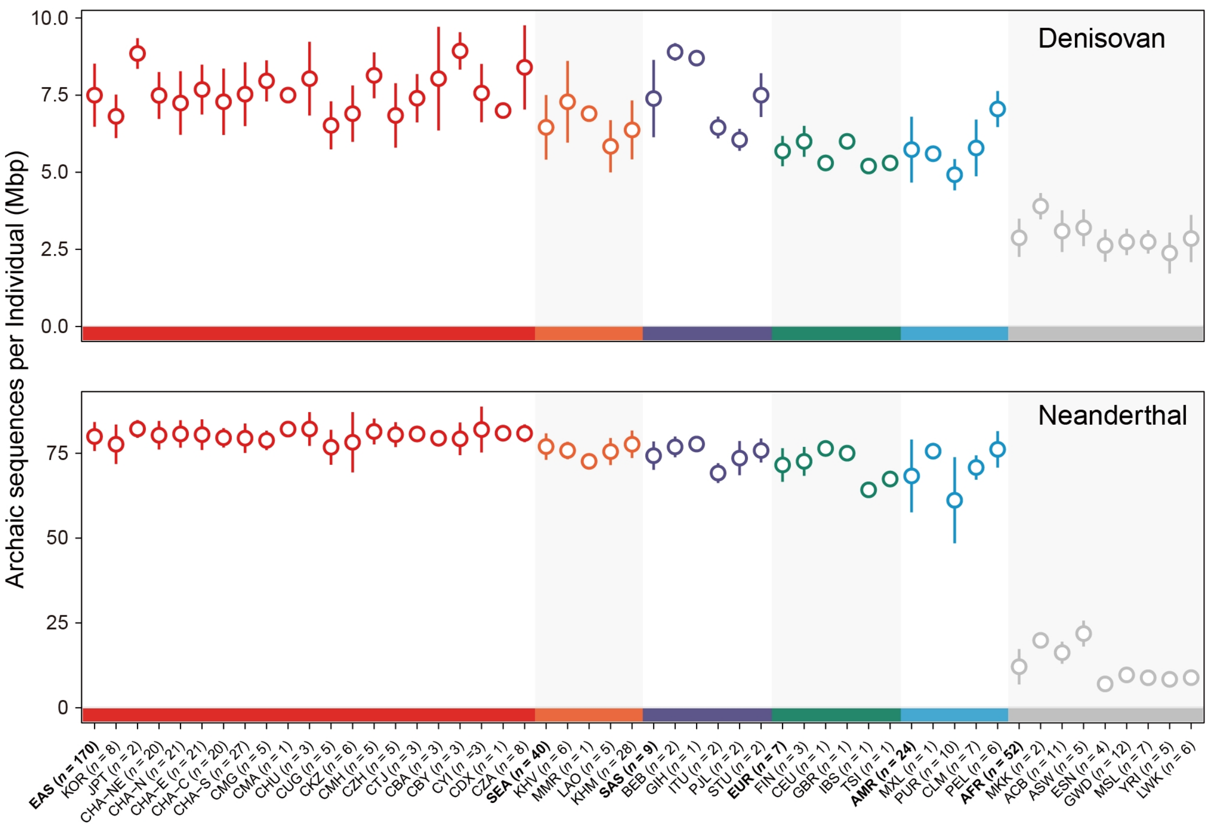


**Supplementary Fig. 9 | Archaic introgression segments per individual across global populations.** Total genomic lengths of introgressed archaic sequences are summarized at the diploid level by aggregating the contributions from both haplotypes for each individual. Populations are grouped by superpopulation and ordered as follows: East Asian (EAS), Southeast Asian (SEA), South Asian (SAS), European (EUR), American (AMR), and African (AFR).

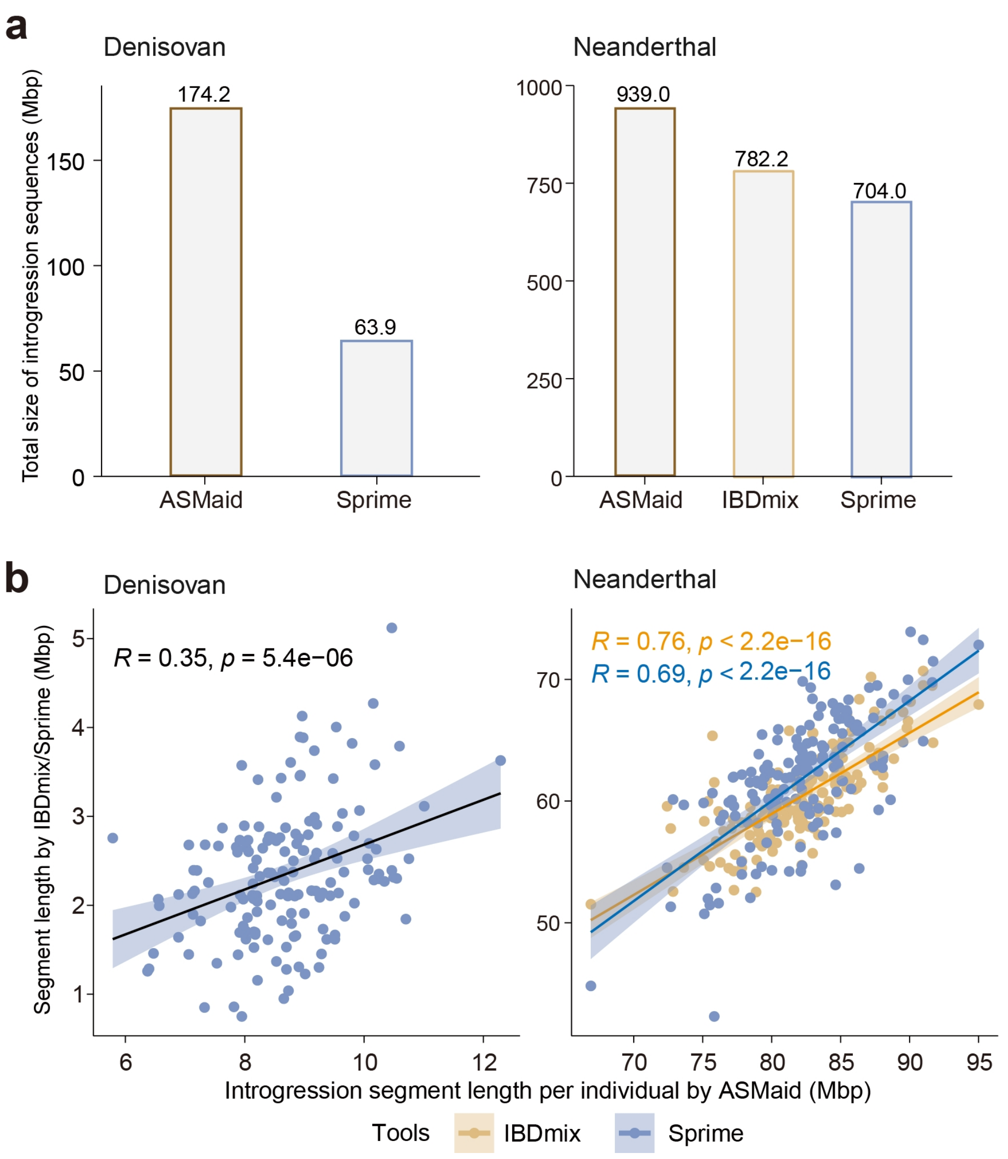


**Supplementary Fig. 10 | Comparison of archaic introgression segments in APGp1 modern human genomes called by ASMaid, IBDmix and Sprime. a**, Non-redundant archaic introgression segments identified across APGp1 individuals by each method. **b**, Individual-level comparison and correlation between ASMaid and IBDmix/Sprime calls. Pearson's correlation, *R* and *P* values are shown in each panel.

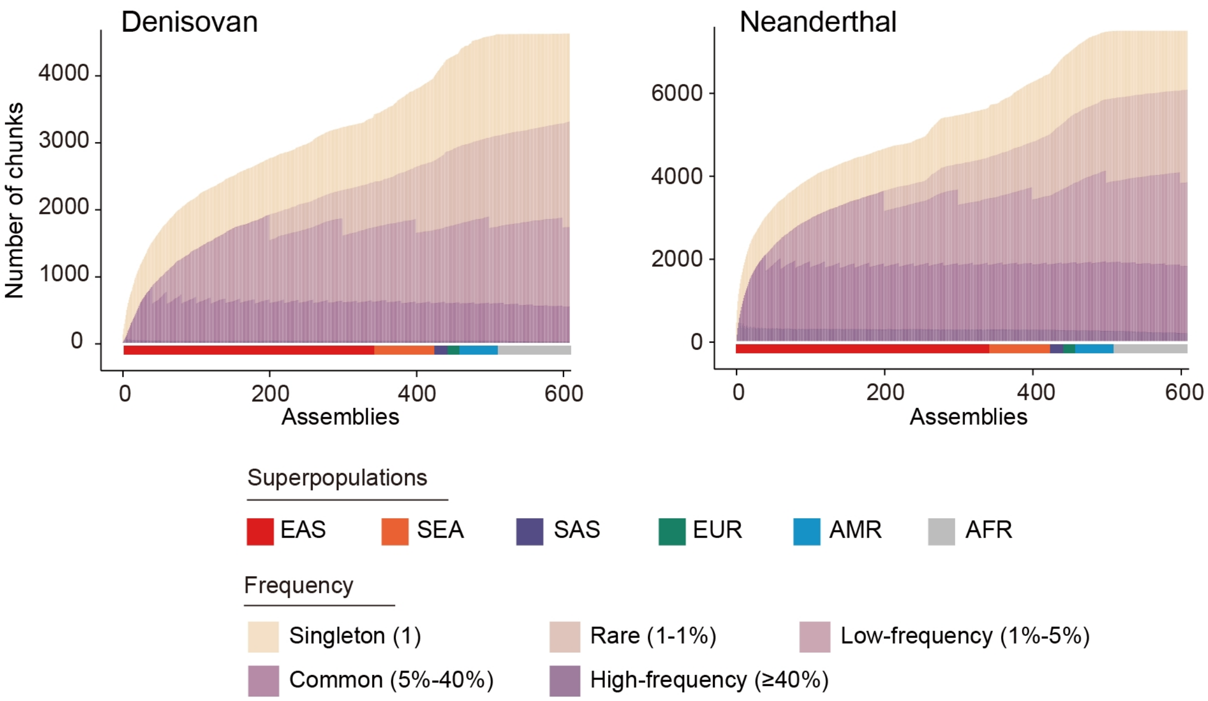


**Supplementary Fig. 11 | Cumulative curves of introgressed chunk counts in modern human genomes.** The non-redundant archaic sequence chunks for Denisovan (left) and Neanderthal (right) are cumulatively added by superpopulation in the following order: EAS, SEA, SAS, EUR, AMR and AFR. The frequencies are calculated based on the current involved sample size.

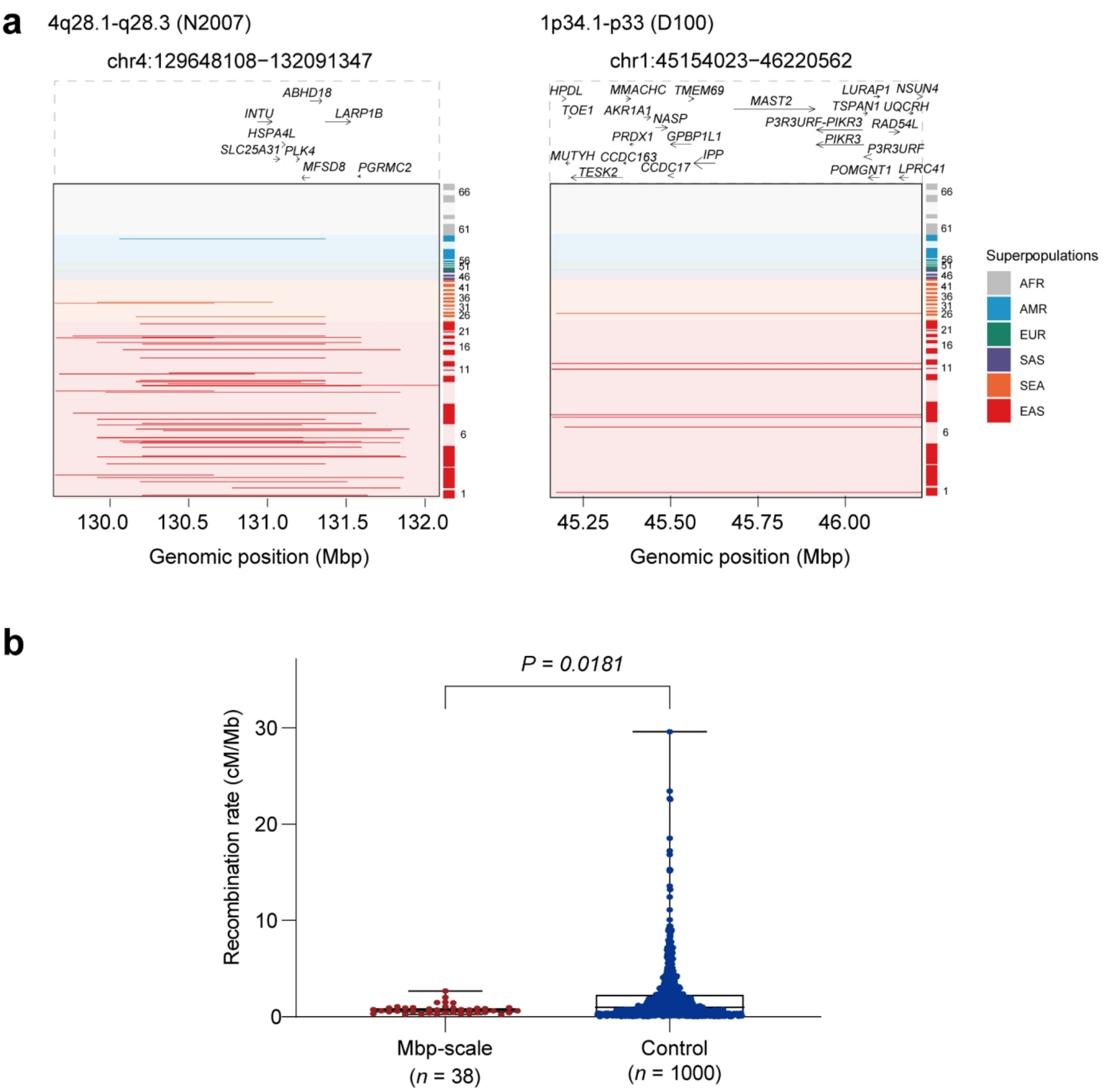


**Supplementary Fig. 12 | Characterization of ultra-long archaic segments (>1 Mbp) in modern human genomes**. **a**, Genomic landscape of ultra-long introgressed haplotype blocks derived from Neanderthal and Denisovan. Archaic-derived segments are highlighted in dark red (EAS), orange (SEA), green (EUR), blue (AMR) or gray (AFR). **b**, Comparison of local recombination rates between ultra-long (>1 Mbp) and short archaic introgressed fragments (<100 kbp, 1000 segments randomly selected). The centerline of each box plot indicates the median and the lower and upper hinges indicate the 25th and 75th percentiles, respectively, and the whiskers extend to the minimum and maximum values.

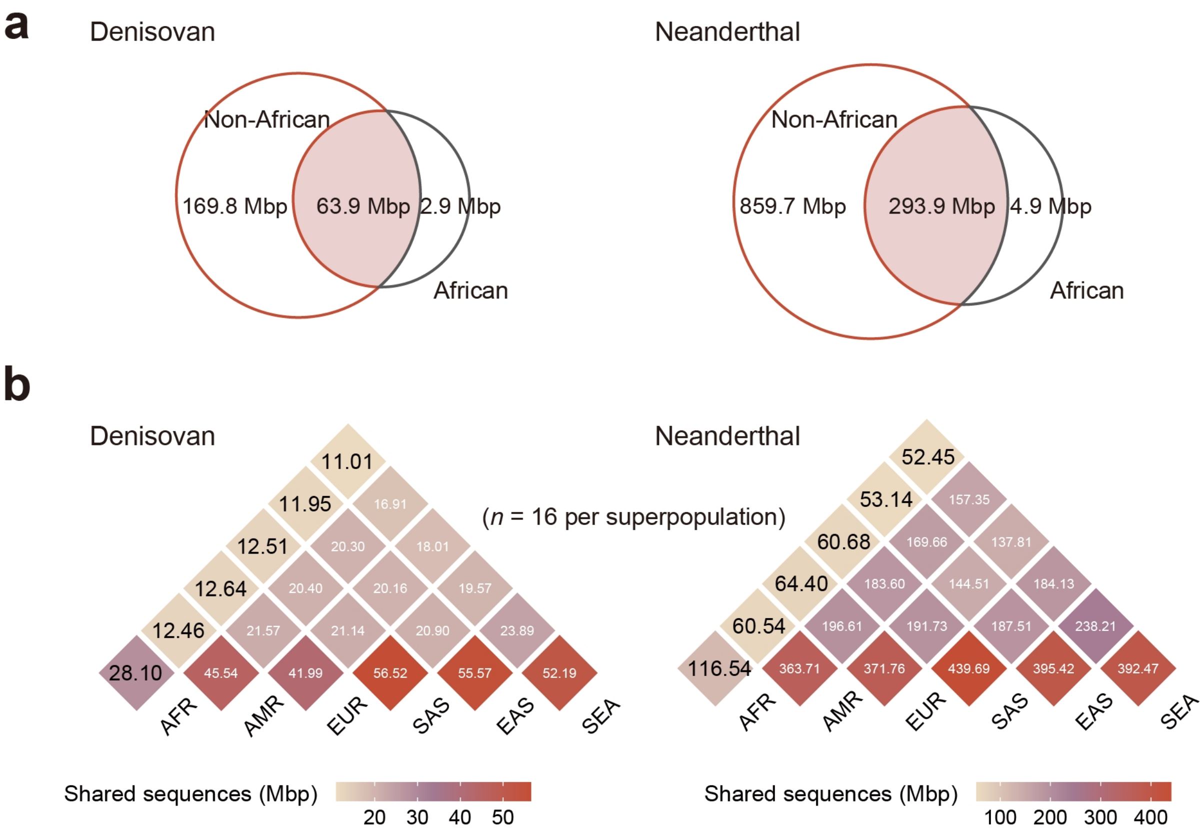


**Supplementary Fig. 13 | Archaic introgression segments in African genomes. a**, Comparative overlap of archaic sequences between African and non-African superpopulations. Venn diagrams show the shared and unique sequences for Denisovan (left) and Neanderthal (right) introgression between African and non-African superpopulations. **b**, Intersection of merged introgression segments across all analyzed superpopulations. To ensure comparability across superpopulations, we randomly downsample 16 assemblies from each superpopulation to accommodate those with limited sample sizes.

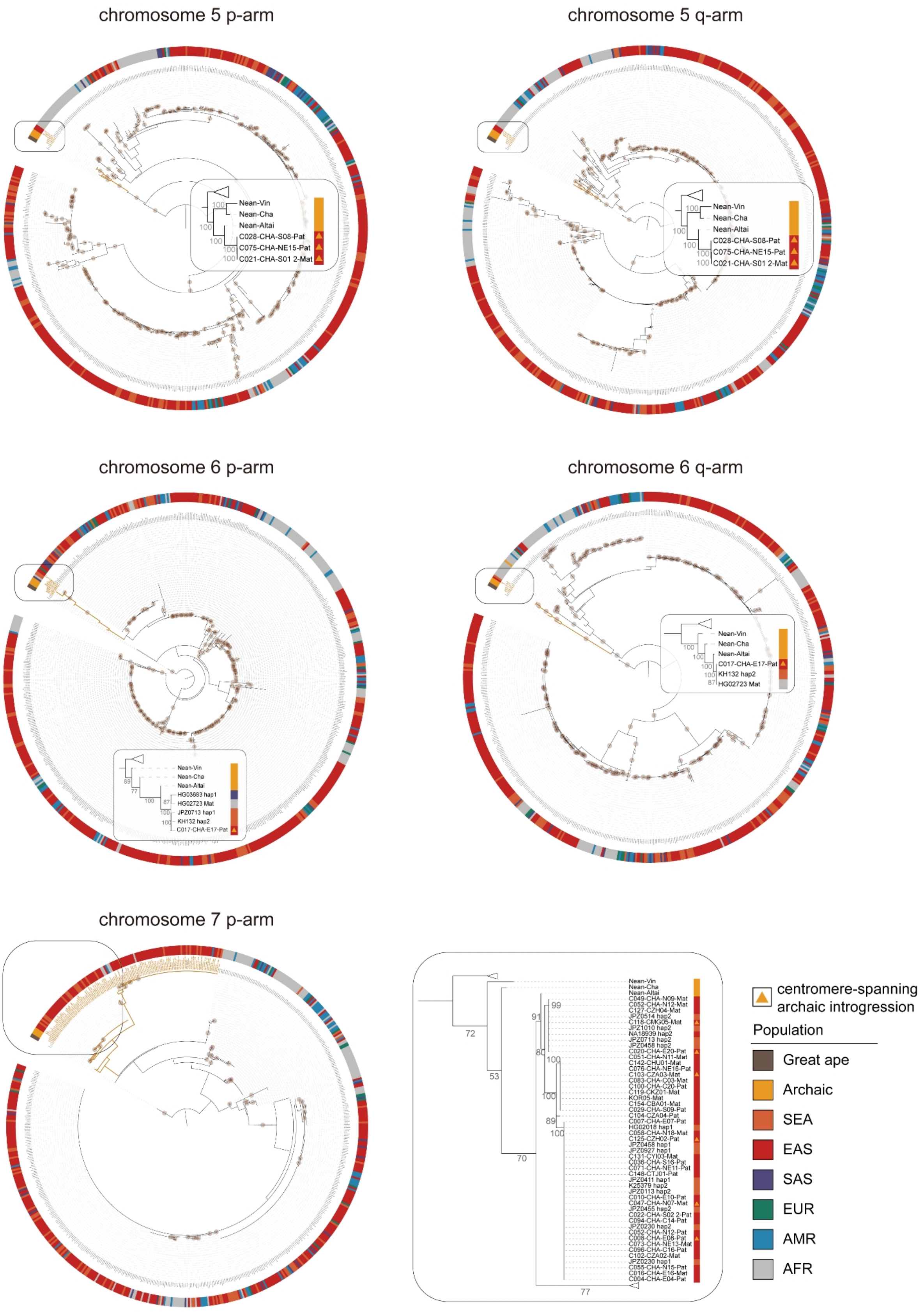
chromosome 5 p-arm
Nean-Vin
Nean-Cha
Nean-Altai
C028-CHA-S08-Pat
C075-CHA-NE15-Pat
C021-CHA-S01 2-Mat
chromosome 5 q-arm
Nean-Vin
Nean-Cha
Nean-Altai
C028-CHA-S08-Pat
C075-CHA-NE15-Pat
C021-CHA-S01 2-Mat
chromosome 6 p-arm
Nean-Vin
Nean-Cha
Nean-Altai
HG03683 hap1
HG02723 Mat
JPZ0713 hap1
KH132 hap2
C017-CHA-E17-Pat
chromosome 6 q-arm
Nean-Vin
Nean-Cha
Nean-Altai
C017-CHA-E17-Pat
KH132 hap2
HG02723 Mat
chromosome 7 p-arm
Nean-Vin
Nean-Cha
Nean-Altai
C049-CHA-N09-Mat
C052-CHA-N12-Mat
C127-CZH04-Mat
JPZ0514 hap2
C118-CMG05-Mat
JPZ1010 hap2
NA18939 hap2
JPZ0713 hap2
JPZ0458 hap2
C020-CHA-E20-Pat
C051-CHA-N11-Mat
C142-CHU01-Mat
C076-CHA-NE16-Pat
C103-CZA03-Mat
C083-CHA-C03-Mat
C100-CHA-C20-Pat
C119-CKZ01-Mat
KOR05-Mat
C154-CBA01-Mat
C029-CHA-S09-Pat
C104-CZA04-Pat
C007-CHA-E07-Pat
HG02018 hap1
C058-CHA-N18-Mat
C125-CZH02-Pat
JPZ0458 hap1
JPZ0927 hap1
C131-CYI03-Mat
C036-CHA-S16-Pat
C071-CHA-NE11-Pat
C148-CTJ01-Pat
JPZ0411 hap1
K25379 hap2
JPZ0113 hap2
C010-CHA-E10-Pat
C047-CHA-N07-Mat
JPZ0455 hap2
C022-CHA-S02 2-Pat
C094-CHA-C14-Pat
JPZ0230 hap2
C052-CHA-N12-Pat
C008-CHA-E08-Pat
C073-CHA-NE13-Mat
C096-CHA-C16-Pat
C102-CZA02-Mat
JPZ0230 hap1
C055-CHA-N15-Pat
C016-CHA-E16-Mat
C004-CHA-E04-Pat
centromere-spanning archaic introgression
Population
Great ape
Archaic
SEA
EAS
SAS
EUR
AMR
AFR

chromosome 10 p-arm

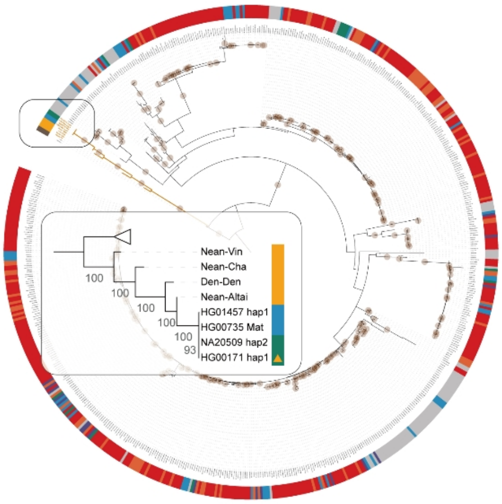


chromosome 10 q-arm

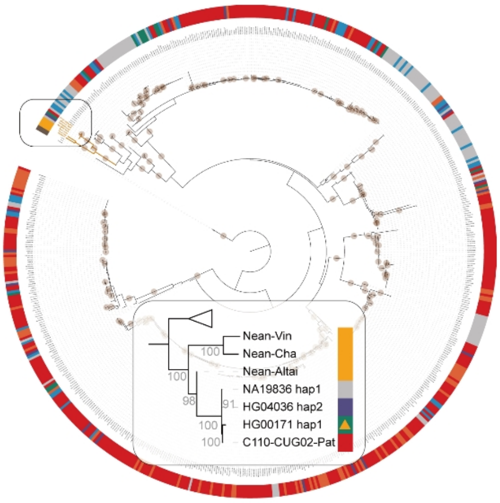


chromosome 11 p-arm

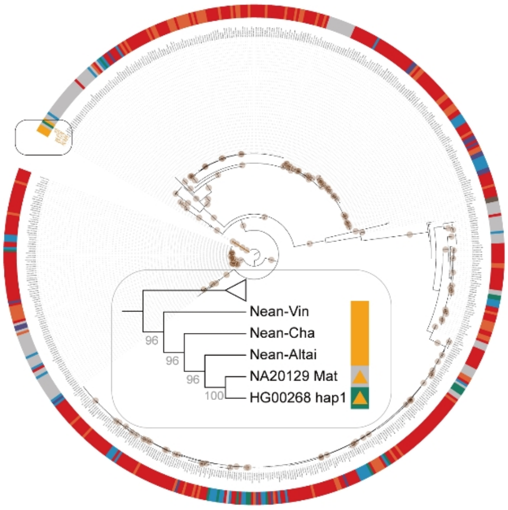


chromosome 11 q-arm

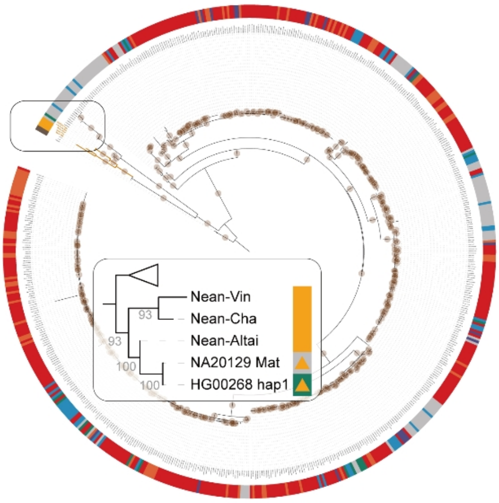


chromosome 12 p-arm

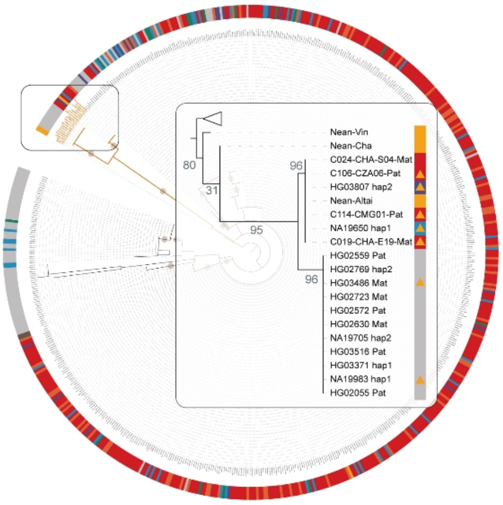


chromosome 12 q-arm

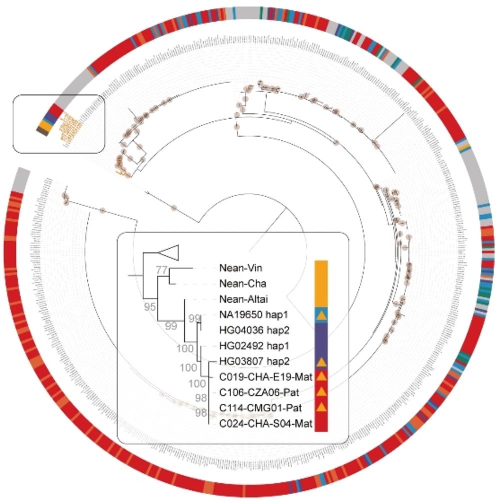

**Supplementary Fig. 14 | Phylogenetic evidence of candidate archaic introgression in centromeric regions.** Maximum-likelihood phylogenetic trees are constructed for homologous peri-centromeric regions identified as harboring putative archaic introgression signals. The q arm from chromosome 7 is excluded from this analysis due to their poor alignment quality during coordinate liftover from individual assemblies to the T2T-CHM13 reference. The boxes highlight the introgressed clades, with detected introgressed haploid assemblies highlighted by yellow triangles. Nodes with bootstrap support values greater than 95 are indicated by brown circles.

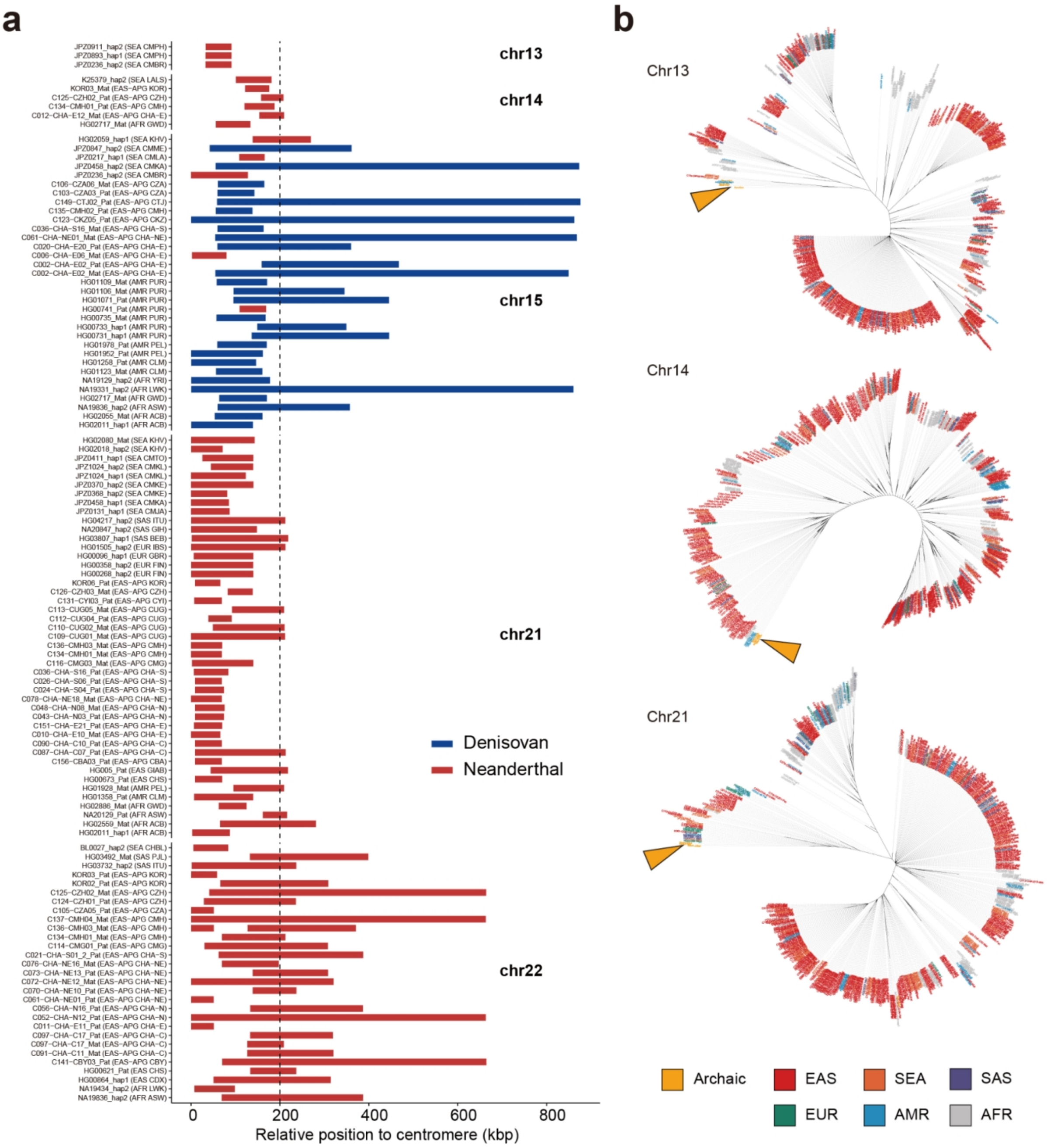


**Supplementary Fig. 15 | Archaic introgression landscape across acrocentric centromeres. a**, Distribution and frequency of introgressed segments within the 200-kbp pericentromeric regions on the q arms of five acrocentric chromosomes. The *x*-axis origin (*x* = 0) denotes the distal boundary of the centromere for each chromosome. **b**, Unrooted phylogenetic trees constructed from 100-kbp q-arm pericentromeric sequences. The q arms on chromosomes 15 and 22 are excluded from this analysis due to their poor alignment quality during coordinate liftover from individual assemblies to the T2T-CHM13 reference. Yellow triangles denote the phylogenetic positions of archaic hominins.

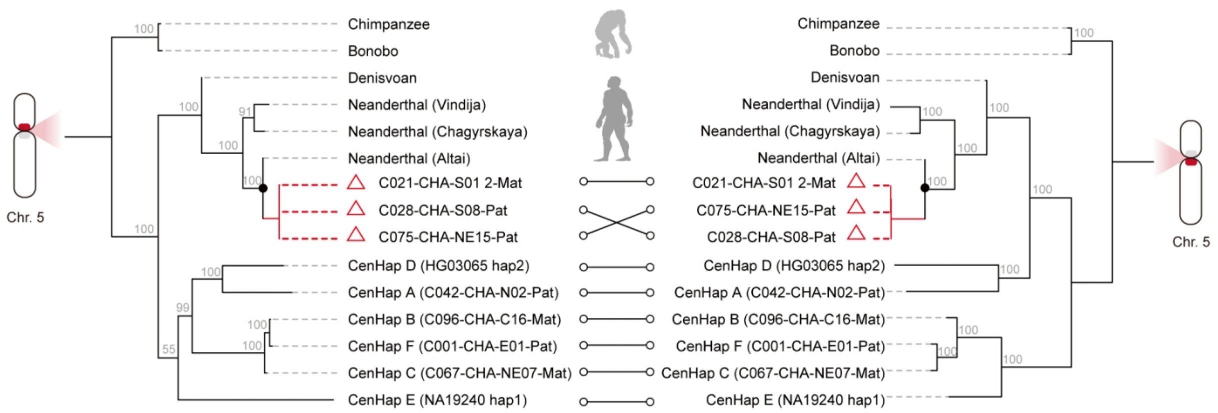


**Supplementary Fig. 16 | Phylogeny and divergence of human centromeres on chromosome 5.** Phylogenetic trees are constructed using ~600-kbp flanking sequences (p and q arm) of chromosome 5 centromere. Red triangles denote the three specific haploid assemblies carrying centromere-spanning introgression signals. Bootstrap support values in 1,000 replicates are indicated in gray at each node.

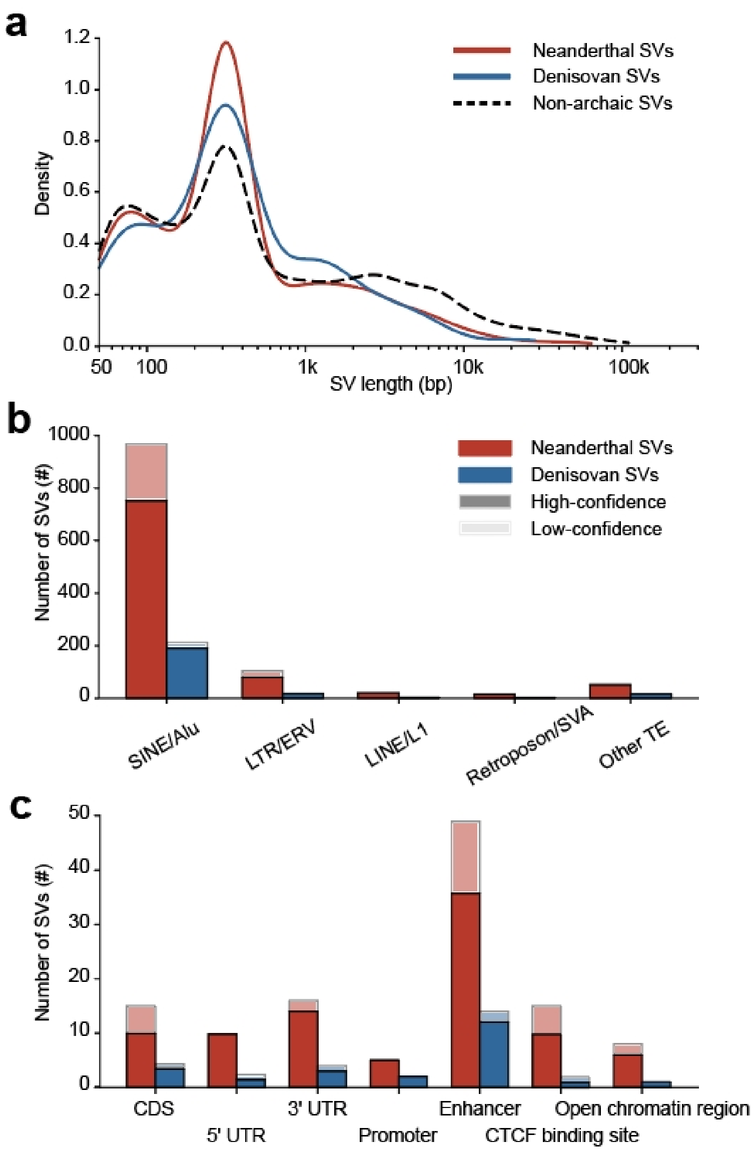


**Supplementary Fig. 17 | Genomic characterization of structural variants (SVs) from archaic introgression. a**, Density distribution of SV lengths for archaic-derived and non-archaic SVs, demonstrating concordant length spectra between the two classes. **b**, Counts of Neanderthal- and Denisovan-derived SVs intersecting major classes of repetitive elements. High-confidence SVs denote those validated through direct genotyping of archaic genomes. **c**, Functional annotation of archaic-derived SVs across various genomic categories. Variants are categorized by their overlap with specific genomic features.

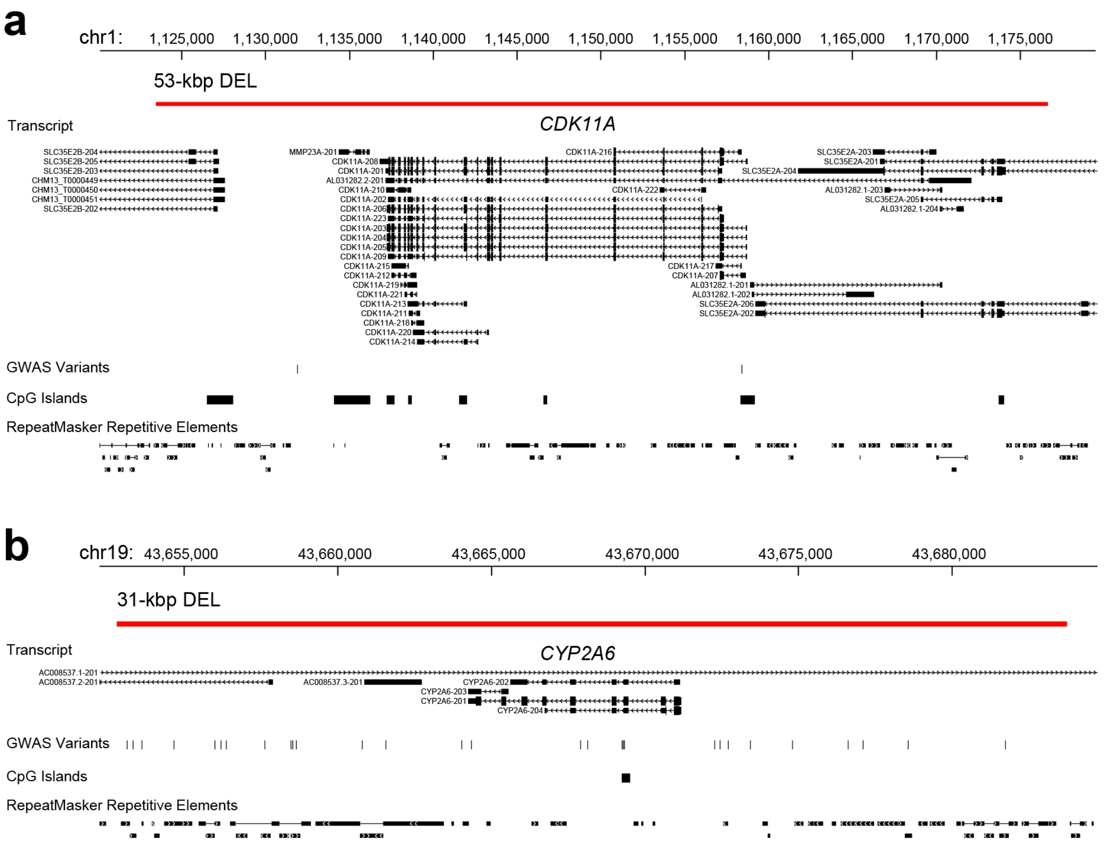


**Supplementary Fig. 18 | Large archaic-derived deletions directly interrupting protein-coding sequences. a**, A ~53-kbp Neanderthal-derived deletion resulting in the complete loss of *CDK11A*. **b**, A ~31-kbp Neanderthal-derived deletion disrupting *CYP2A6*.

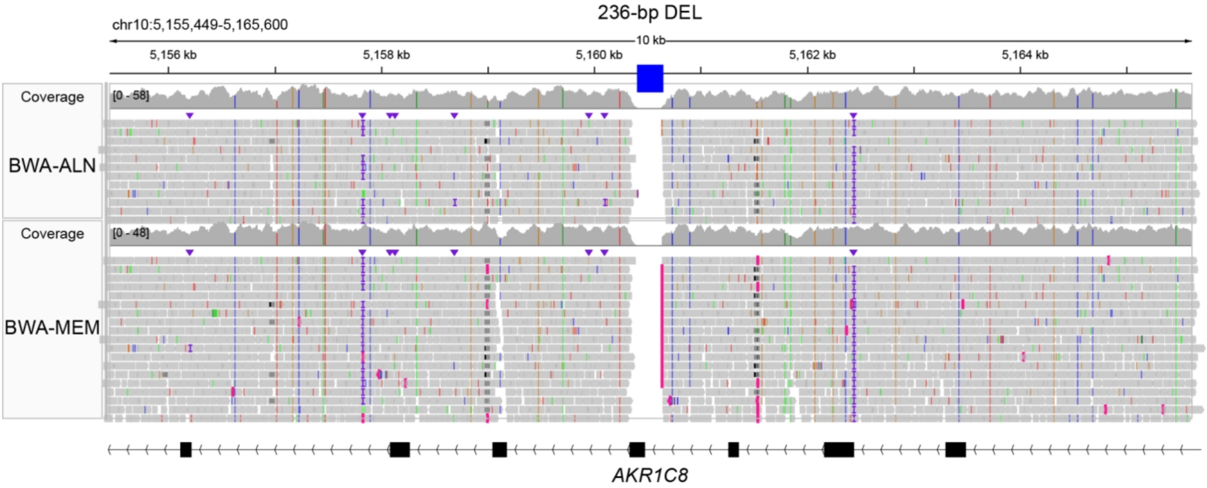


**Supplementary Fig. 19 | High-coverage read mapping validation of a Neanderthal-derived deletion in *AKR1C8*.** IGV (Integrative Genomics Viewer) screenshot displays the alignment of high-coverage raw sequencing reads from the Altai Neanderthal genome to the T2T-CHM13 reference at the *AKR1C8* locus. A 236-bp deletion is clearly evidenced by the absence of coverage, supporting the Neanderthal origin of this structural variant in modern humans. The mapping coverage and alignments generated by BWA ALN and BWA MEM are shown, respectively.

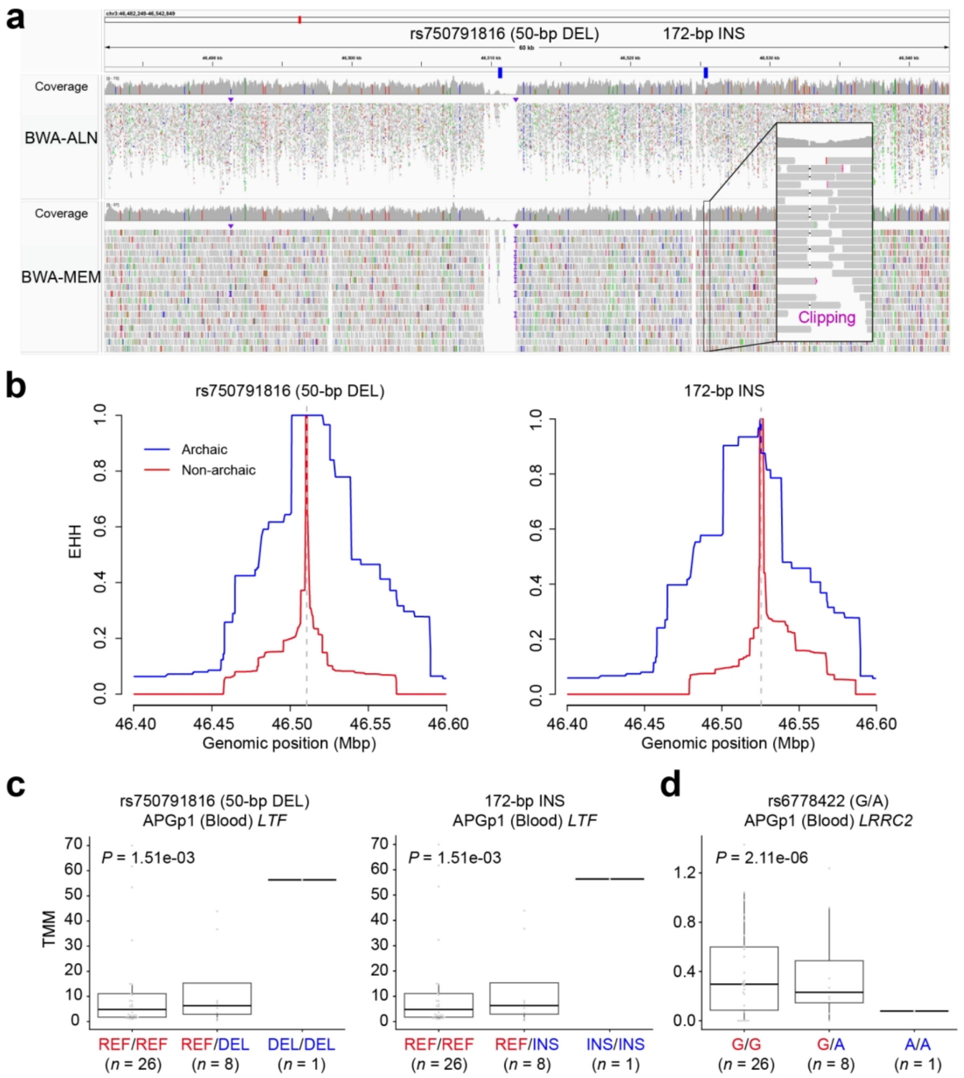


**Supplementary Fig. 20 | Characterization of Neanderthal-derived SVs at the 3p21.31 locus. a**, IGV screenshot of read mapping from the Altai Neanderthal genome to the T2T-CHM13 reference, validating two Neanderthal-derived SVs: a 50-bp deletion (rs750791816) and a 172-bp insertion. Clipping signals are highlighted as evidence of the insertion event. Mapping profiles generated by BWA ALN and BWA MEM are shown, respectively. **b**, Extended Haplotype Homozygosity (EHH) decay analysis for the two pAID-SVs. The blue and red lines represent archaic and non-archaic alleles, respectively. **c**, Association between introgressed SV alleles and *LTF* expression in blood samples from APGp1 individuals. The two-tailed *P* values for the *t*-statistics are derived from a linear regression model estimated using ordinary least squares. **d**, The introgression SV-linked pAID-SNV (rs6778422) acts as a *cis*-eQTL downregulating the expression of *LRRC2* in blood. The centerline of each box plot indicates the median and the lower and upper hinges indicate the 25th and 75th percentiles, respectively. The vertical line of each boxplot extends to 1.5× the interquartile range from each hinge.

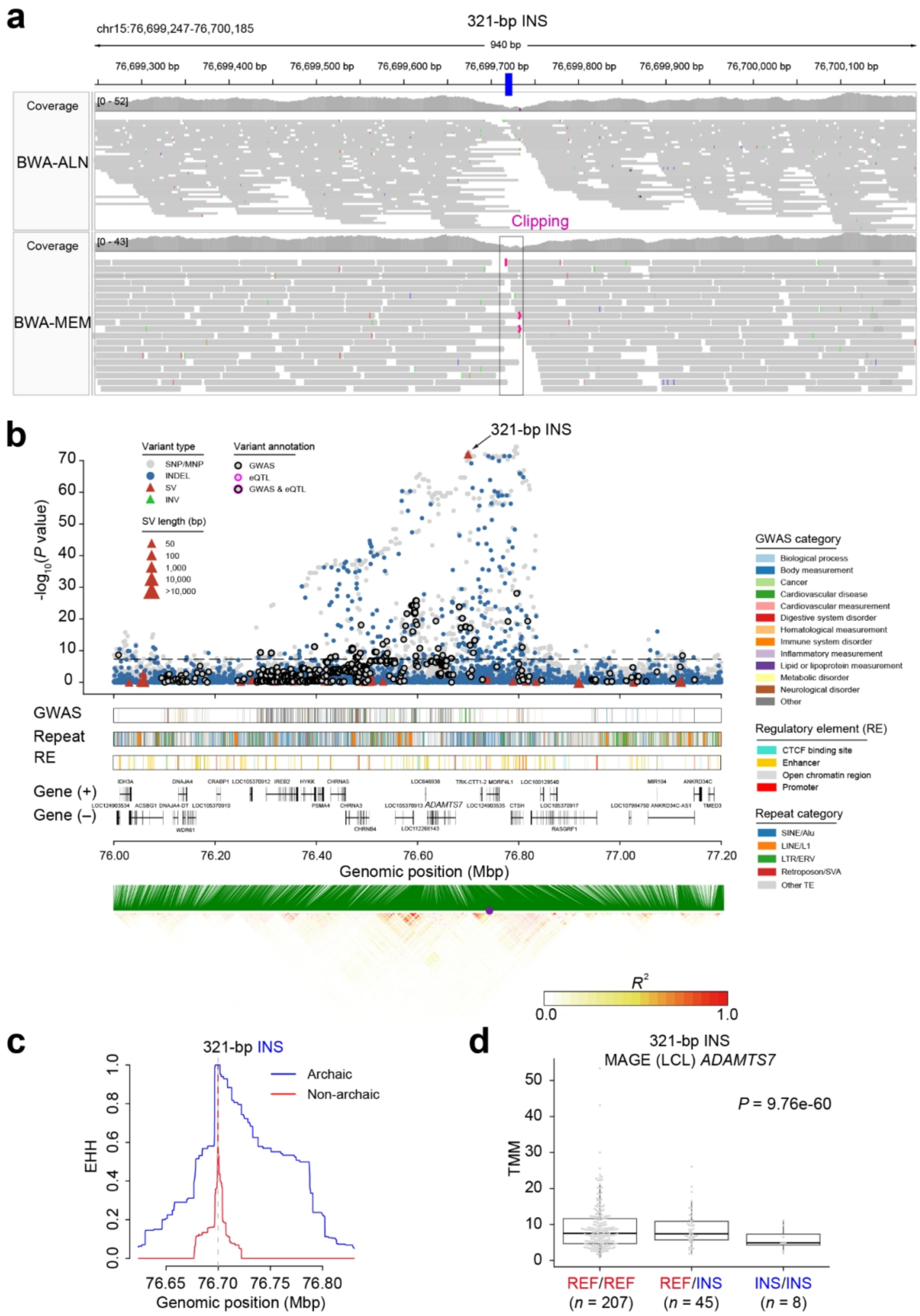


**Supplementary Fig. 21 | Characterization of a Neanderthal-derived insertion at locus 15q25.1. a**, IGV screenshot of read mapping from Altai Neanderthal genome to the T2T-CHM13 reference, validating a 321-bp Neanderthal-derived insertion. Clipping signals are highlighted as evidence of the insertion event. Mapping profiles by BWA ALN and BWA MEM are shown, respectively. **b**, Genomic views of the 15q25.1 locus. The top panel displays the association between introgression and allele frequencies using Fisher's exact test for each variant, colored by variant type and functional annotation, including GWAS and eQTL signals. The middle panel integrates tracks for GWAS signals (colored by trait categories), repetitive elements (annotated by Repeatmasker), regulatory elements (Ensembl 113), and gene models (Refseq). The bottom panel shows a linkage disequilibrium heatmap, with the 321-bp insertion indicated by a purple dot. **c**, Extended Haplotype Homozygosity (EHH) decay analysis for the 321-bp insertion. The blue and red lines represent archaic and non-archaic alleles, respectively. **d**, Association between the 321-bp insertion and downregulated *ADAMTS7* expression in lymphoblastoid cell lines (LCLs) from the Multi-ancestry Analysis of Gene Expression (MAGE) dataset. The centerline of each box plot indicates the median and the lower and upper hinges indicate the 25th and 75th percentiles, respectively. The vertical line of each boxplot extends to 1.5× the interquartile range from each hinge.

**a**

**b**

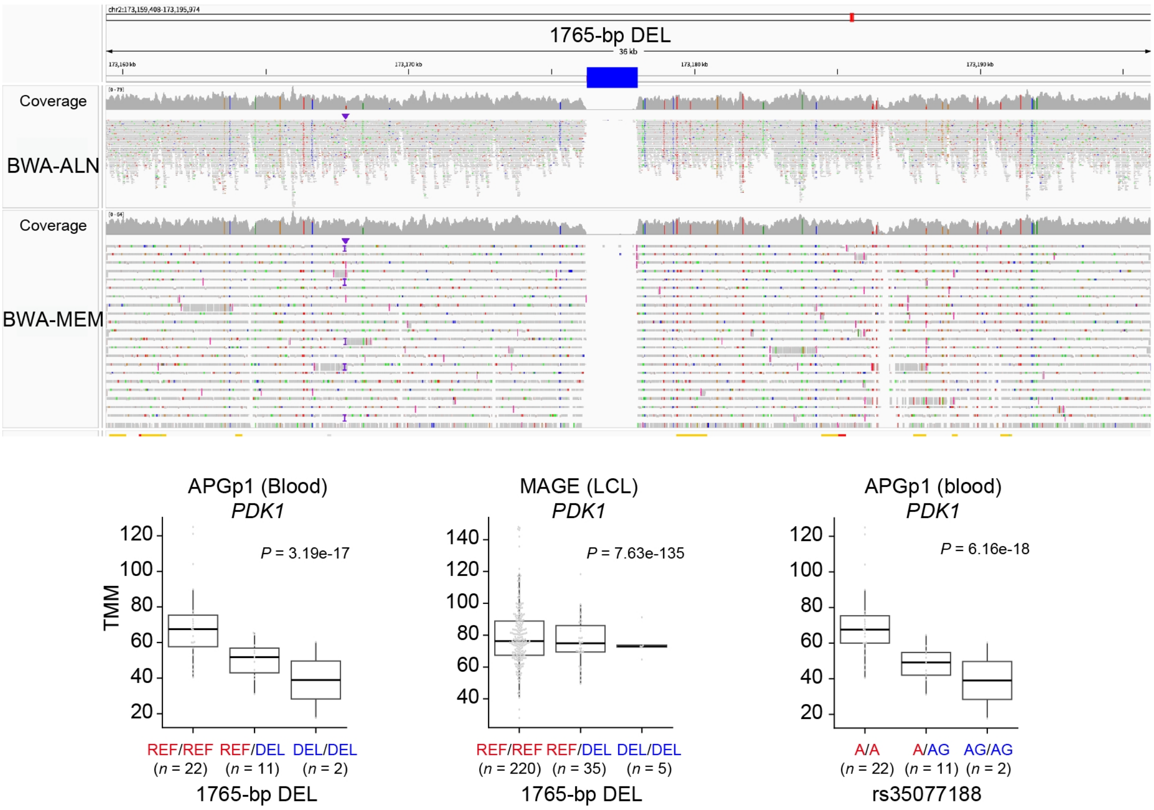


**Supplementary Fig. 22 | Characterization of a Neanderthal-derived deletion at locus 2q31.1. a**, IGV screenshot of read mapping from the Altai Neanderthal genome to the T2T-CHM13 reference, validating a 1,765-bp Neanderthal-derived deletion. Mapping profiles generated by BWA ALN and BWA MEM are shown, respectively. **b**, The introgressed 1,765-bp deletion and its linked InDel (rs35077188) are associated with the downregulation of *PDK1* expression in blood from APGp1 individuals and lymphoblastoid cell lines (LCLs) from the Multi-ancestry Analysis of Gene Expression (MAGE) dataset. The centerline of each box plot indicates the median and the lower and upper hinges indicate the 25th and 75th percentiles, respectively. The vertical line of each boxplot extends to 1.5× the interquartile range from each hinge.

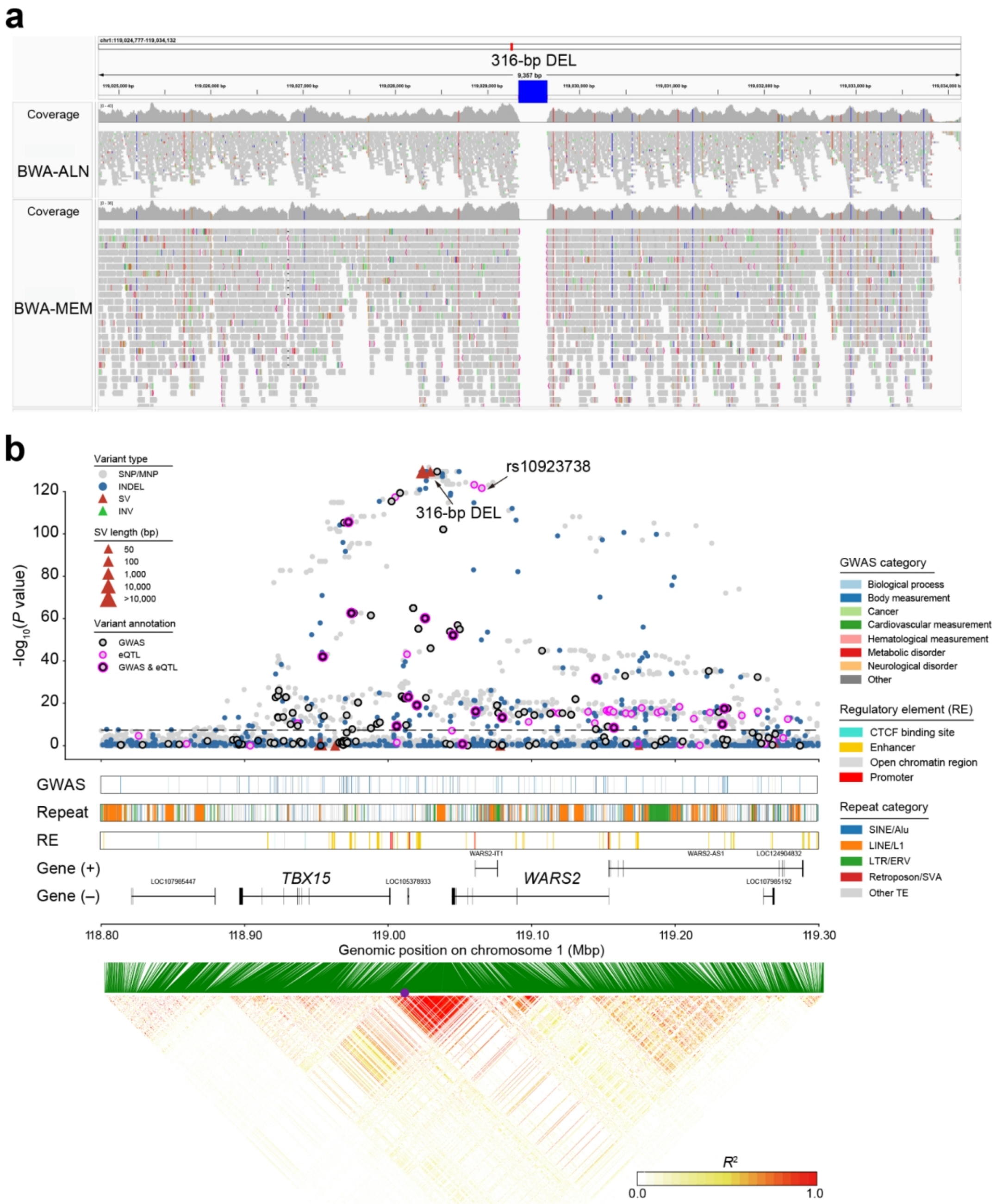


**Supplementary Fig. 23 | Characterization of a Denisovan-derived deletion at the *TBX15-WARS2* locus (1p12). a**, IGV screenshot of read mapping from the Denisovan genome to the T2T-CHM13 reference, validating a 316-bp Denisovan-derived deletion. Mapping profiles by BWA ALN and BWA MEM are shown, respectively. **b**, Genomic view of the 1p12 locus. The top panel displays the association between introgression and allele frequencies using Fisher's exact test for each variant, colored by variant type and functional annotation, including GWAS and eQTL signals. The middle panel integrates tracks for GWAS signals (colored by trait categories), repetitive elements (annotated by Repeatmasker), regulatory elements (Ensembl 113), and gene models (Refseq). The bottom panel shows a genetic linkage disequilibrium heatmap, with the 316-bp deletion highlighted by a purple dot.

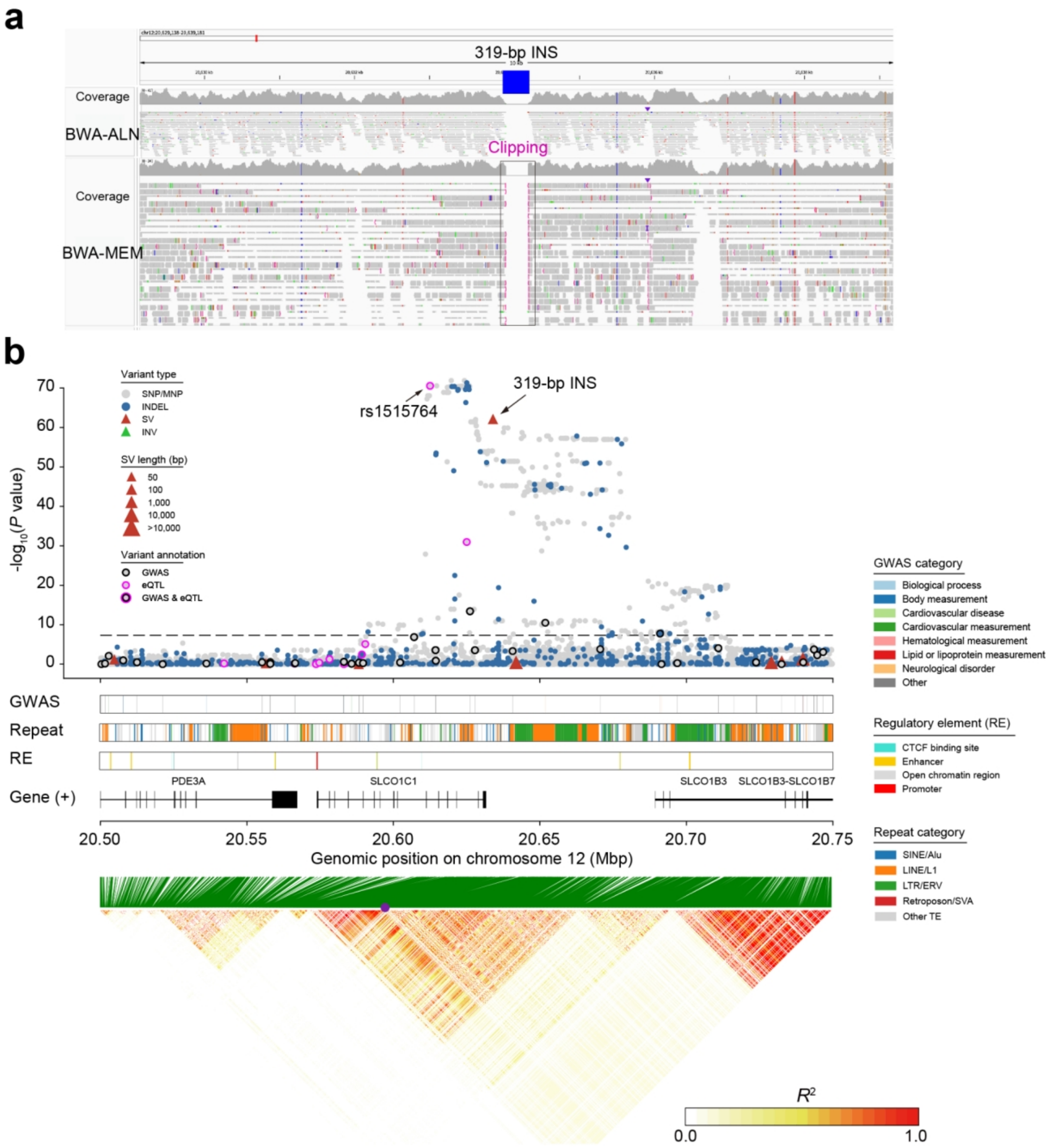


**Supplementary Fig. 24 | Characterization of a Denisovan-derived insertion at the 12p12.2 locus. a**, IGV screenshot of read mapping from the Denisovan genome to the T2T-CHM13 reference, suggesting the absence of the 319-bp Denisovan-derived insertion, which implies the deep structures and divergence among Denisovan lineages. Mapping profiles by BWA ALN and BWA MEM are shown, respectively. **b**, Genomic view of the12p12.2 locus. The top panel displays the association between introgression and allele frequencies using Fisher's exact test for each variant, color-coded by variant type and functional annotation, including GWAS and eQTL signals. The middle panel integrates tracks for GWAS signals (colored by trait categories), repetitive elements (annotated by Repeatmasker), regulatory elements (Ensembl 113), and gene models (Refseq). The bottom panel shows a genetic linkage disequilibrium heatmap, with the 319-bp insertion indicated by a purple dot.

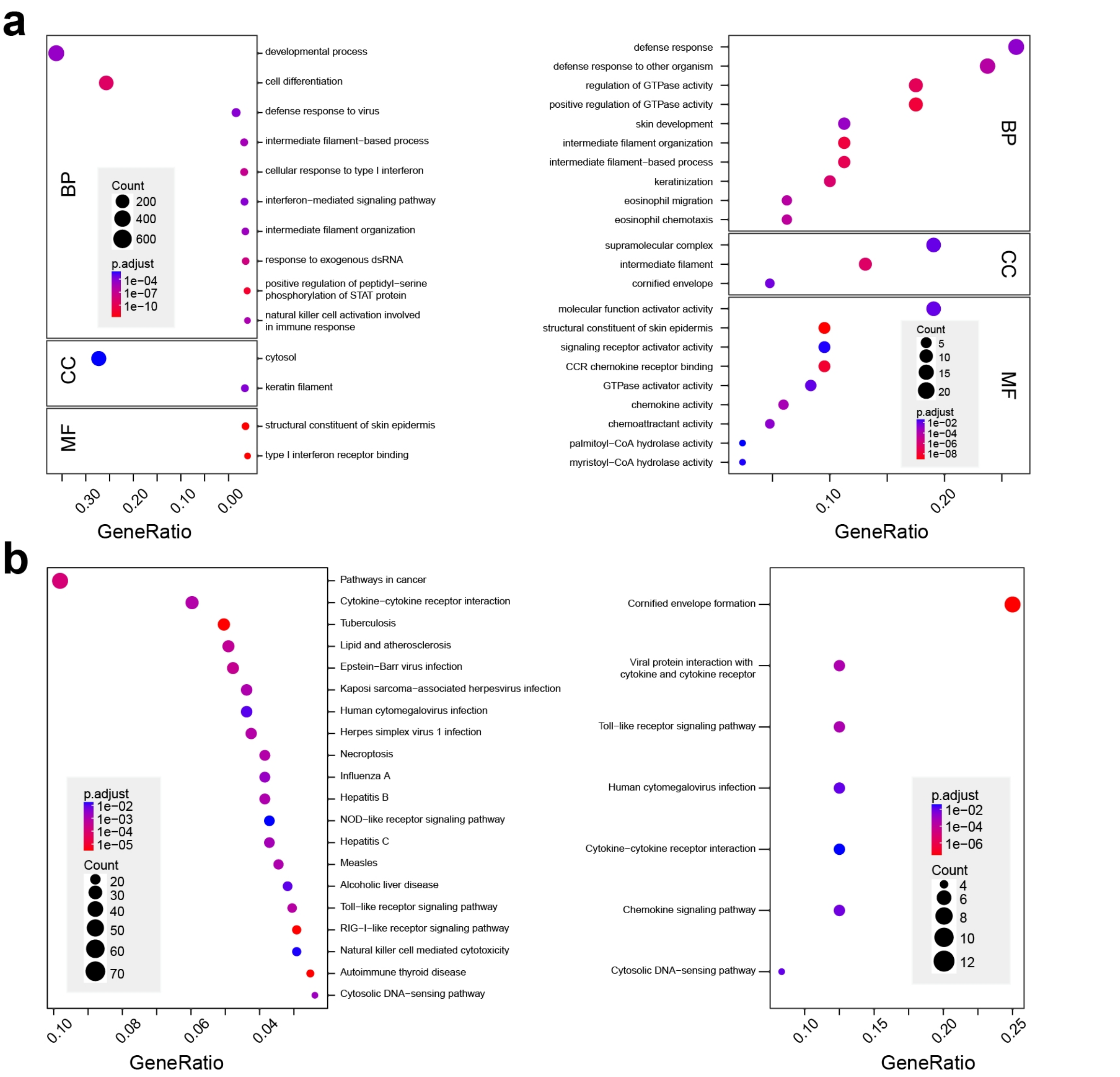


**Supplementary Fig. 25 | Functional enrichment of high-frequency (HF) archaic introgressed genes in modern human genomes.** Gene Ontology (GO; **a**) and KEGG (**b**) enrichment analysis is performed for HF genes derived from Neanderthal (left) and Denisovan (right) introgression.

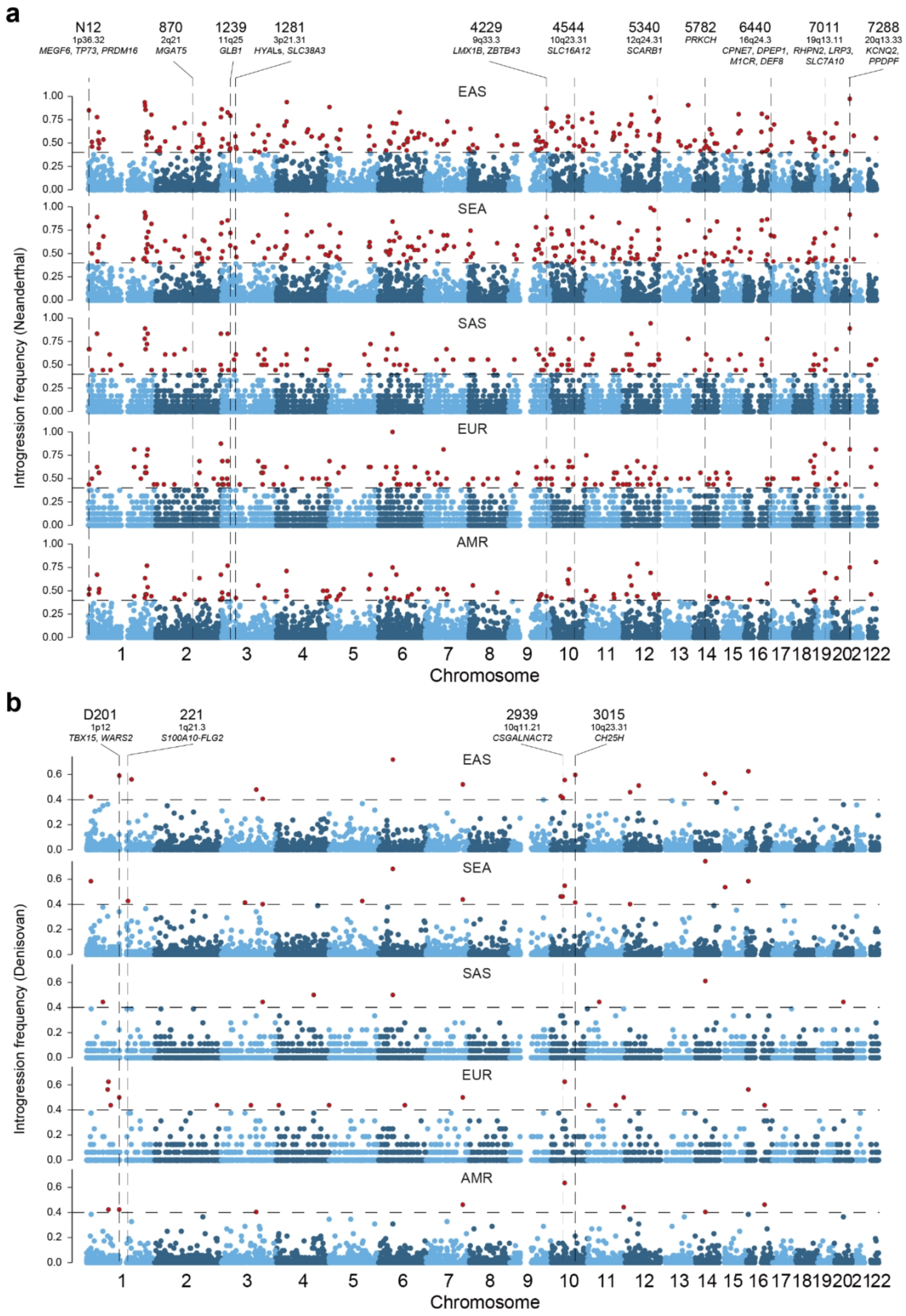


**Supplementary Fig. 26 | Frequency distribution of archaic introgressed chunks across all chromosomes. a**, Neanderthal-derived chunks. **b**, Denisovan-derived chunks. Chunks with an archaic allele frequency exceeding 0.4 are highlighted in red. Selected high-frequency chunks are labeled with their corresponding IDs and key genes located within these regions.

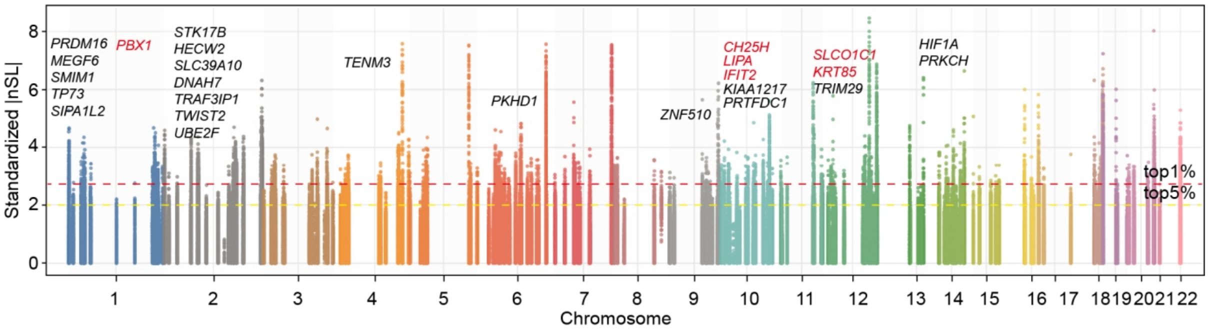


**Supplementary Fig. 27 | Manhattan plot of standardized |nSL| scores for SNVs in high-frequency introgressed chunks in the EAS superpopulation.** The dashed horizontal lines indicate the top 5% and top 1% thresholds for positive selection signals. Candidate genes under selection are annotated, with labels in black denoting Neanderthal-derived introgressed genes and those in red indicating Denisovan-derived introgressed genes.

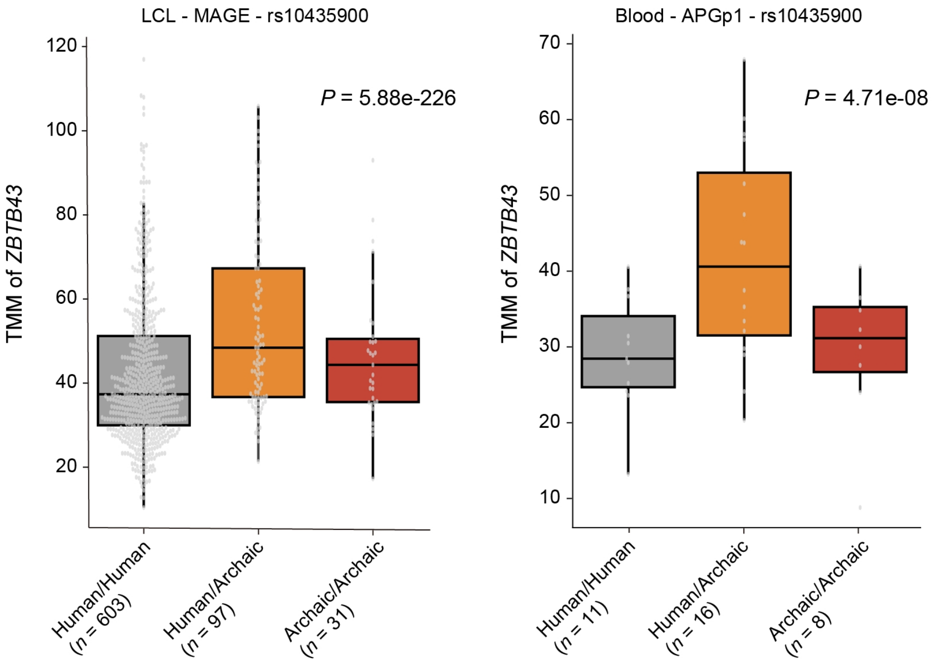


**Supplementary Fig. 28 | Association between *ZBTB43* expression and Neanderthal introgression.** Boxplots illustrate the normalized expression levels of *ZBTB43* stratified by genotype at the Neanderthal-introgressed locus rs10435900. Analyses were performed using Lymphoblastoid cell lines (LCLs) in the Multi-ancestry Analysis of Gene Expression (MAGE) dataset (left) and whole blood samples from APGp1 (right). Genotypes (*x*-axis) are categorized as non-introgressed (modern human) homozygotes, heterozygotes, and Neanderthal-introgressed homozygotes. In each box plot, the horizontal line shows the median value and the whiskers the 25% and 75% quartile values. The two-tailed *P* values for the *t*-test are derived from a linear regression model estimated using ordinary least squares.

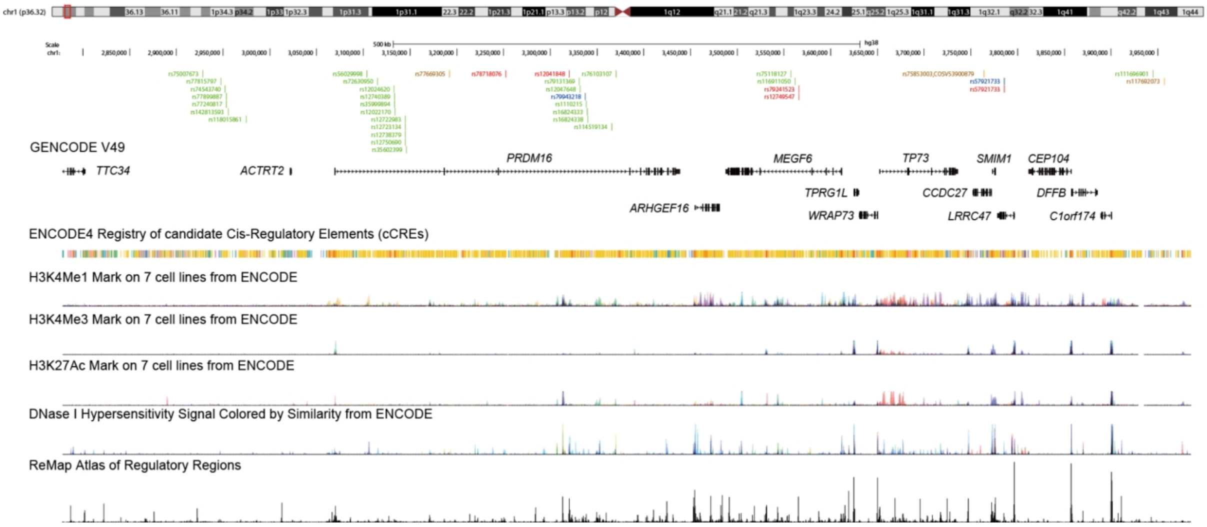


**Supplementary Fig. 29 | Functional landscape of the Neanderthal introgression chunk N12 (1p36.32).** This panel presents a view of the UCSC Genome Browser tracks for the Neanderthal introgression chunk N12. These tracks include gene models, candidate cis-regulatory elements (cCREs), chromatin accessibility signals (DNase I hypersensitivity sites), histone modification marks associated with active enhancers (H3K27ac), active promoters (H3K4me3), and poised/active regulatory elements (H3K4me1). Introgressed variants are highlighted and color-coded based on their predicted functional impact, including enhancer (green), CTCF binding site (red), transcription-factor binding site (blue), and open chromatin region (orange).

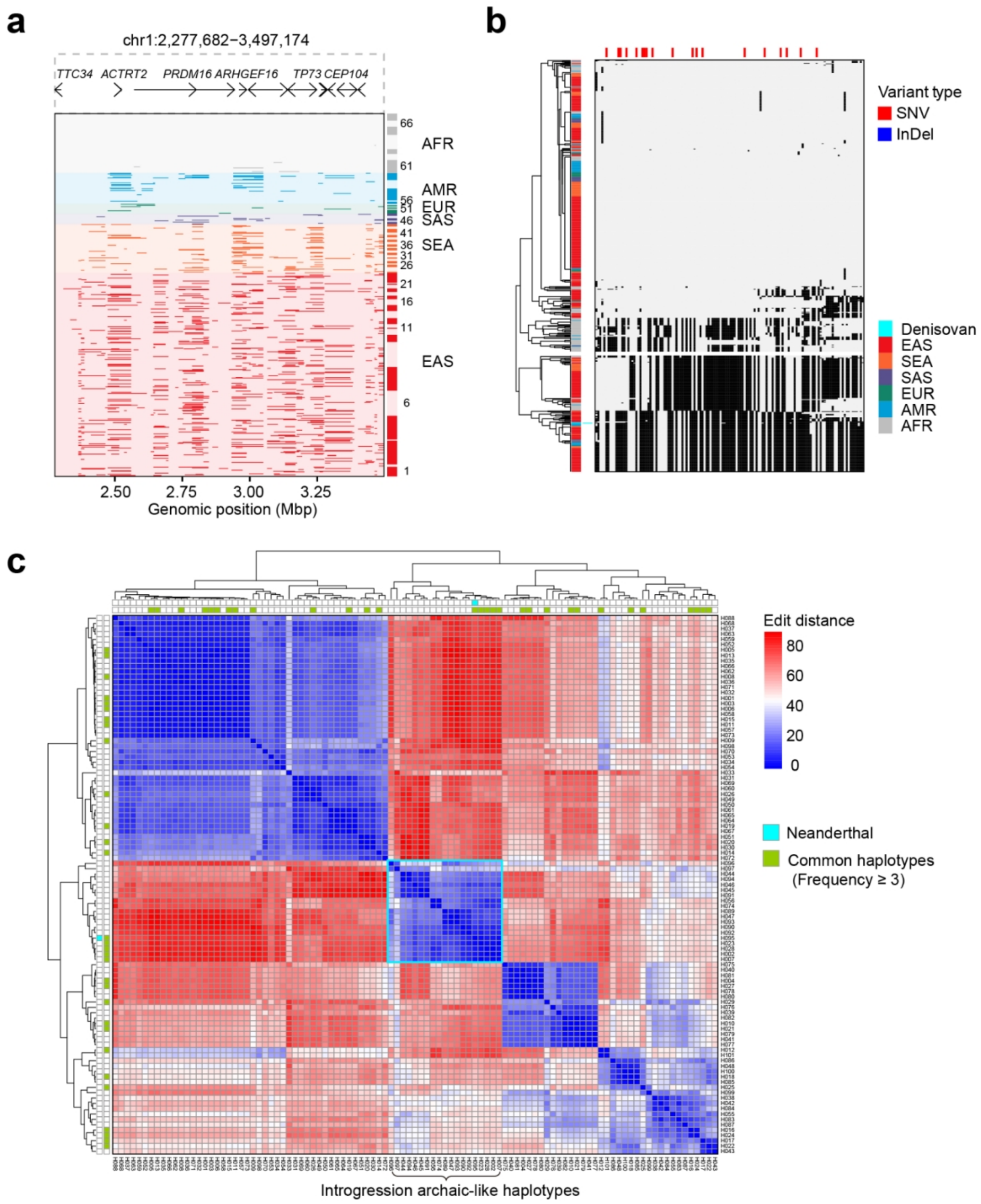


**Supplementary Fig. 30 | Genomic haplotypes of the Neanderthal introgression chunk N12 at 1p36.32. a**, Genomic landscape of the ~1-Mbp introgressed haplotype block (T2T-CHM13 coordinate: chr1:2,309,368-3,389,548), encompassing multiple genes including *PRDM16*. Population IDs are defined in Supplementary Fig. 7. Neanderthal-derived segments are displayed in dark red (EAS), orange (SEA), green (EUR), blue (AMR) and gray (AFR). **b**, Fine-scale haplotype heatmap of the core introgression region (chr1:2,482,304-2,529,545; 47.24 kbp). Rows represent 610 haploid assemblies plus the high-coverage Altai Neanderthal genome, clustered hierarchically. Columns represent 134 variant positions, with 19 Neanderthal-derived SNVs marked by red vertical lines. Allele states are indicated in gray (ancestral) and black (derived). These variants define 101 unique haplotypes in this region. **c**, Pairwise genetic distance between all identified haplotypes. Common haplotypes (frequency ≥ 3, *n* = 29) are indicated by green. The Neanderthal haplotype (H095) clusters closely with common introgressed haplotypes H002 (*n* = 53), H007 (*n* = 12), H023 (*n* = 4) and H028 (*n* = 3).

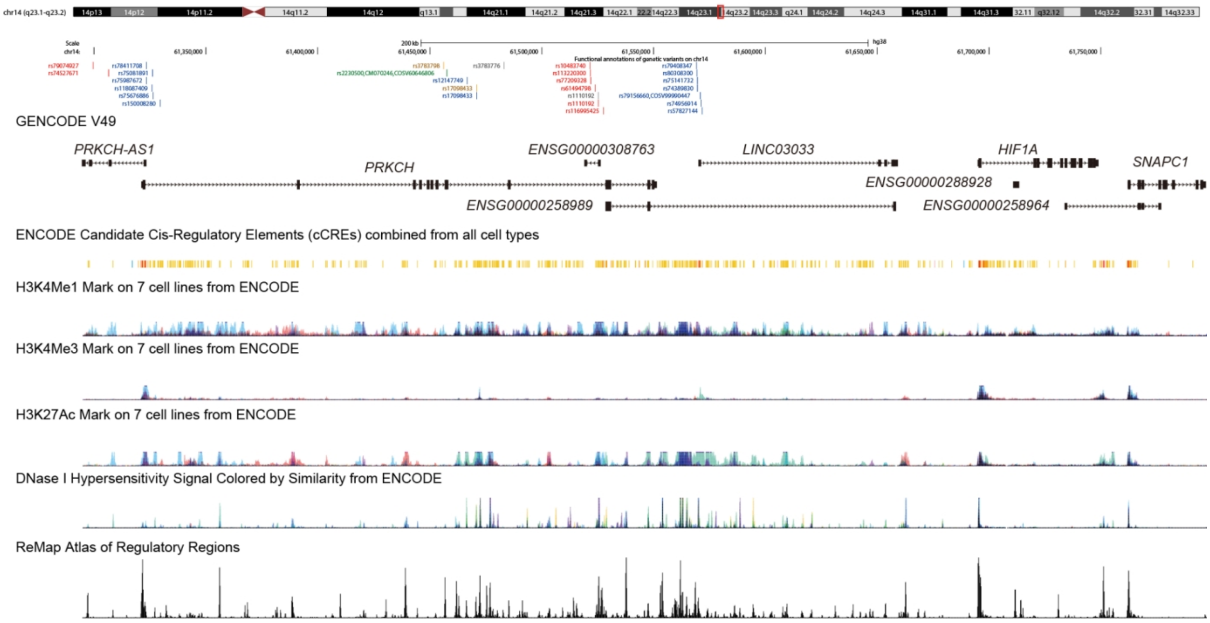


**Supplementary Fig. 31 | Functional landscape of the Neanderthal introgression chunk N5782 (14q23.1).** This panel presents a view of the UCSC Genome Browser tracks for the introgressed haplotype block N5782. The displayed tracks include gene models, candidate cis-regulatory elements (cCREs), chromatin accessibility signals (DNase I hypersensitivity sites), histone modification marks associated with active enhancers (H3K27ac), active promoters (H3K4me3), and poised/active regulatory elements (H3K4me1). Introgressed variants are highlighted and color-coded based on their predicted functional impact, including enhancer (red), promoter (blue), CTCF binding site (orange) and missense variant (green).

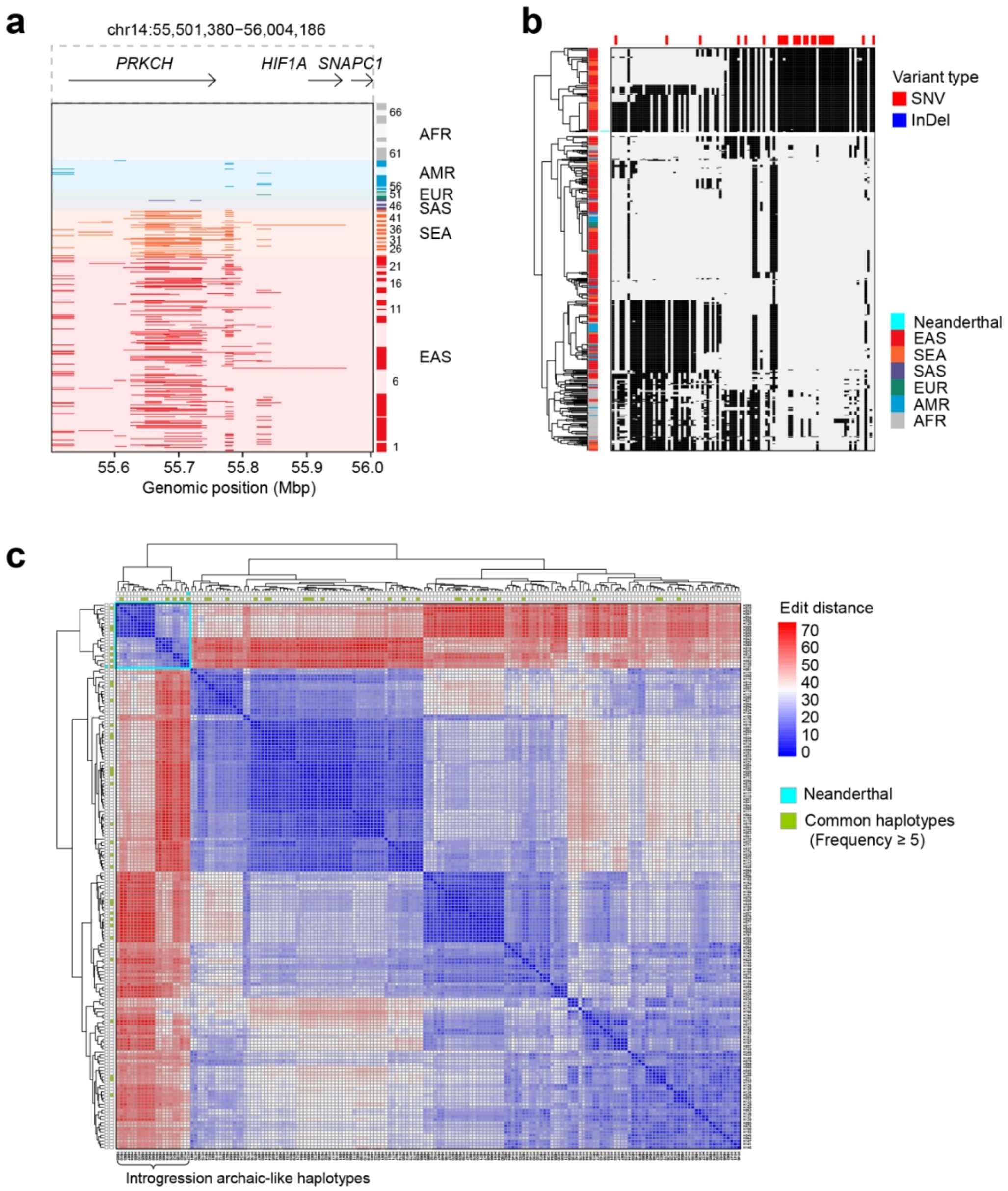


**Supplementary Fig. 32 | Genomic haplotypes of the Neanderthal introgression chunk N5782 at 14q23.1. a**, Genomic landscape of the ~419.8-kbp introgressed chunk (T2T-CHM13 coordinates: chr14:55,542,624-55,962,450). The region encompasses multiple genes including *PRKCH*, *HIF1A* and *SNAPC1*. Haplotypes from global haploid assemblies are plotted as rows, grouped and ordered by populations as defined in Supplementary Fig. 7. Neanderthal-derived segments are highlighted in dark red (EAS), orange (SEA), green (EUR), blue (AMR) or gray (AFR). **b**, Fine-scale haplotype heatmap of core introgression region (chr14:55,627,128-556,719,485; 44.82 kbp). Rows represent 610 haploid assemblies plus the high-coverage Altai Neanderthal genome, clustered hierarchically. Columns represent 103 variant positions, with 25 Neanderthal-derived SNVs marked with red vertical lines. Allele states are shown in gray (ancestral) and black (derived). These variants define 177 unique haplotypes in this region. **c**, Pairwise genetic distances between all unique haplotypes. Common haplotypes (frequency ≥ 5, $n$ = 31) are annotated by green. The Neanderthal haplotype (H127) clusters closely with common introgressed haplotypes H002 ($n$ = 44), H005 ($n$ = 18), H008 ($n$ = 14), and H010 ($n$ = 12), H012 ($n$ = 10) and H018 ($n$ = 8).

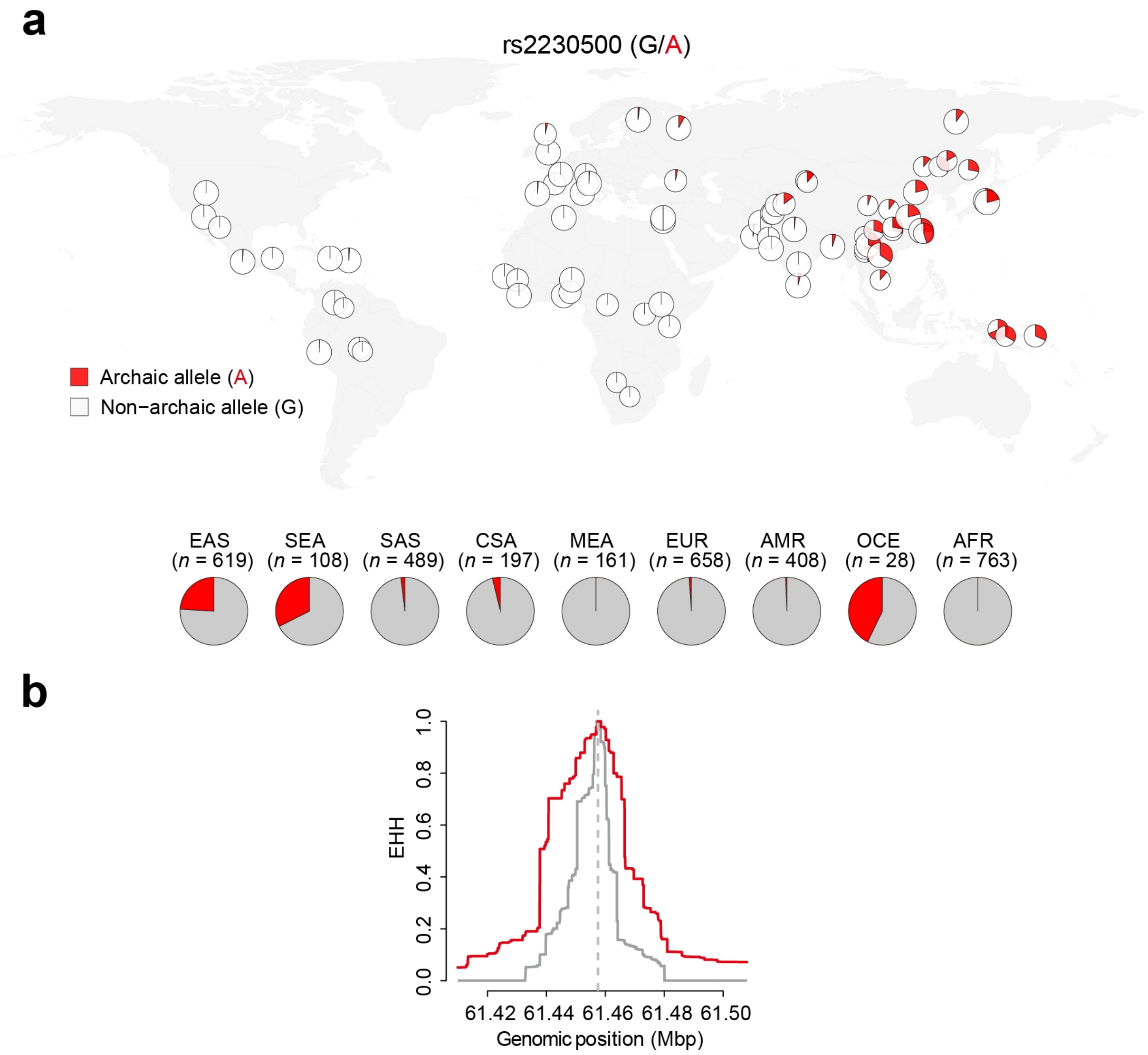


**Supplementary Fig. 33 | Global distribution and selection for a Neanderthal-derived allele at rs2230500 within *PRKCH*. a**, Global frequency distribution of the introgressed (archaic, derived, A) and non-introgressed (non-archaic, G) alleles at the focal variant rs2230500. Pie charts show the proportion of each allele across populations in the 1000 Genomes Project (1KGP) and the Human Genome Diversity Project (HGDP), with a pronounced regional enrichment of the archaic allele in EAS, SEA, and OCE superpopulations. **b**, Extended Haplotype Homozygosity (EHH) decay analysis for a 80-kbp region centered on rs2230500. The EHH decay curves are shown separately for haplotypes carrying the introgressed (derived) allele (red) and the non-archaic allele (gray).

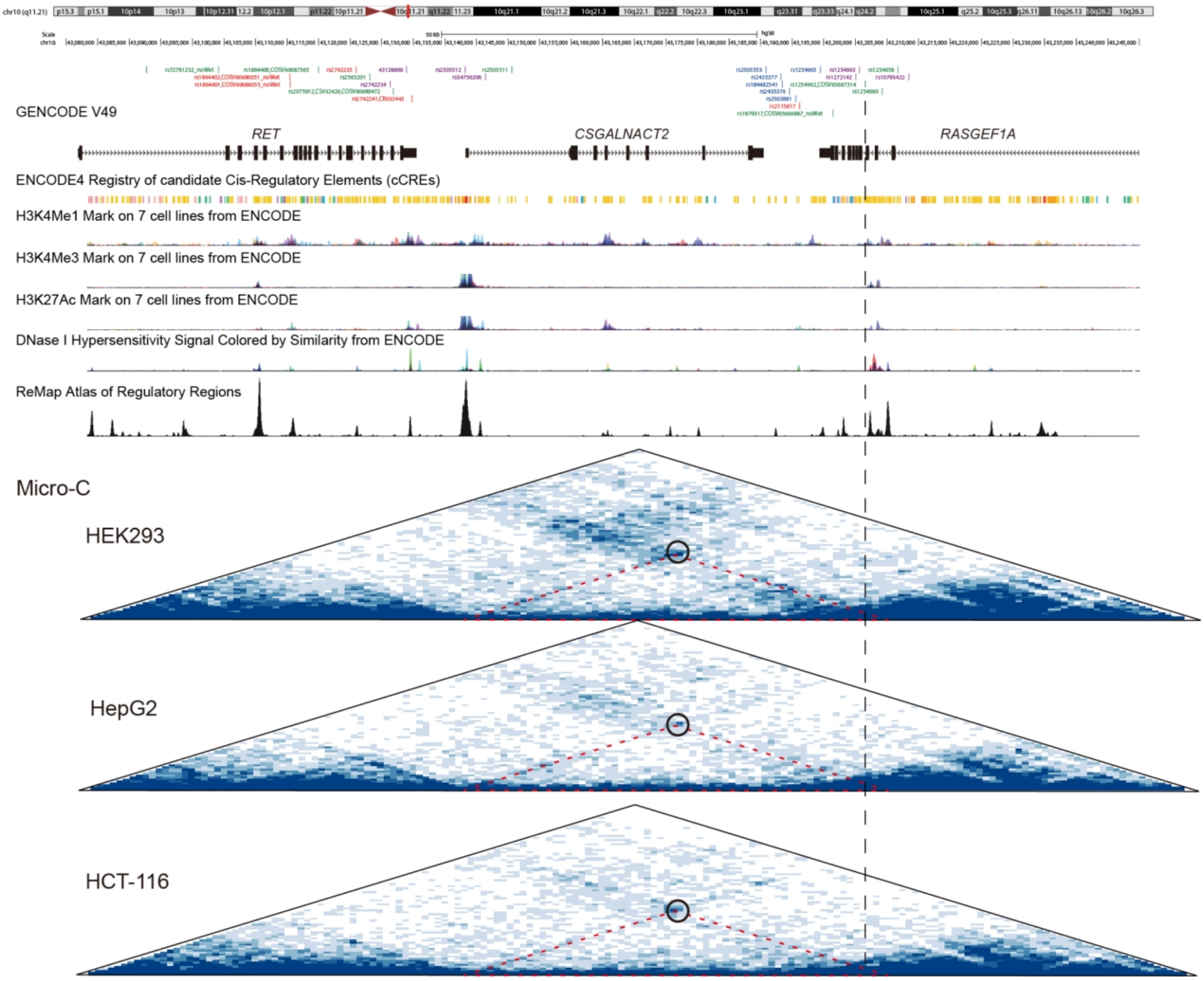


**Supplementary Fig. 34 | Functional landscape of the Denisovan-introgression chunk D2939 (10p11.21).** This panel presents a view of the UCSC Genome Browser tracks and Micro-C chromatin interaction maps in three cell lines, around the introgression block D2939. The displayed tracks include gene models, candidate *cis*-regulatory elements (cCREs), chromatin accessibility signals (DNase I hypersensitivity sites), histone modification marks associated with active enhancers (H3K27ac), active promoters (H3K4me3), and poised/active regulatory elements (H3K4me1). Introgressed variants are highlighted based on their predicted functional impact, including enhancer (red), intron variant (green), downstream gene variant (blue) and undefined variants (purple). Putative enhancer-promotor loops are highlighted by circles.

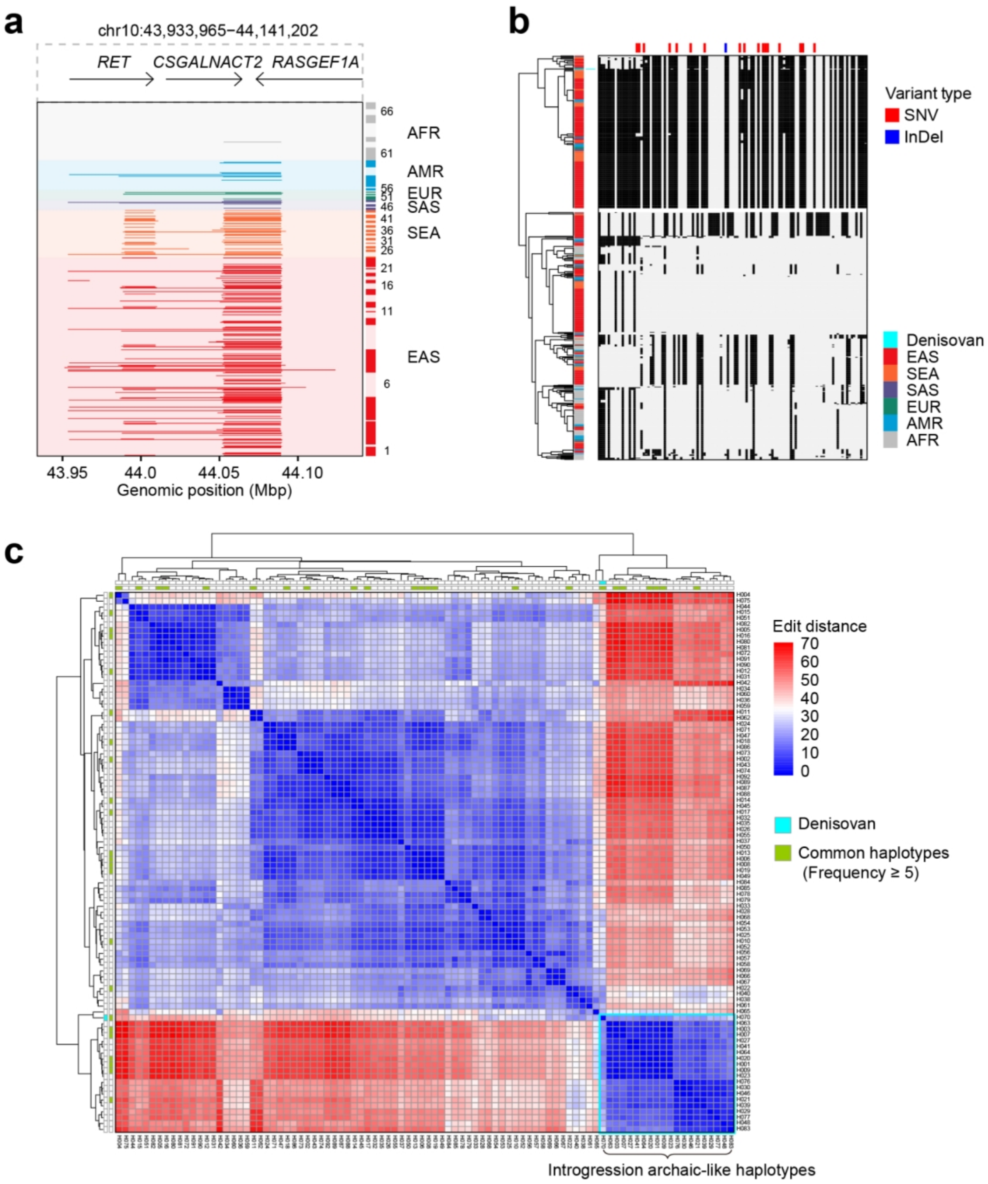


**Supplementary Fig. 35 | Genomic haplotypes of the Denisovan introgression chunk D2939 at 10q11.21. a**, Genomic landscape of the ~172.7-kbp introgressed block (T2T-CHM13 coordinates: chr10:43,951,235-44,123,932). The region encompasses multiple genes including *CSGALNACT2*, *RET* and *RASGEF1A*. Haplotypes from haploid assemblies are plotted as rows, grouped and ordered by populations defined in Supplementary Fig. 7. Denisovan-introgressed segments are displayed in dark red (EAS), orange (SEA), green (EUR), blue (AMR) or gray (AFR). **b**, Fine-scale haplotype heatmap of the core introgression region (chr10:43,980,253-44,102,763; 122.51 kbp). Rows represent 610 haploid assemblies plus the high-coverage Altai Denisovan genome, clustered hierarchically. Columns represent 115 variant positions, with 18 Denisovan-derived SNVs and one InDel marked with red and blue vertical lines, respectively. Allele states are shown in gray (ancestral) and black (derived). These variants define 92 unique haplotypes in this region. **c**, Pairwise genetic distances between all unique haplotypes. Common haplotypes (frequency ≥ 5, *n* = 22) are annotated by green. The Denisovan haplotype (H070) clusters closely with common introgression haplotypes H001 (*n*= 98), H003 (*n* = 51), H007 (*n* = 29), H009 (*n* = 16), H020 (*n* = 6) and H021 (*n* = 5).

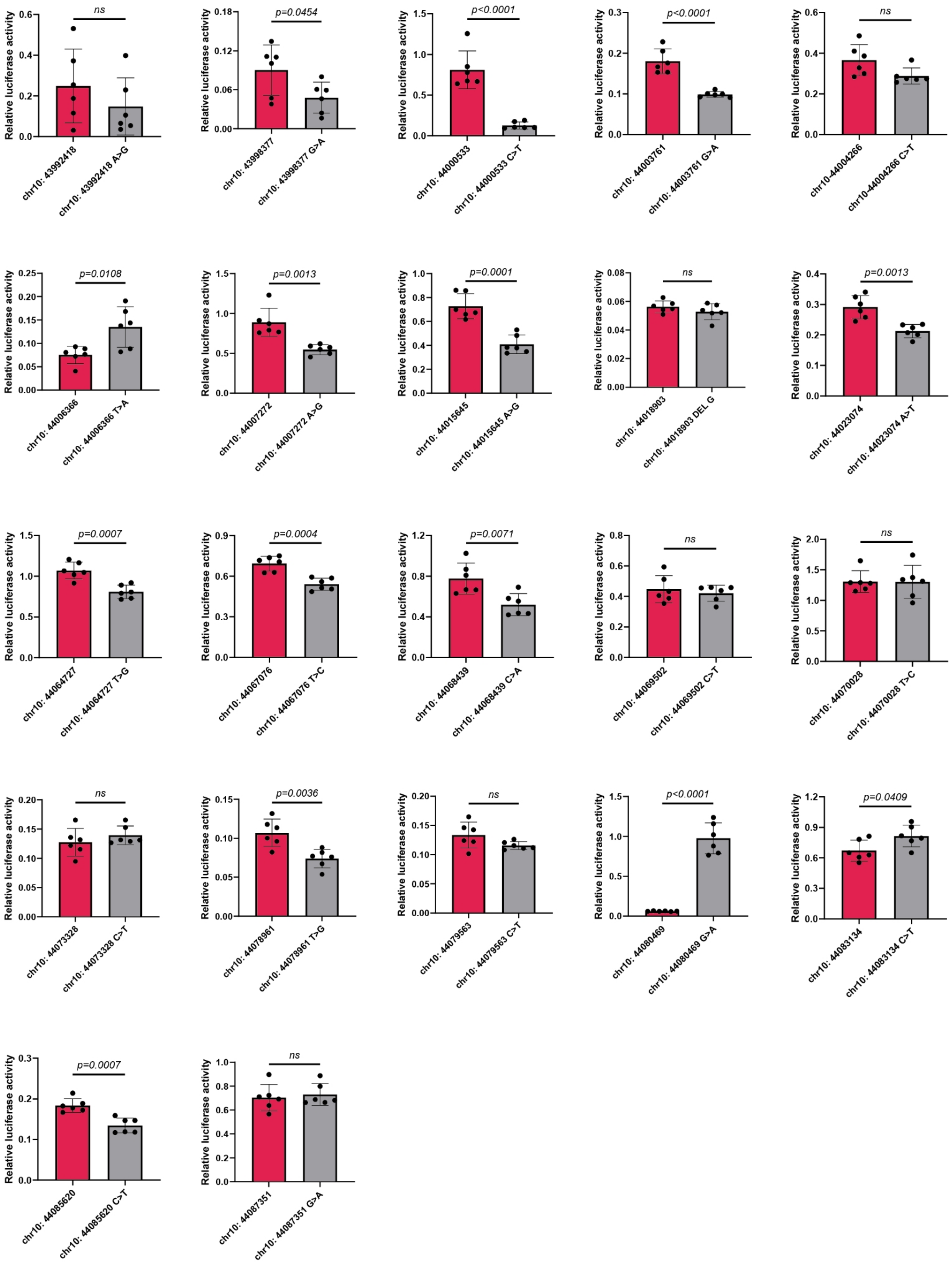


**Supplementary Fig. 36 | Functional profiling of regulatory activity via dual-luciferase reporter assays for Denisovan-derived variants at the *CSGALNACT2* locus.** Relative luciferase activity is measured in HEK293-T cells to assess the regulatory impact of the archaic variants. Data are presented as mean ± s.e.m. Statistical significance levels are calculated using two-sided *t*-test ($n$ = 6 replicates for each of the two groups).

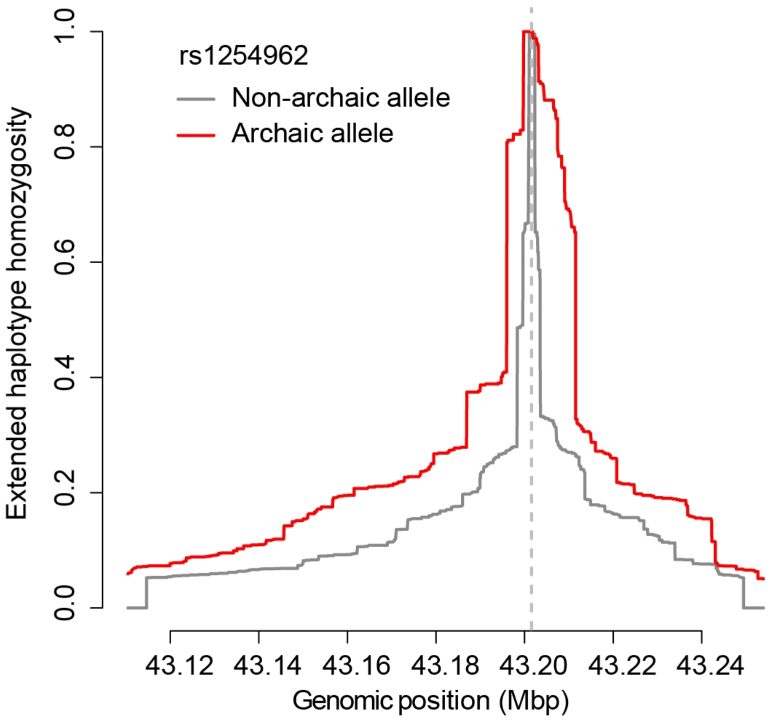


**Supplementary Fig. 37 | Signature of positive selection for the archaic allele at rs1254962 (*CSGALNACT2*).** Extended Haplotype Homozygosity (EHH) decay profiles within a 120-kbp region centered on the introgression variant rs1254962. Decay curves are plotted separately for haplotypes carrying the introgressed allele (red, A) and non-archaic allele (gray, G).

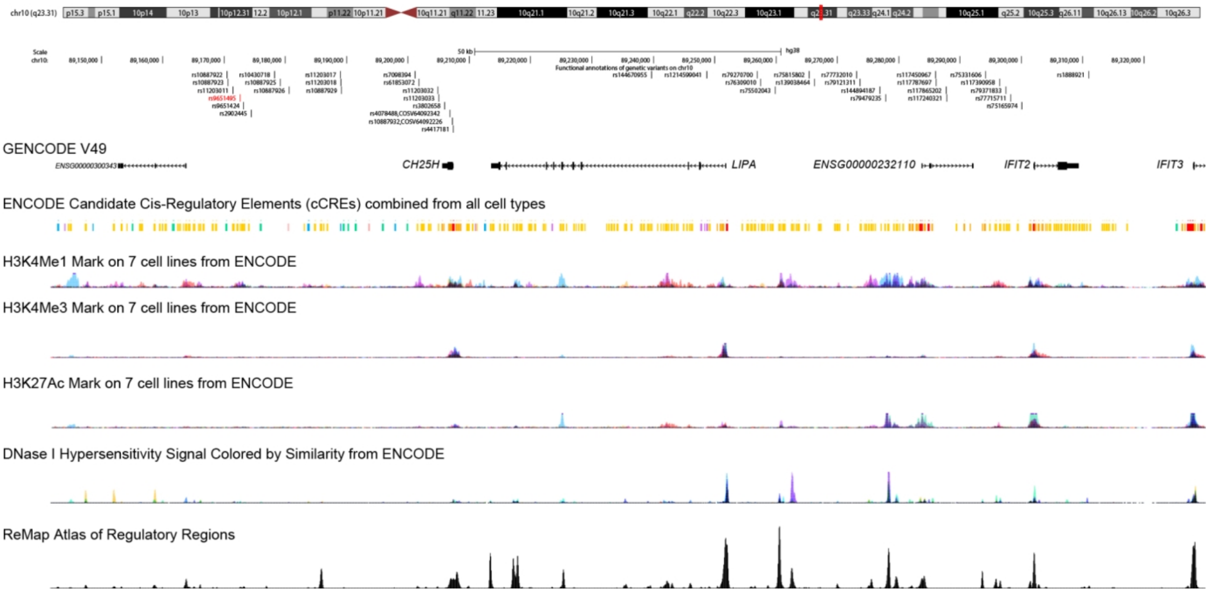


**Supplementary Fig. 38 | Functional landscape of the Denisovan introgression chunk D3015 (10q23.31).** This panel presents a view of the UCSC Genome Browser tracks for the introgressed haplotype block D3015, including gene models, candidate *cis*-regulatory elements (cCREs), chromatin accessibility signals (DNase I hypersensitivity sites), histone modification marks associated with active enhancers (H3K27ac), active promoters (H3K4me3), and poised/active regulatory elements (H3K4me1). Introgressed variants are specifically annotated, including enhancer (red) and undefined variants (gray).

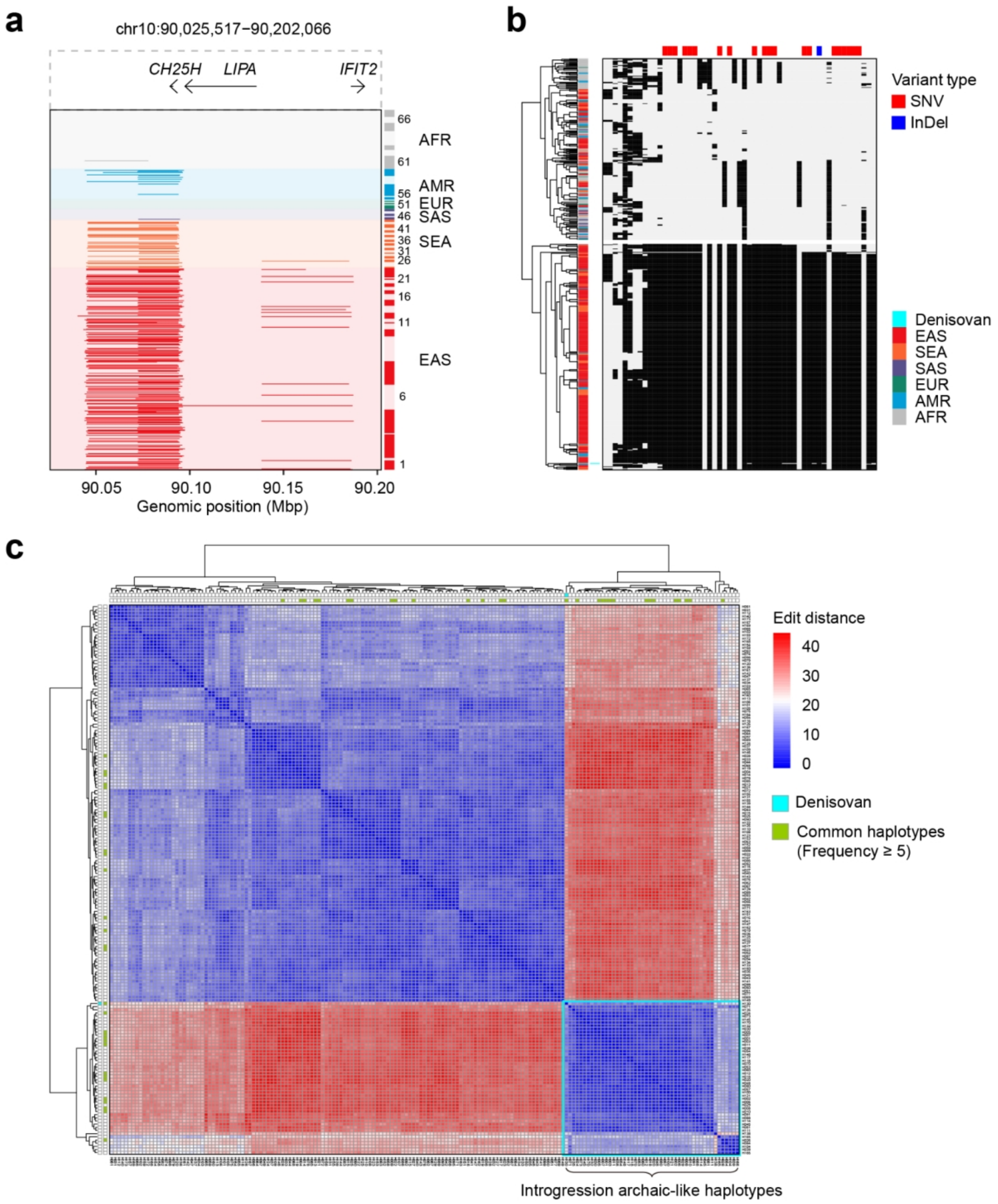


**Supplementary Fig. 39 | Genomic haplotypes of the Denisovan introgression chunk D3015 at 10q23.31. a**, Genomic landscape of the ~147.1-kbp introgressed block (T2T-CHM13 coordinates: chr10:90,040,229-90,187,353). The region encompasses multiple genes including *CH25H*, *LIPA* and *IFIT2*. Haplotypes from haploid assemblies are plotted as rows, grouped and ordered by populations defined in Supplementary Fig. 7. Denisovan-introgressed segments are displayed in dark red (EAS), orange (SEA), green (EUR), blue (AMR) or gray (AFR). **b**, A fine-scale haplotype heatmap of the core introgression region (chr10:90,025,601-90,092,916; 67.3 kbp). Rows represent 610 haploid assemblies plus the high-coverage Altai Denisovan genome, clustered hierarchically. Columns represent 55 variants, with 21 Denisovan-derived SNVs and one InDel marked by red/blue vertical lines. Allele states are shown in gray (ancestral) and black (derived). These variants define 173 unique haplotypes in this region. **c**, Pairwise genetic distances between all unique haplotypes. Common haplotypes (frequency ≥ 5, $n$ = 28) are annotated by green. The Denisovan haplotype (H130) clusters closely with 14 common introgressed haplotypes.

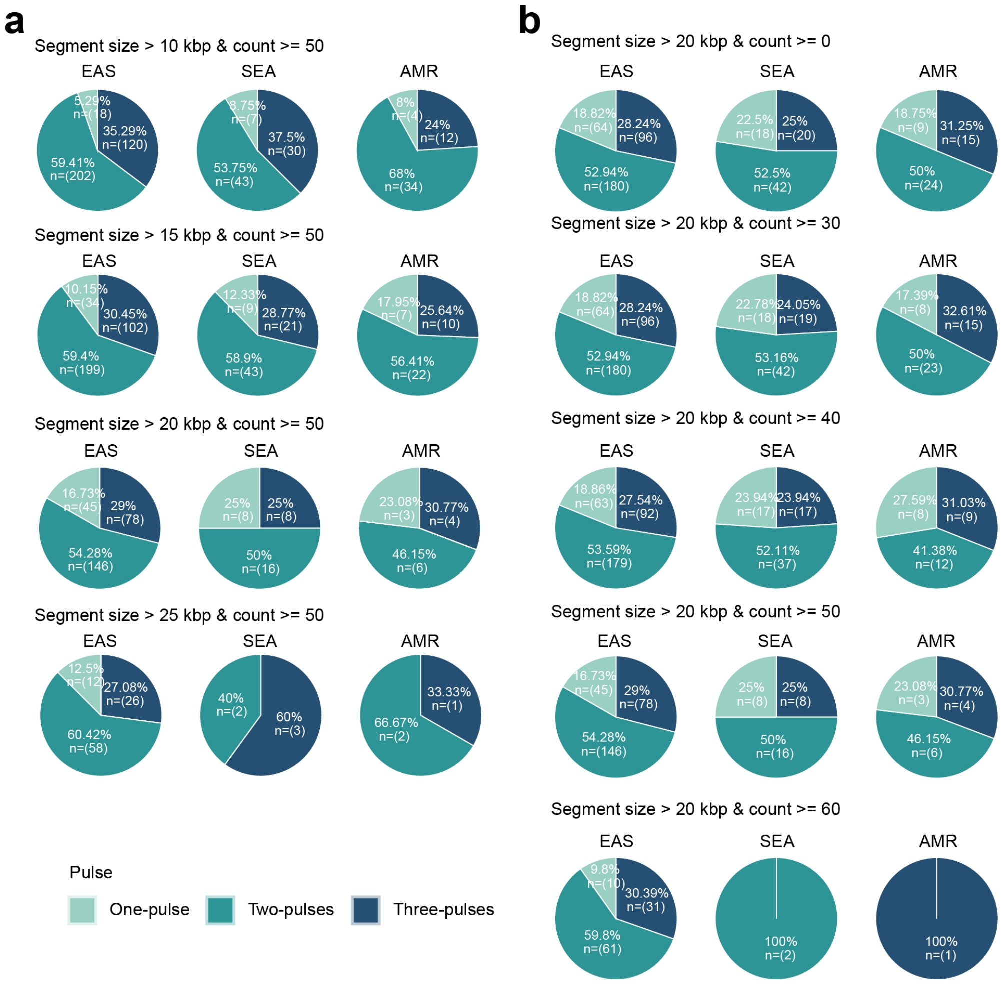


**Supplementary Fig. 40 | Inference of Denisovan introgression pulses using Gaussian mixture modeling (GMM).** The optimal number of Denisovan introgression pulses was inferred from Gaussian mixture modeling on haploid assemblies, with systematic evaluations of the impacts of archaic segment size (**a**) and segment count (**b**) on model selection. Pie charts represent the proportion of haploid genomes best fit by one, two, or three Denisovan introgression components, stratified by ancestry group (EAS, SEA and AMR). **a**, Different minimum segment length thresholds (10 kbp, 15 kbp, 20 kbp and 25 kbp) are applied to test the robustness of multiple archaic admixture events. **b**, Influence of the number of introgressed segments per assembly on pulse inference. Analyses were performed on assemblies with segments >20 kbp, testing various minimum segment count thresholds (>0, >30, >40, >50, and >60 segments).

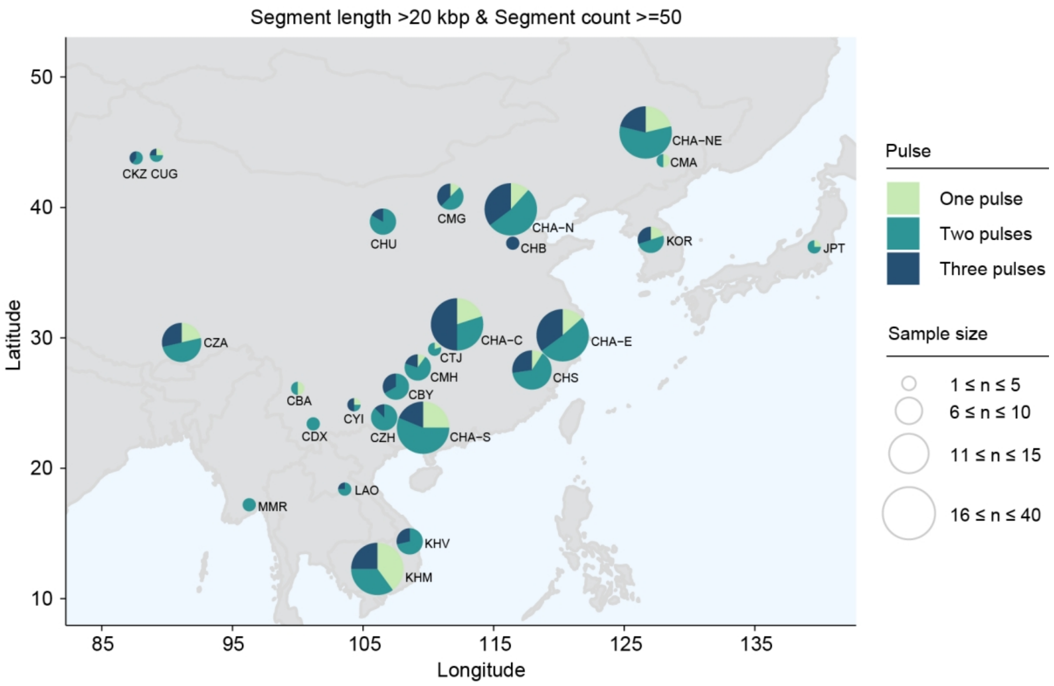


**Supplementary Fig. 41 | Geographical distribution of optimal Denisovan introgression models across East Eurasian populations.** The map shows the proportion of haploid genomes best explained by one, two or three Denisovan introgression components, represented by pie charts for each population.

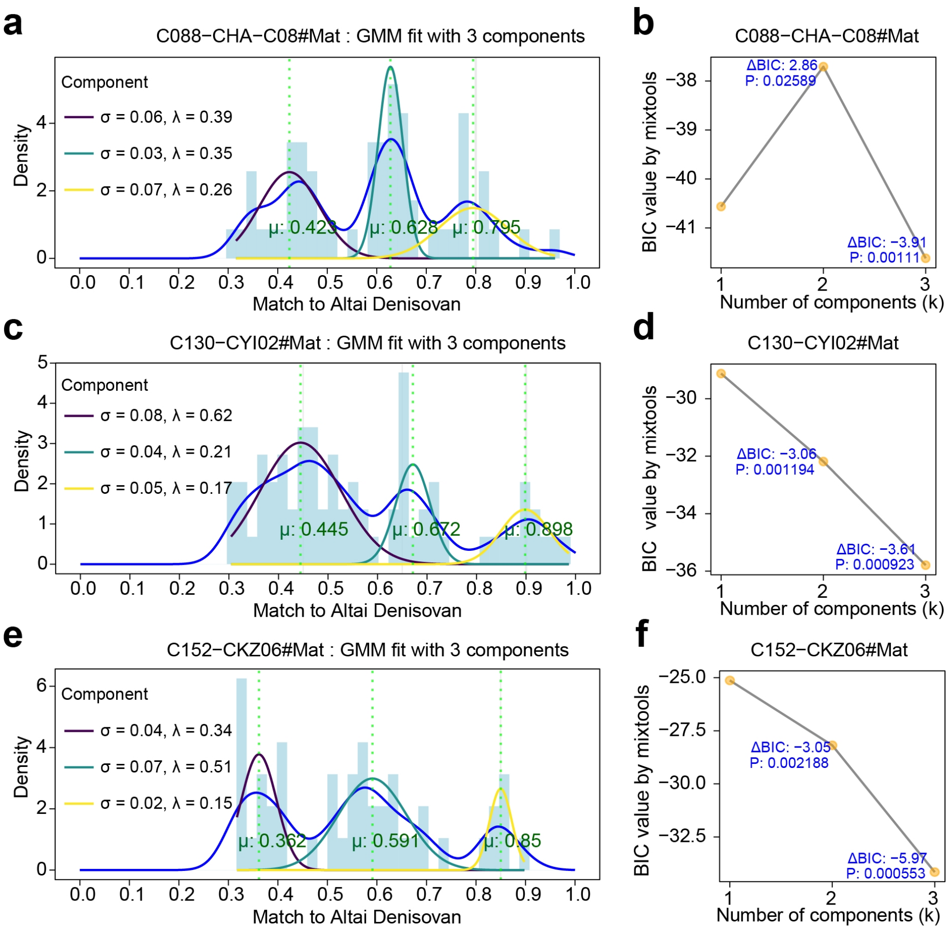


**Supplementary Fig. 42 | Representative examples of match rate distributions and Gaussian mixture modeling for inferring Denisovan introgression pulses in haploid-resolved assemblies. a**, **c** and **e**, Density distributions of match rates (blue curves) to the Altai Denisovan reference for three representative haploid genome assemblies (C088-CHA-C08#Mat, C130-CYI02#Mat and C152-CKZ06#Mat), overlaid with Gaussian components fitted under a three-pulse model. The three components are color-coded in deep violet, green verditer, and light goldenrod, representing distinct introgression lineages. Peaks correspond to inferred introgression events, with vertical dashed lines marking their mean match rates. **b**, **d** and **f**, Bayesian Information Criterion (BIC) values for one, two, and three-component Gaussian mixture models corresponding to the haploid assemblies shown in a, c, and e, respectively. ΔBIC values relative to the preceding model and *P* values from likelihood-ratio tests between nested models are displayed.

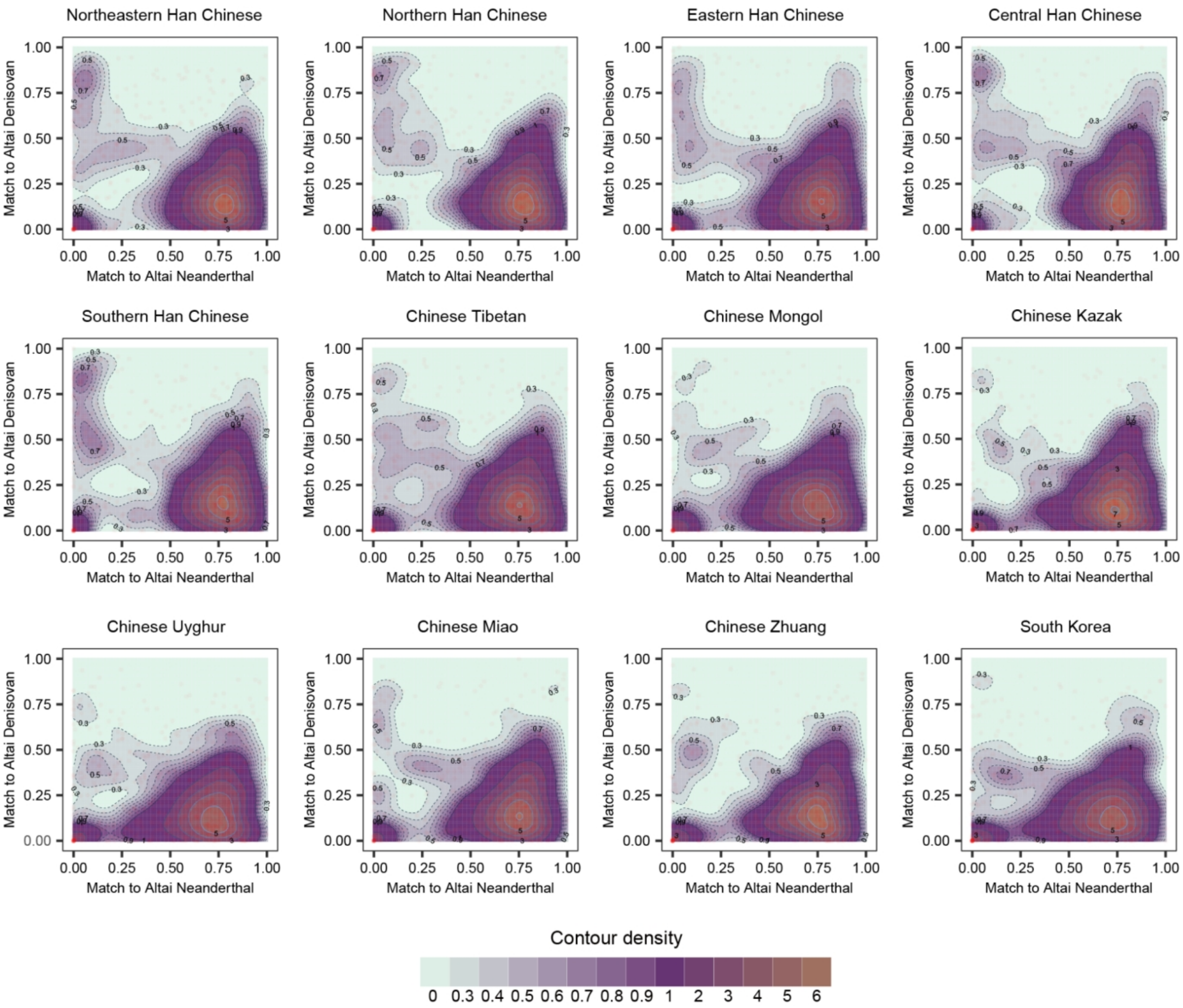


**Supplementary Fig. 43 | Contour density plots of match rates for introgressed segments against the Altai Neanderthal and Altai Denisovan genomes.** Match rate here is defined as the proportion of putative archaic-specific alleles with a segment that match the given archaic genome, as previously defined by Sprime (Browning et al., 2018). At least ten variants compared to the Neanderthal genome and at least ten variants compared to the Denisovan genome are required. The gradient colors and labeled contour lines represent the density height. Solid contour lines are drawn at integer multiples of 1, while dashed lines indicate multiples of 0.1 between 0.3 and 0.9 for finer resolution.

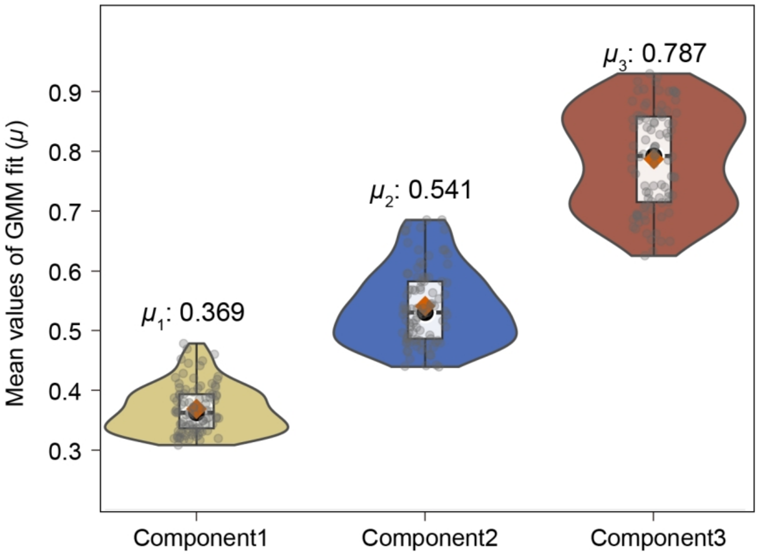


**Supplementary Fig. 44 | Distribution of mean match rates ($\mu$) for the three Denisovan introgression pulses.** Violin plot shows the mean match rate ($\mu$) for each of the three inferred Denisovan introgression pulses, Pulse 1 (Brass), Pulse 2 (Sapphire blue), and Pulse 3 (Oxide red), identified by Gaussian mixture modeling across 90 haploid genomes best fit by a three-component model. Within each violin, the centerline of each box plot indicates the median and the lower and upper hinges indicate the 25th and 75th percentiles, respectively. The vertical line of each boxplot extends to 1.5× the interquartile range from each hinge. Based on these $\mu$ value distributions, three Denisovan ancestry affinity categories are defined: low ($0.3 < MR < 0.45$), moderate ($0.45 < MR < 0.65$), and high ($0.7 < MR < 1.0$) affinity.

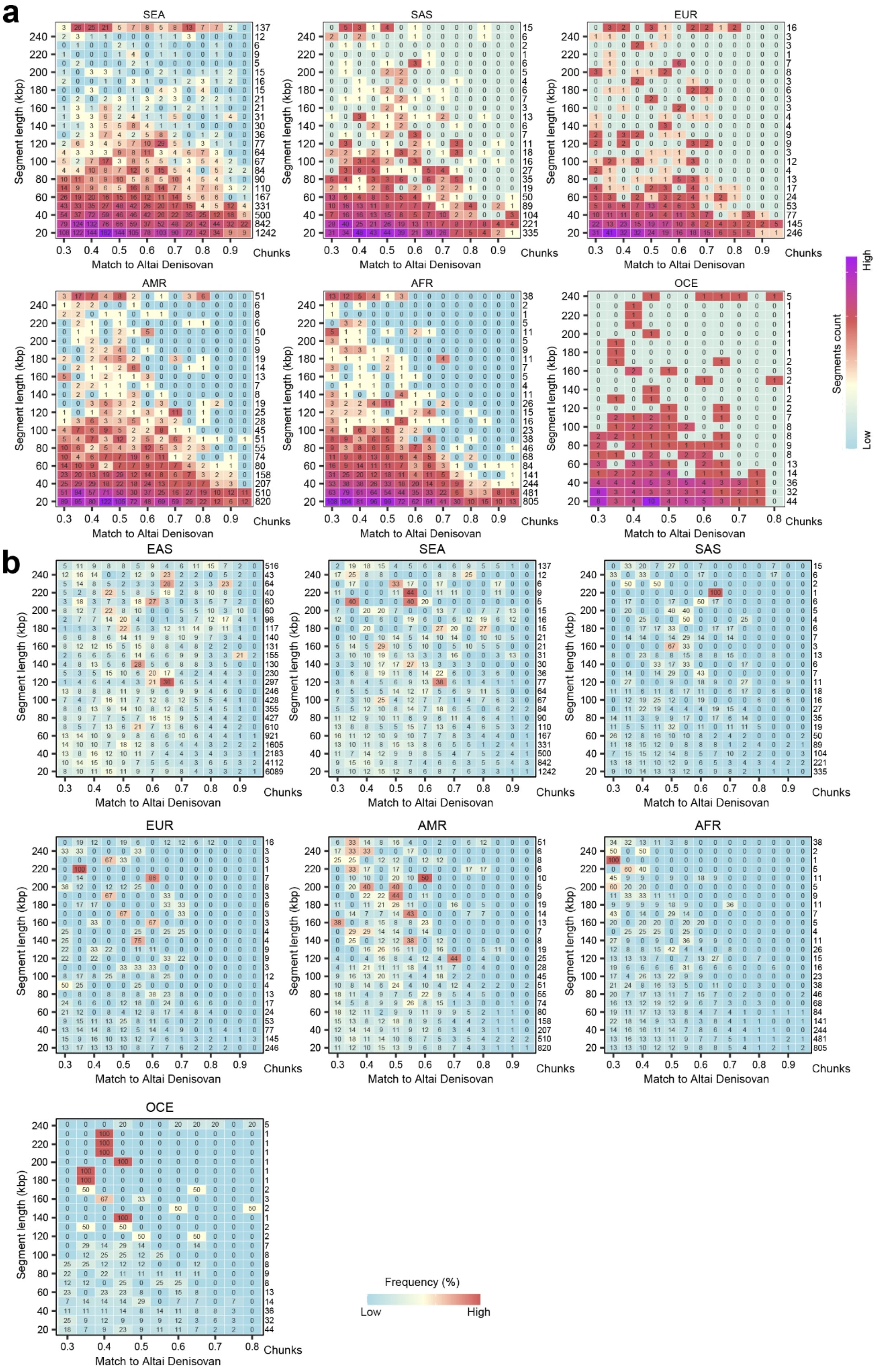
a
SEA
SAS
EUR
AMR
AFR
OCE
Segment length (kbp)
Match to Altai Denisovan
Chunks
Segments count
High
Low
b
EAS
SEA
SAS
EUR
AMR
AFR
OCE
Segment length (kbp)
Match to Altai Denisovan
Chunks
Frequency (%)
Low
High

**Supplementary Fig. 45 | Characterization of Denisovan-derived segments stratified by match rate, segment length and ancestral group. a**, Heatmaps illustrating the absolute counts of introgressed segments across binned match rates to the Altai Denisovan genome (*x*-axis; 0.05 increments from 0.3 to 1.0) and binned segment lengths (*y*-axis; 10-kbp increments from 20 kbp). The rightmost column provides the total number of segments per length bin. Cell colors and numerical annotations indicate the segment count within each match-rate and length bin. **b**, Heatmaps displaying the proportional distribution (row-wise percentage) of introgressed segments within each length bin across the same match-rate categories. Systematic profiling reveals distinct patterns of Denisovan ancestry structure. In Mainland Southeast Asian (SEA) populations, low-match-rate segments (0.4-0.45) are concentrated in shorter blocks (~20-50 kbp). As segment length increases, intermediate-match-rate segments (0.45-0.65) become predominant, peaking in segments longer than 110 kbp. For segments exceeding 150 kbp, high-affinity components (match rate 0.7-1.0) account for the largest proportion. The proportional view reinforces these patterns and further emphasizes population-specific profiles of Denisovan ancestry.

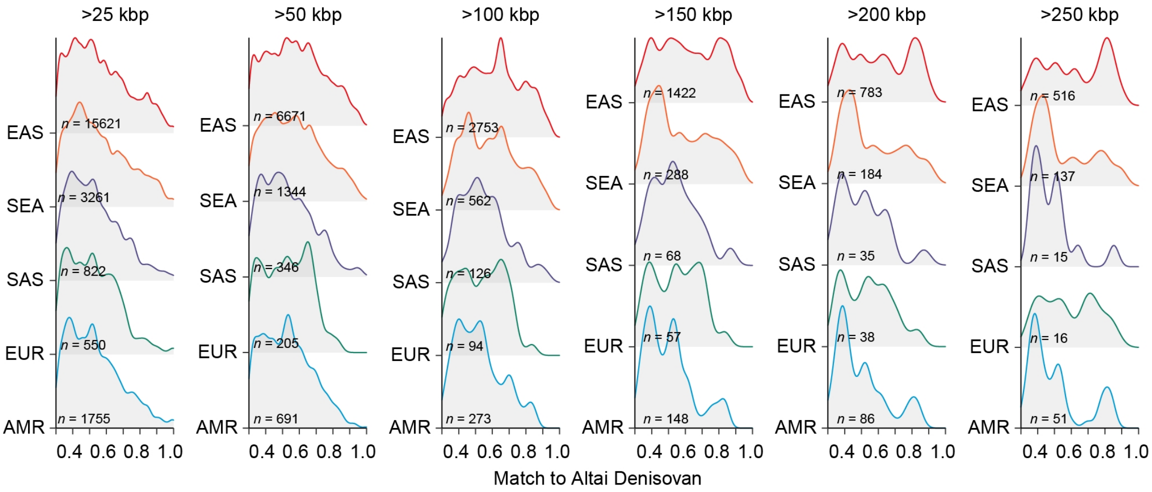


**Supplementary Fig. 46 | Distribution of Denisovan match rates across superpopulations under varying segment-length thresholds.** Ridge plots illustrate the density distribution of match rates to the Altai Denisovan genome, faceted by ancestry group (rows) and minimum segment-length thresholds (≥25, 50, 100, 150, 200 and 250 kbp), enabling detailed inspection of distribution shapes across populations. Systematic variation in match rate distributions is observed as segment-length thresholds increase. At the 25-kbp threshold, a pronounced peak appears in the low-affinity range across populations. As the threshold reaches 100 kbp, a distinct moderate-affinity peak emerges specifically in EAS. At thresholds ≥ 150 kbp, a high-affinity component becomes prominent, signaling the presence of recently introgressed Denisovan segments closely related to the Altai reference.

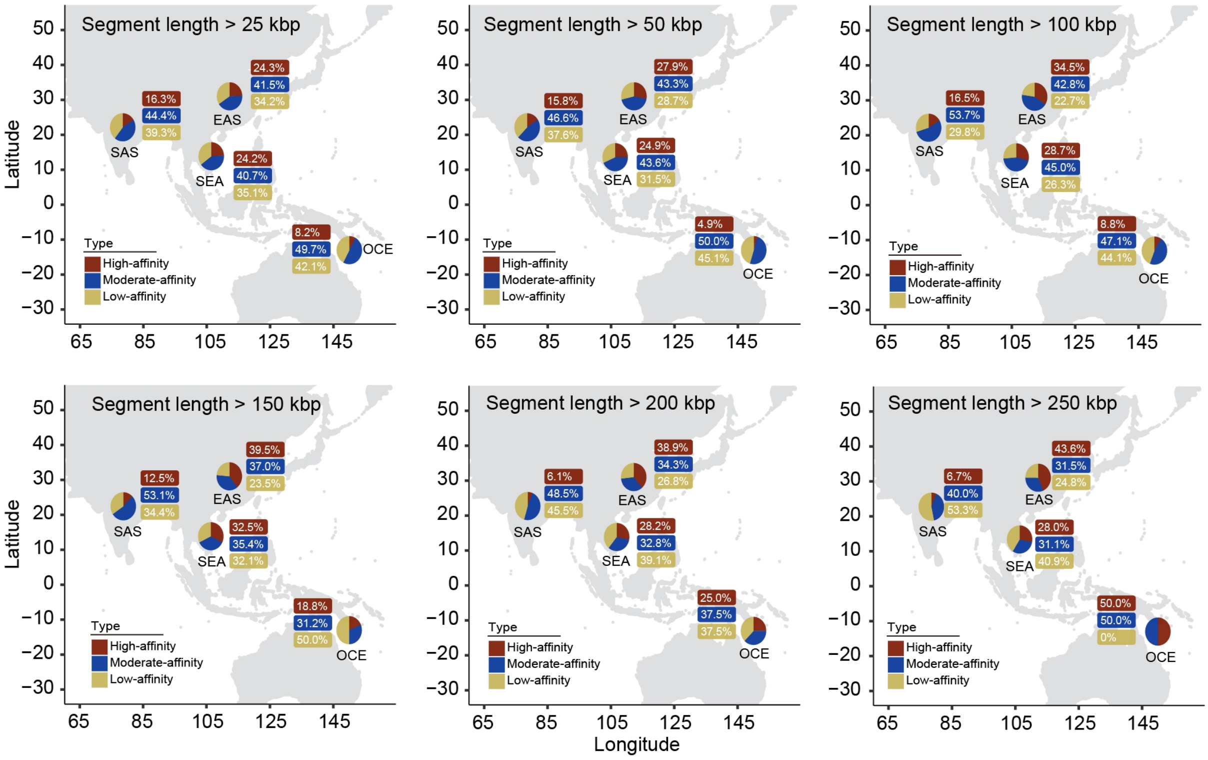


**Supplementary Fig. 47 | Geographic distribution of Denisovan ancestry affinity across East Eurasia under varying segment-length thresholds.** Pie charts on each map show the proportion of low-, moderate-, and high-affinity introgression segments for populations across Eurasian, with maps faceted by minimum segment-length thresholds (≥25, 50, 100, 150, 200 and 250 kbp). As the length threshold increases, the proportion of high-affinity segments in EAS becomes more pronounced and shows a growing disparity in frequency compared to SEA and SAS.

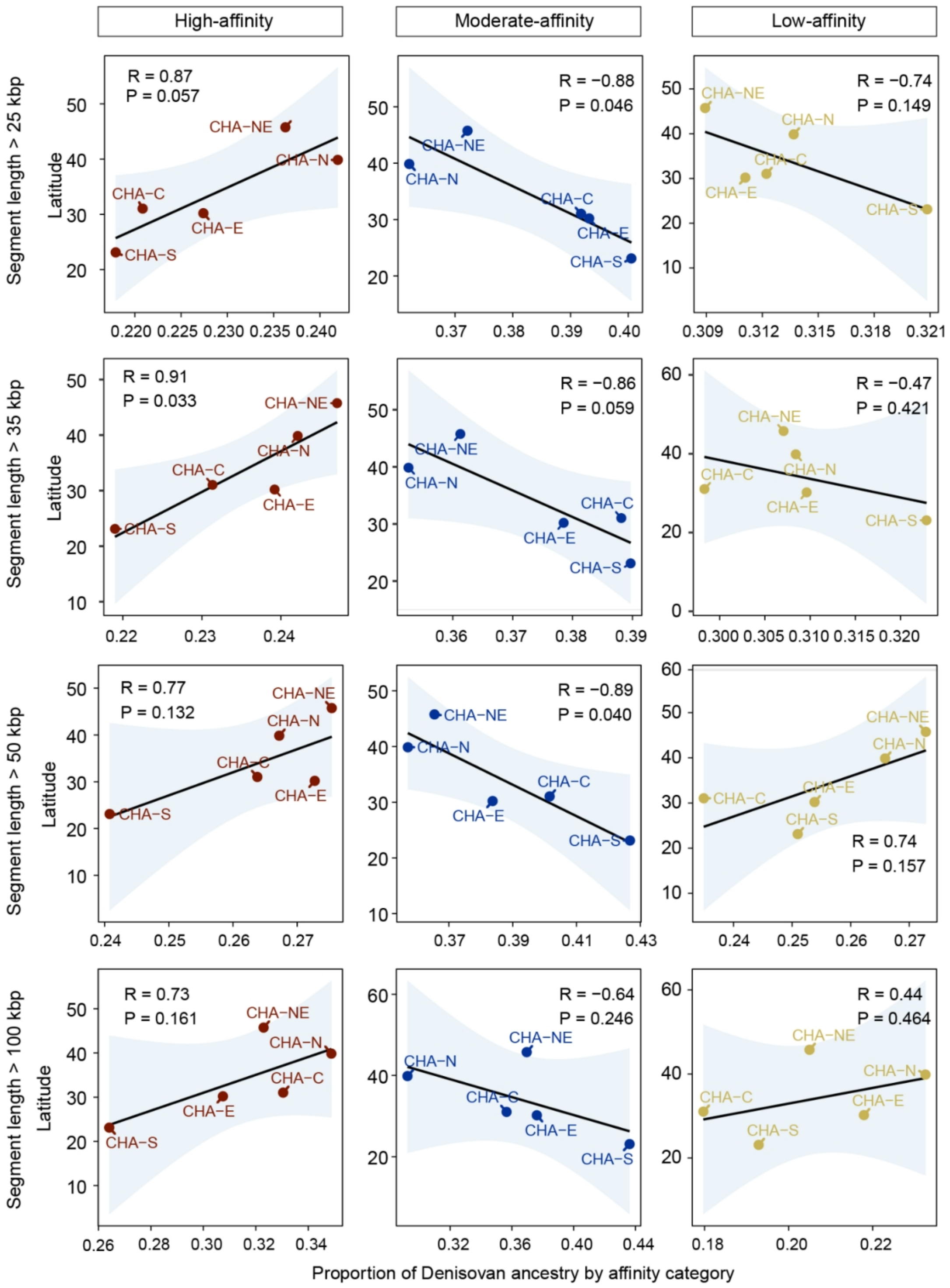


**Supplementary Fig. 48 | Latitudinal distribution of Denisovan ancestry affinity components among Han Chinese populations under varying segment-length thresholds.** The multi-panel plot illustrates the relationship between Denisovan ancestry proportions and latitudes across three affinity categories (high, moderate and low) and four segment-length thresholds (≥25, 35, 50, and 100 kbp). Each panel displays the proportion of a specific affinity component (*x*-axis) against the latitudes of Han Chinese subgroups (*y*-axis).

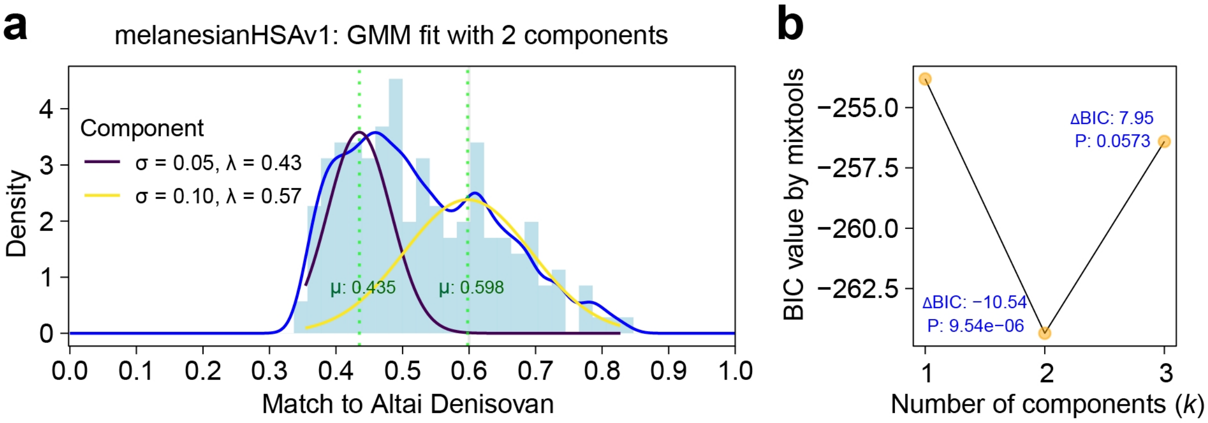


**Supplementary Fig. 49 | Match rate distributions and Gaussian mixture modeling for inferring Denisovan introgression pulses in a Melanesian genome. a**, Density distribution of match rates (blue curve) to the Altai Denisovan, overlaid with Gaussian components fitted under a two-pulses model. The two components are colored in deep violet and light goldenrod, representing distinct introgression lineages. Peaks correspond to inferred introgression events, with vertical dashed lines marking the mean match rates. **b**, Model selection based on Bayesian Information Criterion (BIC) values for one-, two-, and three-component Gaussian mixture models. ΔBIC values relative to the preceding model and *p*-values from likelihood-ratio tests between nested models are annotated.

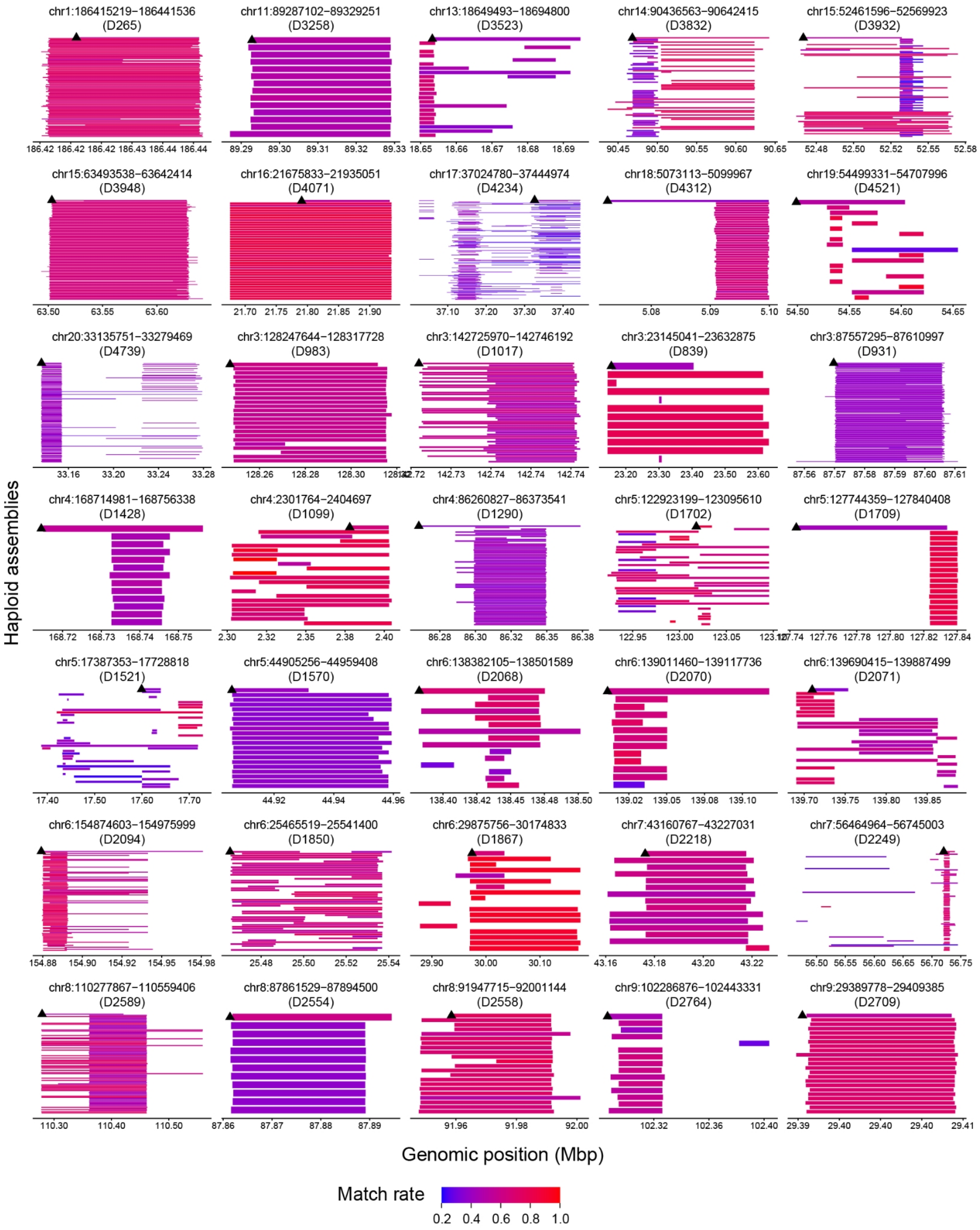


**Supplementary Fig. 50 | Shared Denisovan introgressed segments between Melanesian and other modern human haploid assemblies.** Each row represents a haploid assembly, with the Melanesian individual highlighted in the top row and marked by a black triangle. Tracks are colored by match rates to the Altai Denisovan reference genome.

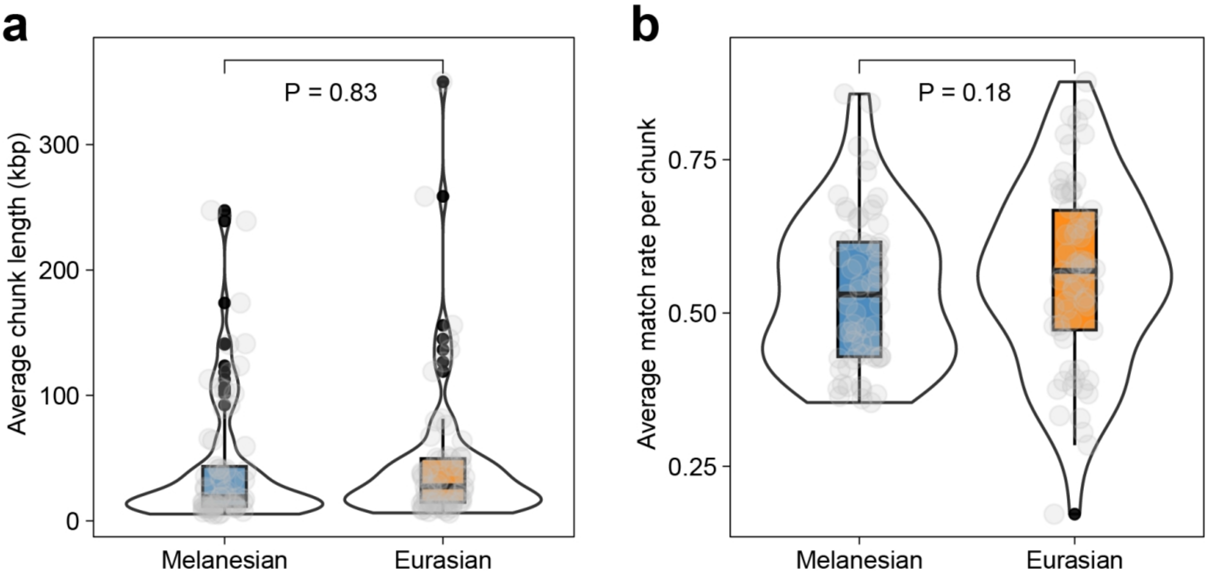


**Supplementary Fig. 51 | Comparison of shared Denisovan introgression segments between Melanesian and Eurasian haploid assemblies.** Violin plots display the distribution of (**a**) segment lengths and (**b**) match rates for Denisovan-derived segments shared between Melanesian and Eurasians. Pairwise *t*-tests show no significant difference in either segment length ($P = 0.83$) or match rate ($P = 0.18$).

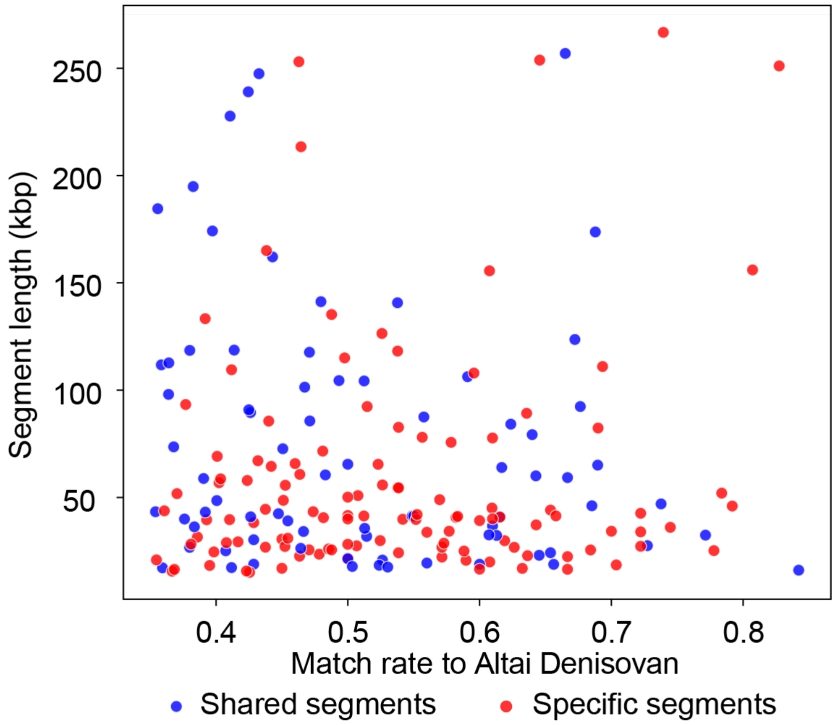


**Supplementary Fig. 52 | Distribution of introgressed segment length and match rate for shared *versus* Melanesian-specific segments.** Scatter points are colored by segment category: shared between Melanesian and other populations ($n$ = 77) and specific to the Melanesian individual ($n$ = 117).

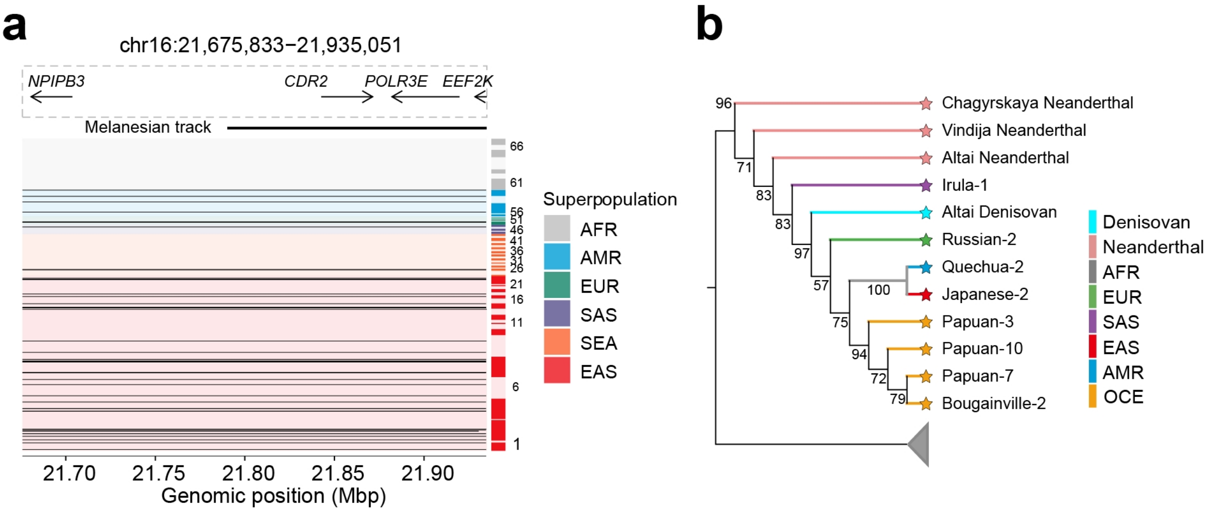


**Supplementary Fig. 53 | A shared Denisovan introgressed region D4071 at 16p12.2 between Melanesian and other modern human populations. a**, Genomic landscape of the 259.2-kbp introgressed haplotype block, encompassing multiple genes including *CDR2*, *POLR3E*, *EEF2K*. Haplotypes from haploid assembly genomes are plotted as rows, grouped and ordered by populations as defined in Supplementary Fig. 7. Denisovan-derived segments are highlighted in dark. **b**, A maximum-likelihood phylogenetic tree of the shared region (T2T-CHM13 coordinates, chr16:21,791,205-21,931,930). The phylogeny was constructed using samples from the SGDP dataset carrying this archaic segment. The topology shows that introgressed haplotypes from SAS individuals branch closest to the Denisovan reference, followed by Eurasian and Melanesian lineages, consistent with a shared ancestral origin prior to the divergence of these populations. Phylogenetically distant non-introgressed clades are collapsed for clarity.
